# Deciphering spatiotemporal patterns of rhizodeposition with a functional-structural root model: *RhizoDep*

**DOI:** 10.1101/2025.03.30.646173

**Authors:** Frédéric Rees, Tristan Gérault, Marion Gauthier, Romain Barillot, Céline Richard-Molard, Alexandra Jullien, Claire Chenu, Christophe Pradal, Bruno Andrieu

## Abstract

Rhizodeposition, i.e. the release of organic matters by roots, constitutes a significant fraction of the plant carbon (C) budget and plays a key role in soil-plant interactions. However, its spatial and temporal dynamics remain poorly understood. We developed *RhizoDep,* a new functional-structural root model that simulates 3D root growth, respiration, and rhizodeposition based on C balance and root morphology at the individual root segment level. Our model successfully reproduced the dynamics of belowground C flows observed in a previous pulse-labelling field experiment on spring wheat. Our simulations revealed that root C exudation largely dominated over mucilage secretion and cap cells sloughing in terms of C release. The spatial distribution of exudation rate along the roots was driven by the preferential unloading of sugars to support root elongation and emergence, and was modulated by the formation of apoplastic barriers. Furthermore, our results demonstrated that, for a given C allocation flow to roots, variations in root hairs or lateral root number had minimal effects on rhizodeposition, whereas changes in root tissue density had a significant impact. *RhizoDep* offers a new opportunity to explore the dynamics of C exchange at the plant-soil interface and to identify traits and environmental conditions that favor rhizodeposition.

**Highlight:** Using the new model *RhizoDep*, we simulated distinct spatial and temporal patterns of rhizodeposition along the roots of spring wheat over its complete growth cycle, and identified their main drivers.

**Graphical abstract:** 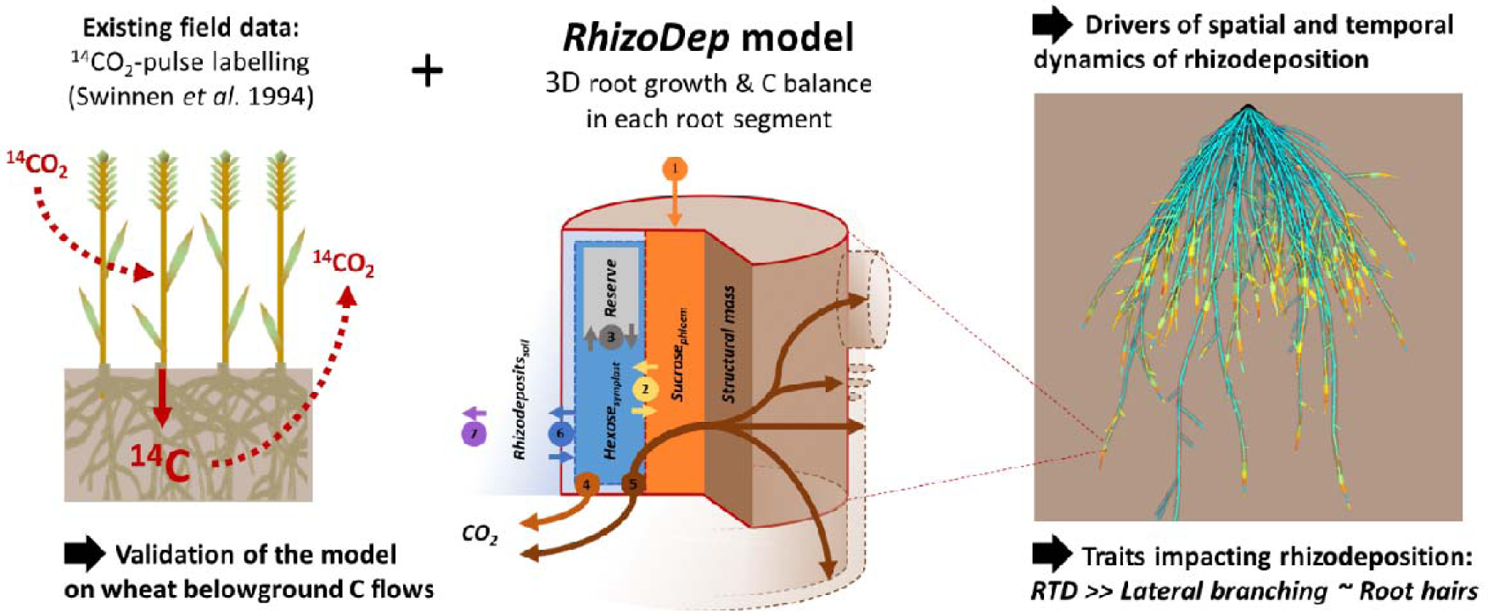

## 1 Introduction

Rhizodeposition, i.e. the release of organic carbon (C) by plant roots into the soil over their life, is a major yet poorly quantified component of the plant C cycle. This process accounts for 5-15% of net photosynthesized C and more than a third of C allocated to roots (Nguyen, 2003; Pausch and Kuzyakov, 2018). Rhizodeposits encompass various materials including water-soluble, low-molecular weight exudates, polymers (e.g. mucilage), sloughed cells, and volatile organic compounds (Jones *et al*., 2009). While root exudates have sometimes been considered as an undesired loss of C by the plant (Jones and Darrah, 1996), they have been increasingly recognized as fundamental to plant’s adaptation to its environment. For instance, plants benefit from rhizodeposition by solubilizing phosphorus via the release of carboxylates (Hinsinger, 2001) or mobilizing soil nitrogen (N) through exudate-induced acceleration of soil organic matter (SOM) mineralization (Cheng *et al*., 2014; Henneron *et al*., 2020). Additionally, mucilage secretion, and to some extent cap cell release, can favor root penetration, maintain plant water uptake, and protect the root tip against contaminants or pathogens (Morel *et al*., 1986; Uden, 2000; Hawes *et al*., 2002; Cannesan *et al*., 2011; Carminati *et al*., 2016). Certain exudates and volatiles also contribute to plant defense mechanisms (Bertin *et al*., 2003; Czarnota *et al*., 2003; Guo *et al*., 2017) and communication (Dam *et al*., 2016; Rasmann *et al*., 2017). Rhizodeposits significantly influence SOM storage, either positively by contributing to the formation of long-lasting, microbial-processed organic compounds (Sokol *et al*., 2019), or negatively through *rhizospheric priming effect* (Villarino *et al*., 2021). In this context, the spatial localization of rhizodeposits within the soil profile is critical, as the rate of SOM decomposition usually decreases with soil depth (Balesdent *et al*., 2018). Distinct rhizodeposits have contrasted effects on SOM dynamics (Mary *et al*., 1993), yet few studies have quantified the individual contribution of root exudates, mucilage, sloughed cells and volatiles to the overall release of C by roots. While mucilage secretion and cells sloughing contribute have been suggested to contribute to rhizodeposition at least 10 times less than root exudates (Nguyen, 2003; Rees *et al*., 2005), low-molecular-weight exudates were shown to constitute only 48%-86% of total rhizodeposited C in maize (Jones and Darrah, 1993).

Despite growing recognition of its importance, the spatial and temporal evolution of rhizodeposition over different scales has remained largely unexplored (Galindo-Castañeda *et al*., 2024). Root tips are widely considered the primary site of exudation (Badri and Vivanco, 2009; Jones *et al*., 2009), though some studies have suggested alternative “hotspots” such as the root hair zone, lateral root emergence sites, or even the root base (Badri and Vivanco, 2009; Holz *et al*., 2018; Voothuluru *et al*., 2018). Additionally, the temporal dynamics of rhizodeposition remain unclear. Plant labelling experiments have shown that rhizodeposited C primarily originates from recent photoassimilates, and therefore follows a diurnal pattern (Liljeroth *et al*., 1990*a*; Badri and Vivanco, 2009). However, rhizodeposition rates also evolve over the course of the growing season. In cereals, mass rhizodeposition rates (per gram of root biomass) are usually maximal at early growth stages and then decrease over time, e.g. after tillering (Keith *et al*., 1986; Remus and Augustin, 2016; Pausch and Kuzyakov, 2018). These temporal patterns have been attributed to changes in C allocation, as root exudation is often considered a passive efflux driven by the gradient of metabolites concentration between roots and the growing medium (Jones and Darrah, 1996; Henry *et al*., 2005; Personeni *et al*., 2007). However, studies indicate that variations in the exudation of a given compound do not necessarily follow variations of its concentration inside root cells (Canarini *et al*., 2016; Proctor and He, 2017). Processes such as the re-uptake of specific exudates by roots (Jones and Darrah, 1993) and root morphological changes such as the formation of hydrophobic barriers limiting radial transport may modulate these concentration-driven loss of exudates by the roots. Eventually, the spatiotemporal variability of rhizodeposition arises from complex interactions between genotype, environment, and growth stage, contributing to the high variability observed across studies.

Plant models are valuable tools for integrating these interactions and predicting rhizodeposition dynamics. However, most crop models do not explicitly represent rhizodeposition, often treating it solely as root decay, as in STICS (Beaudoin *et al*., 2023). Given the interplay between root architecture and plant metabolism in rhizodeposition, Functional Structural Plant Models (FSPM) offer a promising approach. Despite their potential, only a few FSPM have incorporated rhizodeposition processes until now. Some models consider rhizodeposition as a fixed fraction of root-allocated C (Barillot *et al*., 2016), while others do not distinguish it from root decay (Louarn and Faverjon, 2018). Landl et al. (2021) extended the *CPlantBox* model that explicitly includes a dynamic root architecture, by considering a term of citrate exudation and mucilage secretion along roots. This represents one of the first attempts to simulate rhizodeposition on a 3D root system. However, their model used constant rhizodeposition rates independent of metabolic activity or C allocation. Two decades earlier, Jones and Darrah (1996) proposed a conceptual model linking sugar exudation to sugars concentration and root cell membrane permeability, later refined to account for a decrease in the permeability with the distance from root tip (Personeni *et al*., 2007). However, these approaches have never been tested at the whole root system scale. Hence, to the best of our knowledge, no existing models explicitly simulate rhizodeposition as a function of root architecture and plant C balance.

This study aims to explore spatial and temporal patterns of rhizodeposition *in silico* using *RhizoDep*, a new mechanistic functional structural root model. We hypothesize that local C balance along roots represents the main factor explaining rhizodeposition patterns. First, we introduce the principles of *RhizoDep* and then validate its simulation outputs against belowground C flows from a previously published ^14^C-labelling field experiment on spring wheat. We then analyze the spatial variability of distinct rhizodeposition processes along wheat roots and finally evaluate the impact of key root traits on rhizodeposition.

## 2 Model description

### 2.1 Model overview

*RhizoDep* is a process-based FSPM that simulates how roots grow, respire, and emit rhizodeposits into the soil, in interaction with a dynamic 3D root architecture and the distribution of available C (Fig. 1). The model operates at fine spatial and temporal scales, typically simulating root segments of 1-mm with an hourly time step. *RhizoDep* incorporates simplified physiological processes that account for major C transfers within the roots and between roots, soil and atmosphere. Aside from soil temperature, which can affect rates in various ways, the main input variable of the model is the net C transferred from the aerial parts of the plant to the root system. This input is represented as sucrose (C_12_H_22_O_11_) transported through the phloem vessels. A portion of this sucrose is then unloaded and converted into hexose (C_6_H_12_O_6_), which serves as the substrate driving key metabolic processes. These include: i) the production of root structural mass, associated with the elongation of roots or root hairs and secondary growth, ii) root maintenance, iii) sugar immobilization in or remobilization from a reserve pool, and iv) rhizodeposition. No priority is assigned to any of these processes in terms of hexose consumption. Additionally, a simple representation of the fate of rhizodeposits outside the roots has been implemented, though such processes are expected to be managed by a dedicated soil model in the future. *RhizoDep* focuses on C dynamics within the root-soil system and does not simulate the aerial parts of the plant, C allocation from shoots to roots, or the dynamics of water and mineral nutrients. Similarly, C costs associated with symbiotic microorganisms (e.g. *Rhizobium*, mycorrhizae) or parasitic organisms (e.g. root-knot nematodes, broomrape) have been excluded but may be incorporated in future versions.

**Fig. 1:**
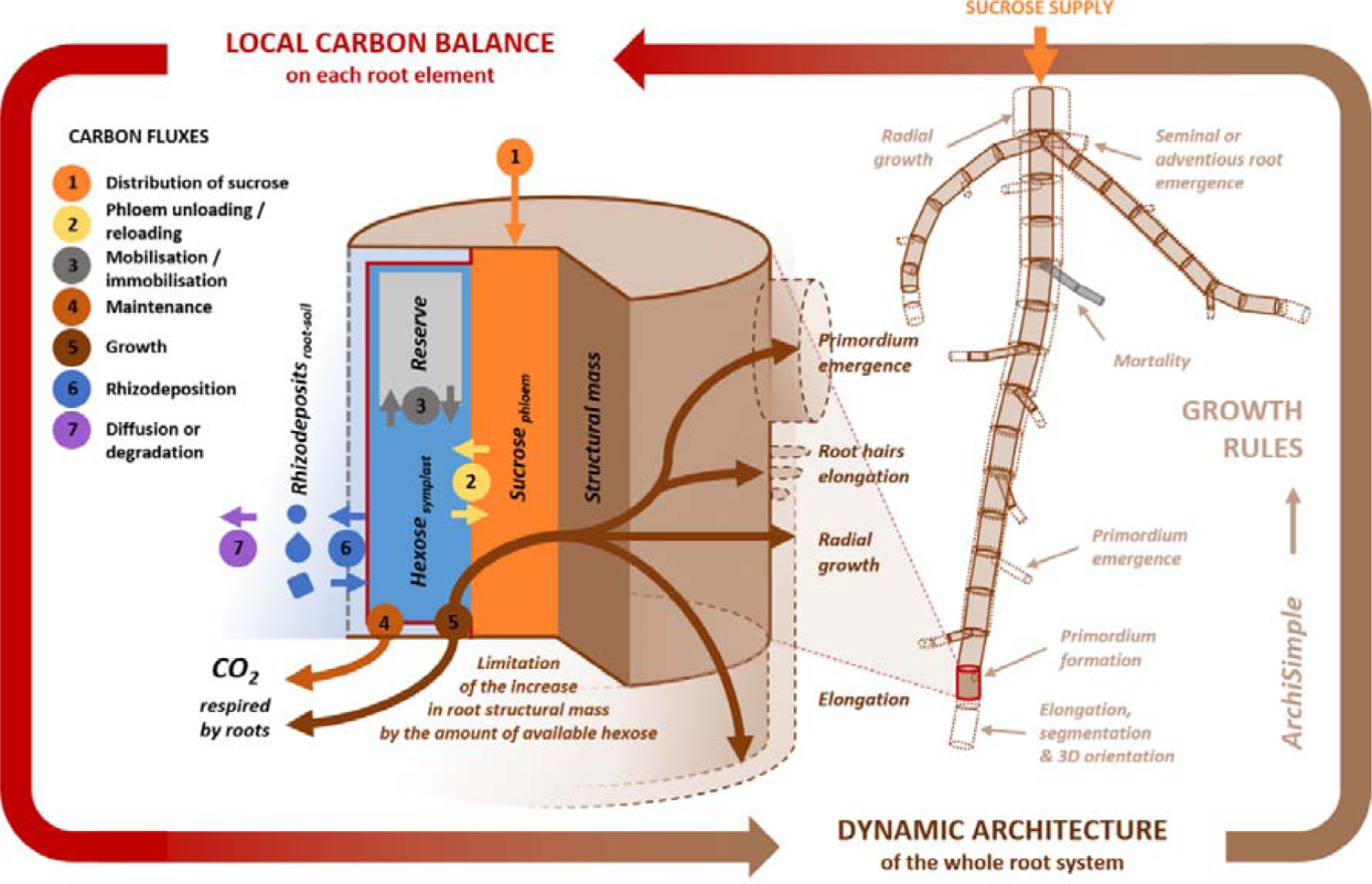
Overview of the *RhizoDep* model. Photoassimilated C enters the root system as sucrose (model input), which is then distributed among each root element (1) and can be converted into hexose (2). Hexose may be stored in a reserve pool (3), used for maintenance (4), or consumed for growth, contributing to structural mass production - either through longitudinal growth (for an apex) or radial growth (for a segment), or through the emergence of new lateral roots or root hairs (5). Once exuded outside the root, hexose can be reabsorbed by the root (6) or lost from the root-soil interface due to microbial degradation, sorption, or diffusion (7). Similarly, other rhizodeposits (e.g., mucilage, sloughed cells) can accumulate at the root-soil interface or be lost. The local C balance within each root element ultimately constrains potential root architecture development, as defined by growth rules adapted from the *ArchiSimple* model (Pagès et al. 2014).

Below, we provide a brief summary of the main features of the model. Additional details, including equations and justifications, are available in the Supporting Information file (SI-1). A list of the key variables and parameters of the model can be found in Supporting Table ST.

#### 2.1.1 Representation of the root-soil system

In *RhizoDep,* the root system is discretized into root elements represented as cylindrical segments, each defined by its length and radius. Any given root axis consists of a series of segments of constant length *l_segment_*, terminating in an apical cylinder. Segments can grow radially but not longitudinally, whereas apices can elongate but cannot expand radially. When an apex exceed *l_segment_* in length, it is segmented to form at least one new segment and one Each root element is characterized by its dimensions, 3D coordinates, structural mass, and individual pools of sucrose, hexose, and reserve C (Fig. 1). Structural C is determined based on the element’s volume, using a constant root tissue density and C content; it can only be modified by longitudinal or radial growth. As sucrose is the primary compound transported through the phloem (Lohaus, 2022), it represents in the model the only form of C that can be transported between root elements. Hexose, on the other hand, includes all mobile C available within the epidermal, cortical, and non-vascular stele’s cells. Reserve C encompasses all starch, fructan, and vacuolar sugars that are not immediately available to support root growth, maintenance, or rhizodeposition. The concentrations of sucrose, hexose, and reserve are assumed homogeneous within each root element and are expressed relative to structural mass (moles per gram). *RhizoDep* does not differentiate between root tissues (e.g. epidermis, cortex, stele) except when calculating the effective exchange surface between the root and the surrounding soil solution. Root hairs are treated as a specific extension of this exchange surface and as a contributor to the structural mass of the root element. The “soil” compartment in *RhizoDep* is limited to the root-soil interface associated with each root element, characterized by its exchange surface and the concentration of hexose, mucilage, and sloughed cells. The use of an intermediary root-soil interface, rather than an explicit soil volume, has been introduced to facilitate coupling *RhizoDep* with external model capable of describing rhizodeposits transformation in the soil.

#### 2.1.2 Distribution of sucrose

The C allocated as sucrose from the aerial parts to the root system is distributed across root elements under the assumption of no resistance to longitudinal transport (see details in SI 1.1). As a result, sucrose concentration remains homogeneous throughout the root system, consistent with the theory of rapid pressure-concentration waves propagating along the phloem (Thompson and Holbrook, 2004). Within each root element, sucrose is converted into hexose (SI 1.2) based on local concentration gradient between sucrose and hexose (Eq. S1) and the rate of hexose consumption for growth in the element (Eq. S2). Conversely, hexose can be reconverted into sucrose depending on the concentration of mobile hexose (Eq. S3), mimicking the phloem reloading occurring in non-growing parts of the root system.

#### 2.1.3 Root growth

The potential growth of each root element is defined based on an adaptation of the growth rules implemented in *ArchiSimple* (Pagès *et al*., 2014). Further details are provided in section SI 1.3. Briefly, lateral root primordia form at the root tip of the “mother” root, maintaining a constant distance between primordia along the same axis. In *ArchiSimple,* most root properties derive from the initial root apical diameter. The diameter of a new lateral root is randomly drawn from a Gaussian distribution centered on a mean diameter proportional to that of the mother root, and the new root lateral emerges following a predetermined dormancy period. The potential elongation rate of a root apex and its lifespan are both proportional to its diameter, while the total elongation duration of a root is proportional to the square of its diameter. Finally, root abscission occurs when the apex of the root, along with the apices of all lateral roots on the same axis, reach the end of their lifespan.

While most of the rules of *ArchiSimple* remain unchanged in *RhizoDep*, a key difference lies in how actual root growth is regulated by available C. In *ArchiSimple*, root elongation is constrained by a single “satisfaction coefficient”, which represents the ratio between biomass input from the shoots and the total cumulative root biomass demand for elongation across all axes. In *RhizoDep*, growth is instead limited by the amount of locally available hexose that can be converted into new structural material. Actual growth rates are further modulated within each root element by the local hexose concentration (Eq. S6). Notably, the mobile hexose available for elongation corresponds to the hexose within the root elongation zone, whose length depends on the apical radius (Eq. S7) and may span multiple root elements (Fig. S1). Additionally, the conversion of hexose into structural mass generates growth respiration (Thornley and Cannell, 2000), consuming an additional fraction of the available hexose (Eq. S12).

#### 2.1.4 Root hairs dynamics

In *RhizoDep*, root hairs on each element are defined by their density (number of hairs per unit root length), as well as their average length, diameter, and age (see SI 1.4). Root hairs are assumed to emerge just above the root elongation zone (Guichard *et al*., 2019) and to elongate until they reach their maximal length *l_hair_max_* (meter). The elongation of living root hairs follows a potential growth rate, regulated by the concentration of hexose available within the supporting root element (Eq. S13). Once emerged, root hairs remain alive for a predefined duration. Given the lack of evidence suggesting that dead root hairs are sloughed off over time (Fusseder, 1987; Nguyen, 2003), we assume that dead root hairs remain attached to the supporting root element - and consequently to the root system - until the root element itself dies.

#### 2.1.5 Root maintenance

In *RhizoDep*, root maintenance is represented by a single function that encompasses all growth-independent processes requiring energy to sustain the functioning of a given root element (see SI 1.5). Maintenance costs are assumed to be proportional to the dry structural mass and increase with the concentration of mobile hexose within the root element (Eq. S14), as suggested in previous modelling studies (Thornley and Cannell, 2000; Barillot *et al*., 2016).

#### 2.1.6 Exchange with reserve

*RhizoDep* simulates a C reserve pool for each element, defined by its concentration in hexose-equivalent (see SI 1.6). Depending on the equilibrium between the reserve concentration and the concentration of mobile hexose, a net immobilization or remobilization of hexose may occur within the root element (Eq. S16-17). The reserve pool is also constrained by a maximal achievable concentration.

#### 2.1.7 Dynamics of rhizodeposits

In *Rhizodep,* rhizodeposition processes are represented as the exudation of sugars, the secretion of mucilage, and the release of root cap cells, while the emission of VOC has been neglected so far. Root exudation is modeled as the transfer of the two mobile C compounds represented in *RhizoDep* - hexose and sucrose - from the root into the soil solution (see SI 1.7). The net efflux of hexose from non-vascular cells and sucrose from phloem (Eq. 1 and 2) is described as the balance between gross exudation, driven by a diffusion gradient of concentration across the root plasma membrane, and active influx from the root-soil interface, modeled using a Michaelis-Menten formalism (Jones and Darrah, 1993; Farrar *et al*., 2003; Personeni *et al*., 2007). The net losses in the amounts of hexose (*Hex,* moles of hexose) and sucrose (*Suc*, moles of sucrose) within a given root element *i* due to exudation are therefore expressed as:

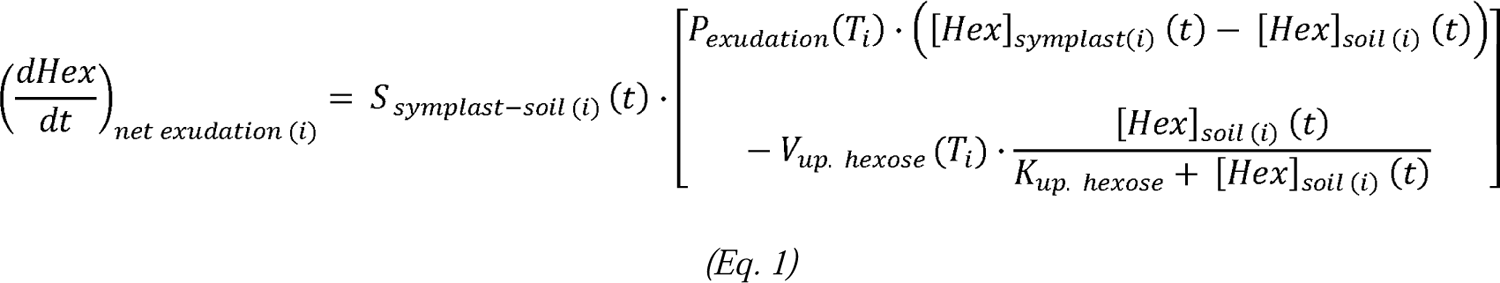

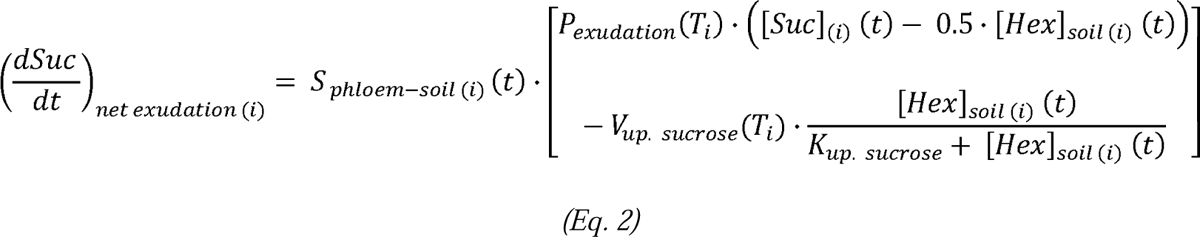

where *[Hex]_symplast (i)_*, *[Hex]_soil (i)_* and *[Suc] _(i)_* represent the concentrations of root mobile hexose, hexose at the root-soil interface and sucrose within the root element *i* (mol gDW^-1^), *T_i_* is the soil temperature perceived by the root element *i* (K), and *P_exudation_* (gDW s^-1^ m^-2^), *V_up. hexose_* and *V_up. hexose_* (mol s^-1^ m^-2^), *K_up_hexose_* and *K_up. sucrose_* (mol gDW^-1^) are fixed or temperature-dependent parameters. The exchange surface for hexose (*S _symplast-soil (i)_*) and sucrose (*S _phloem-soil (i)_*) depends on the morphology of the root element, particularly the presence of barriers to radial solute transport. When new root elements are formed, the apoplast (i.e. the intercellular space outside root plasma membranes) allows the external soil solution to reach the center of the root (Fig. S2), enabling sugar exchange with all root cells, including the phloem vessels. As the endodermis develops between the stele and cortex, direct access of the external soil solution to the root cells within the stele is lost, thereby restricting the exchange surface to epidermal and cortical cells only (Kamula *et al*., 1994). Similarly, if an exodermis forms beneath the epidermis, the exchange surface is further reduced, eventually being limited to the membrane of epidermal cells and living root hairs. Events such as the emergence of lateral roots can temporarily disrupt these transport barriers, increasing the exchange surface between root cells and the surrounding soil solution.

Mucilage secretion corresponds to a unidirectional flux from the root epidermis into the soil. The potential secretion rate is maximal at root tip and is assumed to decrease non-linearly with distance from the tip until becoming negligible (Paull and Jones, 1975) (see S1.8, Eq. S30). The actual secretion rate depends on the external root surface area, and is constrained by both the hexose concentration in peripheral root cells and the surfacic concentration of mucilage at the root-soil interface (Eq. S30-S31). Consequently, mucilage secretion is inhibited when mucilage accumulates at the root-soil interface, as observed during dry conditions (Morré *et al*., 1967).

The release of root cap cells also follows an unidirectional flux of matter. The potential release rate decreases linearly with distance from the root tip and drops to zero above the elongation zone (see SI 1.9). The actual rate of cell release depends on the external surface area of the root element but is independent of mobile hexose availability (Curlango-Rivera *et al*., 2010). However, it is limited by the surfacic concentration of root cap cells accumulated at the external surface, eventually becoming inhibited beyond a certain threshold (Brigham *et al*., 1998) (Eq. S32-S33). Notably, this description of cell release excludes the contribution of root hairs sloughing, which was not incorporated into *RhizoDep* due to a lack of evidence that dead root hairs are shed over time (Fusseder, 1987; Nguyen, 2003).

Once released into the soil, each type of rhizodeposit either accumulates at the root-soil interface or disappears (SI 1.10). In the absence of a dedicated soil model able to represent the degradation of rhizodeposits, a Michaelis-Menten kinetics approach was implemented to mimick microbial degradation for each rhizodeposit (Eq. S34-36).

#### 2.1.8 Carbon balance

At the end of a time step, a local C balance is established by accounting for all relevant processes. This balance determines the changes in the amounts of sucrose (*Suc*, in moles of sucrose), hexose (*Hex*, in moles of hexose), mucilage (*Muc,* in moles of equivalent-hexose) and sloughed cells (*Cells,* in moles of equivalent-hexose) within the various compartments of each root element *i,* which are then used to update the system for the next time step: (see details in SI 1.11):

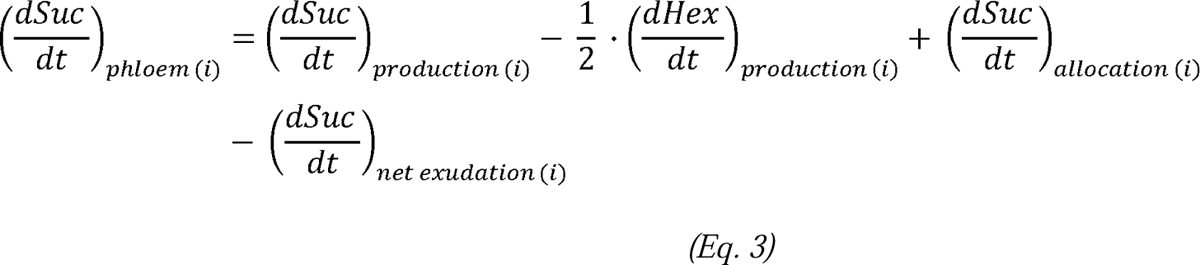

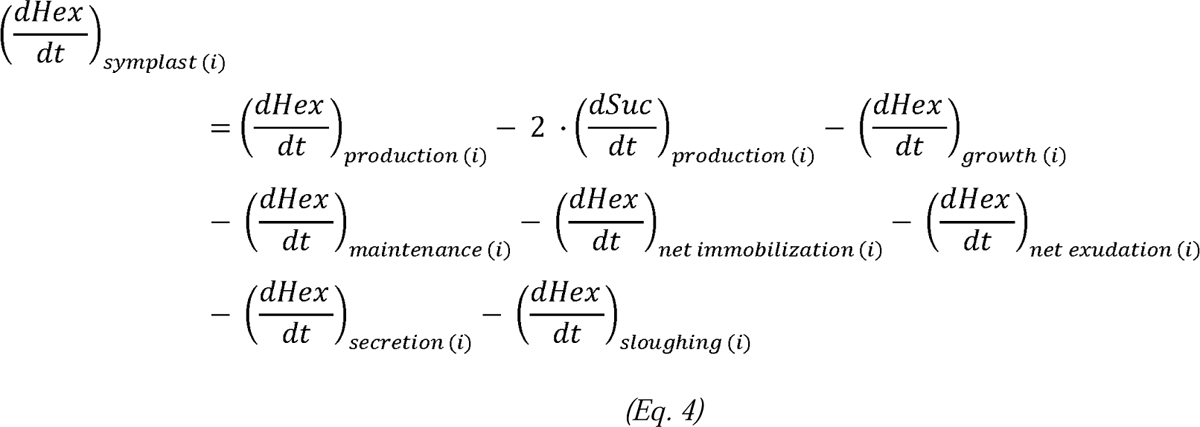

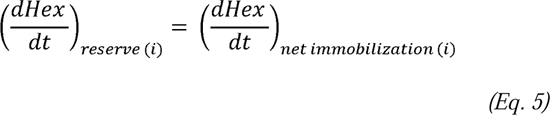

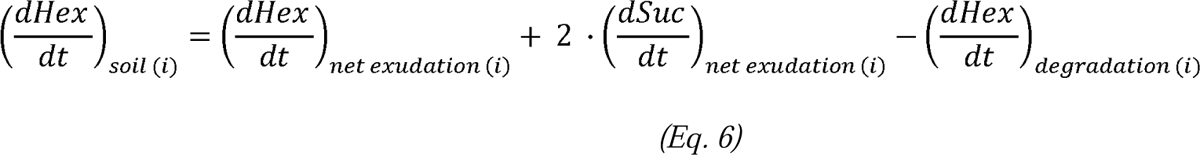

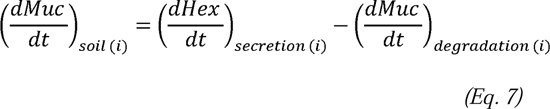

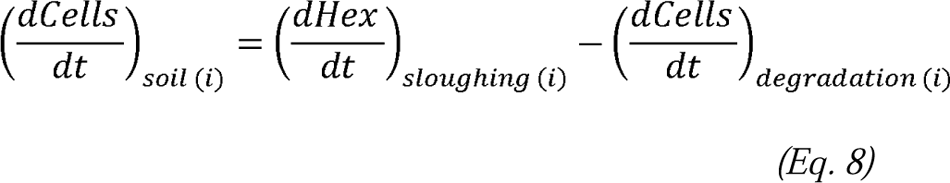

If the C balance results in a negative amount in any compartment of root element *i*, the amount is set to zero, and the deficit is accounted for in the next time step.

### 2.2 Model organization, implementation and resolution

*RhizoDep* is implemented in Python 3 (version > 3.6) (Van Rossum and Drake, 2009) and integrated within the *OpenAlea* platform (Pradal *et al*., 2008). It utilizes the *Multiscale Tree Graph* structure (Godin and Caraglio, 1998; Pradal and Godin, 2020), which allows for the storage and retrieval of all topological, geometrical, and functional properties of each root element at every time step. The geometry of the root system and the spatial distribution of variables along the roots can be visualized in a 3D graph using *PlantGL* (Pradal *et al*., 2009) or *Pyvista* (Sullivan and Kaszynski, 2019). The code is open-source, licensed under CeCILL-C, and available as an *OpenAlea* package on *GitHub* (https://github.com/openalea/rhizodep).

*RhizoDep* is structured in several modules, each corresponding to a specific process (Fig. S3). The model operates using a discrete-time numerical integration scheme with a time step of 1 hour. At each time step, starting at *t* and ending at *t* + Δ*t*, all modules compute the new state variables defined at *t* + Δ*t* based on the state variables from *t* (i.e., the end of the previous time step). This approach ensures that all processes operate simultaneously without anyone process being prioritized in resource consumption and allocation.

## 3 Simulation plan

### 3.1 Validating *RhizoDep* for spring wheat

To validate *RhizoDep*, we used the data from the field study conducted by Swinnen *et al*. (1994a). These authors monitored the entire growth cycle of spring wheat (*Triticum aestivum* var. Minaret) from March to August 1989 in Netherlands. Using repeated ^14^CO_2_ pulse labelling and belowground respiration measurements over several days, they estimated the temporal distribution of photoassimilated C among root biomass production, root respiration, and rhizodeposition. They also derived belowground C fluxes based on the evolution of shoot biomass and shoot:root biomass ratio, fitted with empirical functions (see details in SI 4). To extend their dataset, we supplemented their calculations to estimate C flows at both germination and harvest stages (see SI 5). The estimated C flow allocated to roots over time served as the sucrose input in *RhizoDep,* allowing us to simulate the experiment *in silico.* Soil temperature variations over the simulation period were derived from temperature measurements recorded one year later at the same site (SI 4.6).

Model parametrization was conducted independently of the Swinnen *et al*. (1994a) dataset. A complete list of model parameters, their estimated values, and sources is provided in Supporting Tables ST. Growth parameters for *RhizoDep* were estimated based on our own unpublished pot and rhizobox experiments (see SI 6.1), while other parameters were sourced from literature data for wheat and, in some cases, other plant species (see SI 6.2). For approximately 25% of parameters, no direct data were available, so values were inferred based on similar parameters already defined in the model. In only two instances did the initial parameter estimates produce unrealistic results, requiring manual adjustment by trial and error until plausible sucrose unloading patterns in growing roots were achieved (see SI 6.2).

### 3.2 Testing the influence of specific root traits

To determine which root traits most strongly influence rhizodeposition and its contribution to root C balance, we ran the same baseline simulation for spring wheat (**Sc1**) and compared it against modified scenarios related to three root traits:

- **Root hair density**: Root hairs were either repressed (**Sc2**) or their density was doubled along the roots (**Sc3**).
- **Lateral root formation**: The average distance between lateral root primordia was either doubled (**Sc4**), reducing lateral root number, or halved (**Sc5**), increasing lateral root number.
- **Root tissue density**: Tissue density was either halved (**Sc6**) or increased by 50% (**Sc7**) to capture a plausible range of variation (Pagès et al., 2014).

We hypothesized that scenarios reducing root exchange surface (Sc2 and Sc4) would lead to lower rhizodeposition, whereas increasing the number of active exudation sites (e.g., root tips in Sc5) or lowering C consumption for structural mass production (Sc6) would increase rhizodeposition.

## 4 Simulation results

### 4.1 Simulation of belowground C flows during spring wheat growth

We simulated the temporal evolution of belowground C flows throughout the entire growth cycle of spring wheat (scenario Sc1). As shown in Fig. 2a, the cumulative flows of root biomass production, respiration, and rhizodeposition predicted by *RhizoDep* closely matched those calculated by Swinnen *et al*. (1994b). The final distribution of belowground C among biomass production, root respiration, and rhizodeposition was 0.38, 0.33 and 0.21 gC plant^-1^, respectively - differing by only 7%, 10% and 3% from the estimation of Swinnen *et al*. (1994b). The daily production of root structural mass (gC per day per plant) increased rapidly after mid-April, peaking at the end of May. In contrast, the flows associated with root respiration and rhizodeposition peaked one to two weeks later (Fig 2b), a period corresponding to the onset of stem elongation, as reported by Swinnen *et al*. (1994a). The simulated release of C through dead roots became significant in June and surpassed rhizodeposited C from early July onward (Fig 2b).

**Fig. 2:**
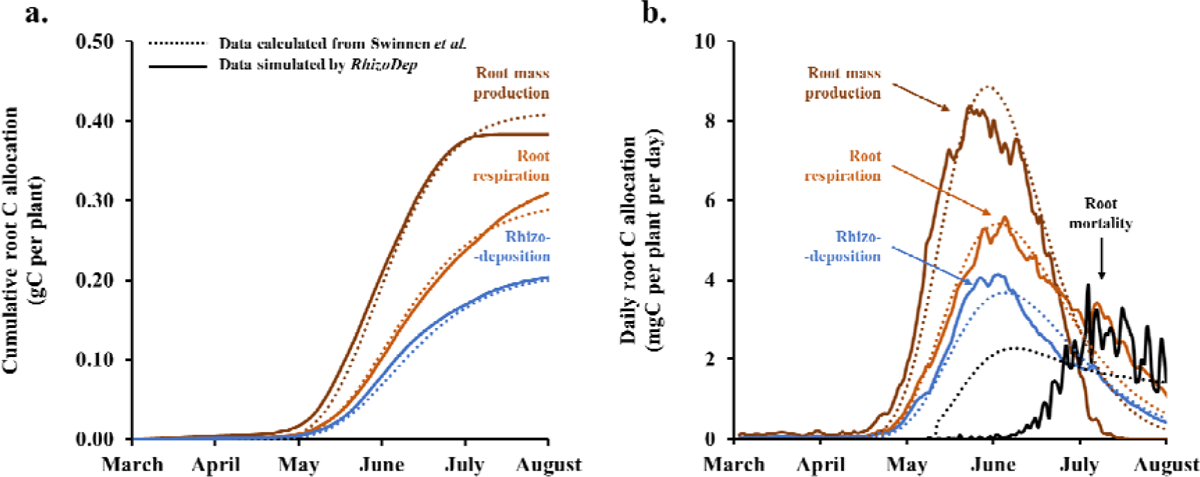
Comparison of belowground C flows of spring wheat, as calculated from the data of Swinnen *et al*. (1994b) (dashed lines) and simulated by *RhizoDep* (continuous lines) in scenario Sc1, using the same C input to the roots. Flows are shown either cumulatively (a) or on a daily basis (b) over the entire growth period. Note that in Swinnen *et al.,* root mass production corresponds to total biomass, whereas in *RhizoDep*, it represents structural mass only.

When integrating rhizodeposited C and dead root C over the top 40 cm of soil and the six-month growth period, the total root-derived C input to the soil was highest in the top 5-cm layer and decreased with depth (Fig. 3). The proportion of rhizodeposits in the total root-derived C inputs ranged between 29% and 40%, peaking in the top 5-cm layer (Fig. 3).

**Fig. 3:**
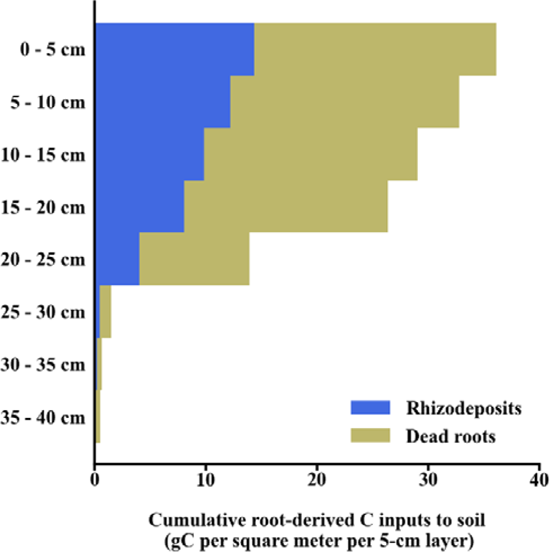
Distribution of the cumulative rhizodeposited C and dead root C over the first 40 cm of soil after 6 months, as simulated by *RhizoDep* in scenario Sc1 based on the work of Swinnen *et al*. (1994b). The simulation ouputs per plant were converted per surface of soil assuming a density of 240 plants m^-2^.

As expected, root exudation was the dominant rhizodeposition flow, accounting for 98.4 % of the rhizodeposited C over the entire growth period. In contrast, the total release of root caps cells and the secretion of mucilage contributed to only 1.3 % and 0.3 % of the rhizodeposited C, respectively (Table 1).

**Table 1:**
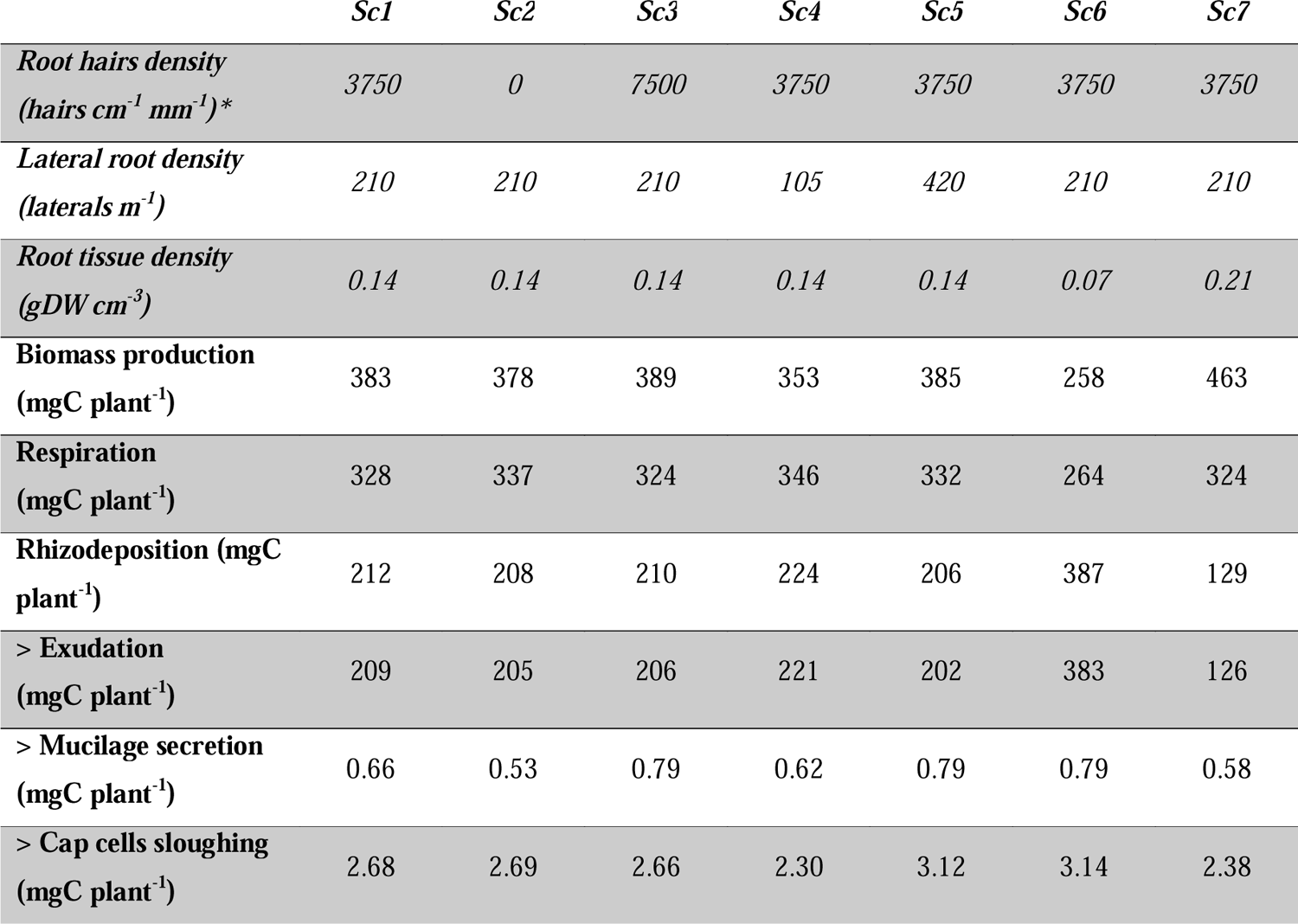
Comparison of cumulative belowground C allocation in spring wheat. 402 **over 6 months of growth**, as simulated by *RhizoDep* across scenarios *Sc1-Sc7. Sc1: reference scenario. Sc2: no root hairs. Sc3: twice more root hairs. Sc4: 50% less lateral roots. Sc5: twice more lateral roots. Sc6: 50% lower RTD. Sc7: 50% higher RTD. * Root hairs density is expressed as the number of hairs per centimeter of root for a root with a 1-mm diameter*.

### 4.2 Spatial distribution of rhizodeposition along the roots

A large spatial variability of rhizodeposition was simulated across the root system. As shown in Fig. 4 and Video 1, the lineic rate of rhizodeposition (gC per day per cm) peaked near growing root tips. To explore this pattern in detail, we examined the first seminal root and analyzed the distribution of the three rhizodeposition processes alongside potential explanatory variables every 30 days (Fig. 5 and Fig. S6-S13). As long as the seminal root continued elongating (t ≤ 104 days), the lineic rhizodeposition rate was highest at 0.5-1 cm and around 3 cm from root tip. The rate plateaued around 6 cm from the tip (Fig. S6), at a level 20 to 30 times lower than at 1 cm, and remained stable further along the root (data not shown). Once elongation ceased, the rhizodeposition rate became uniform along the seminal root axis (Fig. 4 and Fig. S6). The spatial distribution of exudation closely matched that of total rhizodeposition (Fig. 5a, Fig. S7). Exudation was 15% lower than total rhizodeposition at the root tip but differed by less than 1% beyond 10 mm. In comparison, root cap cell release was highest at the tip and decreased linearly over the first cm until disappearing (Fig. 5a, Fig. S8). Mucilage secretion also peaked at the tip, decreasing sharply over the first 2 cm and more gradually along the rest of the root (Fig. 5a, Fig. S9). Exudation profile appeared to be more strongly linked to root mobile hexose concentrations (Fig. S10) than to root exchange surface (Fig. S13). When pooling data from all five time points, net rhizodeposition correlated strongly with root mobile hexose concentration along the first seminal root (r=0.94), while its correlation with root exchange surface was weaker (r=0.79) (Fig. S14). However, when analyzing each time point separately and focusing on mature root segments (> 3 cm from the tip), rhizodeposition varied closely with root exchange surface (R^2^ > 0.98), but was negatively correlated with root mobile hexose concentration (Fig. S15-S16).

**Fig. 4:**
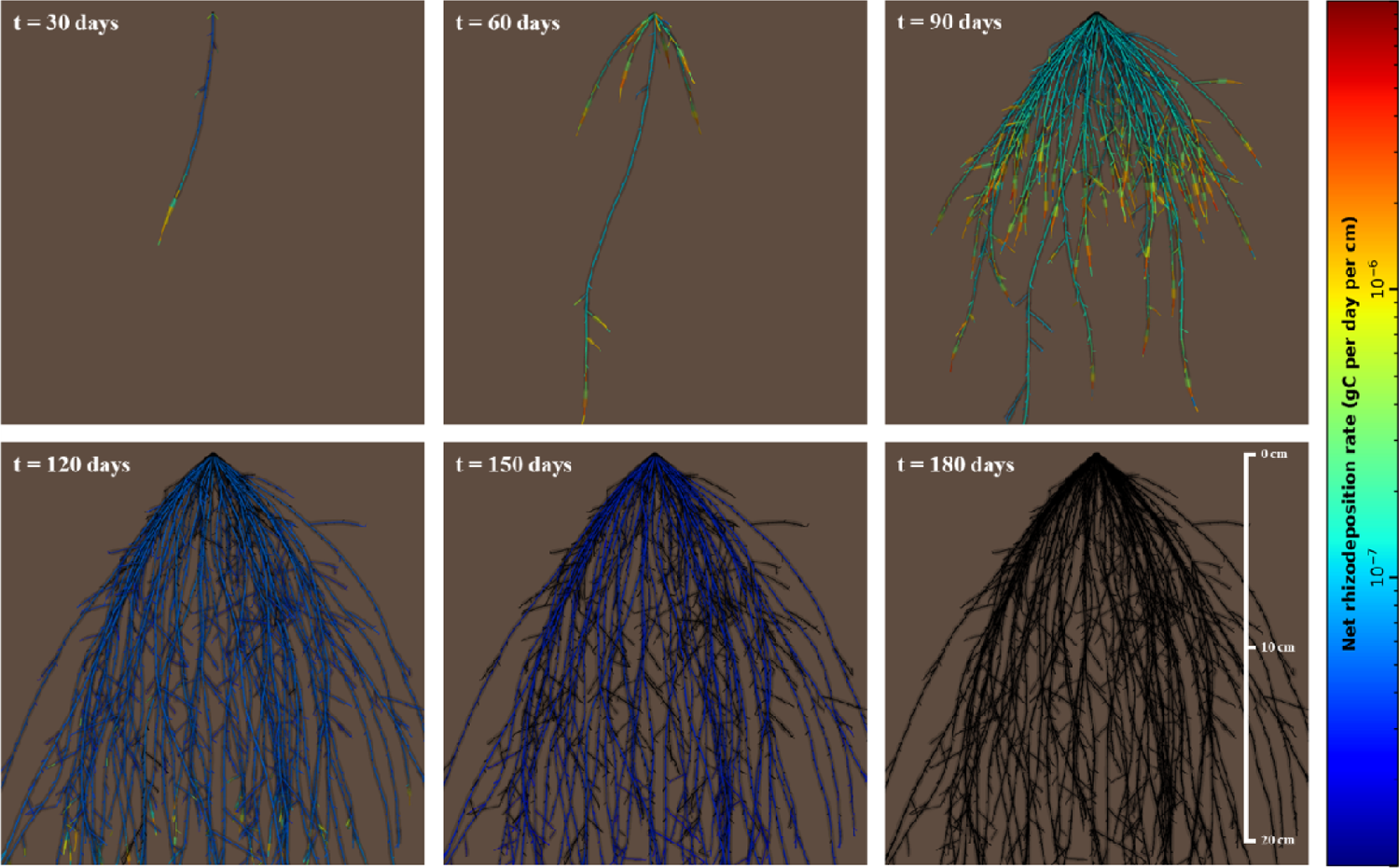
Simulated dynamic growth and rhizodeposition of the spring wheat root system from March to August 1989, based on *RhizoDep* scenario Sc1 and the data from Swinnen et al. (1994b). Colors indicate the net rhizodeposition flow (in gC per cm per day) along each root over time. The volume of living root hairs is represented as a semi-transparent colored cylinder, while dead roots and dead root hairs are shown in black.

**Fig. 5:**
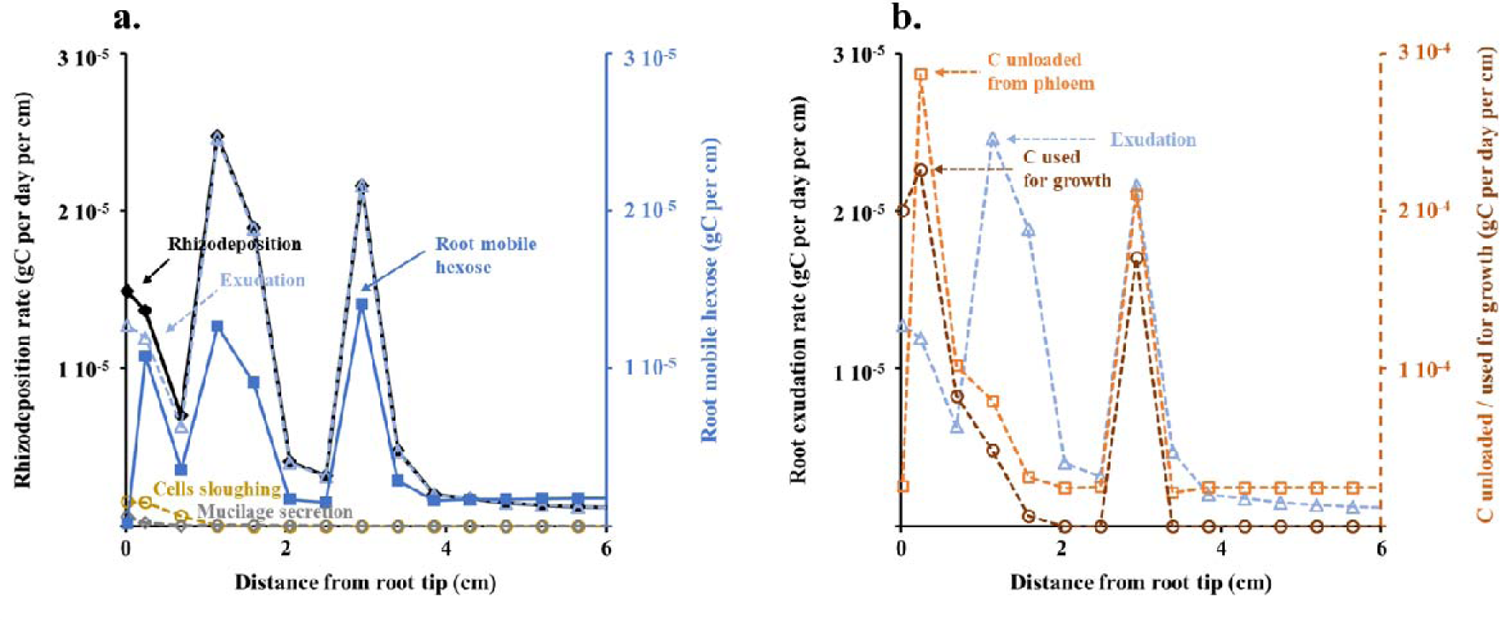
Distribution of rhizodeposition rates and mobile hexose concentration (a), as well as exudation rate, sucrose unloading rate, and C consumption rate for sustaining growth (b) along the first seminal root of spring wheat, 60 days after germination. Simulation were performed using *RhizoDep* in scenario Sc1. Values represent averages over a 7-hour period, during which root elongation was less than 2 mm.

To investigate further the contribution of different parts of the root system, we aggregated segments according to root order and/or to their position within the root axis. Over the 6 months of spring wheat growth, seminal roots and adventitious roots contributed more to rhizodeposition than lateral roots (Fig. S17). The apical parts of seminal and adventitious roots, defined here as the first 4 cm of each axis, contributed more to rhizodeposition than the basal parts above 4 cm from root tip, even though their contribution to total root length was much lower (Fig. S17). However, the contribution of roots’ basal parts to rhizodeposition gradually increased over time until becoming dominant after 4 months (Fig. S18).

### 4.3 Influence of root hairs, lateral roots and root tissue density on wheat’s rhizodeposition

To identify the root traits most influential on rhizodeposition and its contribution to the root C balance, we simulated spring wheat growth and conducted a sensitivity analysis focusing on three root traits. Surprisingly, root hairs had minimal impact on overall C flows: their removal (scenario Sc2) or a twofold increase in their density (Sc3) altered total root structural C production and rhizodeposition by no more than 2% (Table 1). Halving lateral root density (Sc4) led to an 8% decrease in root mass production, while root respiration and rhizodeposition increased by 6% each. Doubling lateral root density (Sc5) resulted in only minor changes, e.g. a 3% reduction in root exudation (Table 1). The most pronounced effects were observed when modifying root tissue density: halving it (Sc6) nearly doubled the allocation of belowground C to rhizodeposition (0.39 gC plant^-1^), while root biomass production and respiration declined by 33% and 19%, respectively (Table 1). Conversely, increasing root tissue density by 50% (Sc7) promoted root biomass production (+21%) at the expense of rhizodeposition, which decreased by 39%.

## 5 Discussion

We developed a mechanistic root functional-structural model, *RhizoDep,* to elucidate the drivers of rhizodeposition dynamics over time and space. The model successfully reproduced belowground C flows similar to those observed in a previous field experiment with spring wheat. Our simulations demonstrated that root exudation was the primary contributor to rhizodeposited C, while mucilage secretion or cells sloughing played only minor roles. When exploring the spatial distribution of rhizodeposition along roots, we found that rhizodeposition peaked at two distinct locations near the growing root tip. This pattern was primarily driven by the preferential unloading of sugars to sustain growth near the root tip and further modulated by changes in root exchange surface. Additionally, our results indicated that, given the same total C allocation to the root system, the presence of root hairs and the number of lateral roots had minimal impact on the root C balance, whereas a reduction in root tissue density significantly increased rhizodeposition at the expense of root biomass production.

### 5.1 Predicting root-derived carbon emission in soil throughout plant growth

*RhizoDep* successfully simulated realistic belowground C flow dynamics in spring wheat. Using the same C input to roots and an independently parametrized model based primarily on literature data, we reproduced the temporal evolution of C allocation to root production, root respiration, and rhizodeposition, as calculated from field experimental data by Swinnen *et al*. (1994b). All three belowground C flows increased until tillering, then declined from stem elongation to harvest, following the same trend as total C allocation to the root system. A similar decrease in plant rhizodeposition rate at the end of the vegetative period has been documented for wheat (Liljeroth *et al*., 1990*b*), rice (Aulakh *et al*., 2001), and maize (Santangeli *et al*., 2024). Our simulations on spring wheat also revealed that, when expressed per gram of root biomass - as commonly reported in experimental studies - rhizodeposition rate peaked at germination and followed three distinct phases: a continuous decrease in March, an increase in April with the emergence of adventitious roots, and a subsequent decrease from May to August (Fig. S5). A comparable 3-phase pattern in sugar exudation per gram of roots was observed in maize by Matsumoto *et al*. (1979), with an intermediate peak before stem elongation. Similarly, Aulakh *et al*. (2001) found that the exudation rate in rice seedlings was 1 to 10 times higher in early growth stages than at later development stages when expressed per gram of roots.

Root mortality, conceptually distinct from rhizodeposition, was the only belowground C flow that significantly deviated between our simulations and Swinnen’s calculations. According to *RhizoDep,* root mortality became significant only in early June, approximately one month later than expected by Swinnen *et al*. (1994b) (Fig. 2b). However, the precise timing of root mortality remained uncertain in both cases (see section S8). Our results indicate that rhizodeposits were primarily released from May to July, whereas dead roots became the dominant source of root-derived C in the final two months, ultimately accounting for 64% of the total root-derived C input to the soil. Estimates of total root-derived C often rely on allometric relationships, such as the one described by Bolinder *et al*. (2007), which assumes that cumulative rhizodeposited C represents 65% of root biomass C at harvest. *RhizoDep* simulations suggest that, while this 65% proportion remains plausible at harvest, it does not apply consistently across earlier growth stages and is subject to large uncertainty. The accuracy of this estimate depends on how much of the actual dead root mass is included in the standing root biomass measured at harvest (Fig. S22).

Our simulations also provided the first assessment of the relative contributions of distinct rhizodeposition processes throughout the entire growth cycle of a crop. We found that the exudation of soluble compounds accounted for more than 98% of the total C rhizodeposited by spring wheat, while the release of cap cells was a minor contributor, and mucilage secretion remained negligible. This estimation was based on independent parametrization of each rhizodeposition process using data from separate studies, sometimes involving plant species other than wheat. Since no study to date has simultaneously measured all three processes under the same growth conditions, our simulations represent to the best of our knowledge, the first estimation of the relative importance of cap cell release and mucilage secretion compared to root exudation. These results align with previous literature-based conclusions that C release through mucilage secretion and cell sloughing is at least 10 times lower than that from root exudation (Nguyen, 2003; Rees *et al*., 2005).

### 5.2 Deciphering the spatiotemporal patterns of rhizodeposition

A key strength of *RhizoDep* is its ability to simulate spatial variations of rhizodeposition along the roots. In our model, exudation rate varies only due to i) changes in the gradient of hexose concentration between the root symplast and the surrounding soil solution and ii) changes in the exchange surface area between root cells and soil. Our simulations on spring wheat revealed that exudation - and consequently, rhizodeposition - typically peaked at two distinct locations near the growing root tip (Fig. 5). This spatial distribution of exudation near the apex followed the distribution of root mobile hexose rather than variations in root-soil exchange surface area (Fig. S14). However, the decline in exudation rate beyond 4 cm from the growing root tip correlated well with the decrease in root-soil exchange surface area, which offset the small increase in root mobile hexose observed in the region (Fig. S15). In the first seminal axis observed 60 days after germination, the second exudation peak, located 3 cm from the root tip, also coincided with peaks in sucrose unloading and hexose consumption for growth (Fig. 5b), corresponding to the emergence of a lateral root (Video 2). In contrast, the first exudation peak, located around 1 cm from the root tip - just beyond the 7-mm long root elongation zone - did not match with the maxima of sucrose unloading and hexose consumption by growth, which were situated closer to the root tip (Fig. 5b). This suggests that spatial variations in rhizodeposition along a growing root are directly or indirectly linked to preferential C unloading, to sustain either lateral root emergence or root elongation. In the latter case, the delay likely reflects the period required for the root segment to mature out of the elongation zone. Experimental studies on pea (Egeraat, 1975; Nielsen *et al*., 1994) and maize (Mariano and Keltjens, 2003) previously identified two “hot spots” of exudation corresponding to the root extremity and the zone of lateral root emergence. In addition, studies on rapeseed demonstrated that exudation is not maximal at the root tip itself but rather a few mm away from it (Hoffland *et al*., 1989, 1992). These spatial exudation patterns naturally emerged in *RhizoDep* from variations in mobile hexose concentration associated with root growth. The disruption of apoplastic barriers such as endodermis and exodermis during lateral root emergence could also contribute to increased exudation at the second peak. However, our analysis of root-soil exchange surface area suggests that this effect is transient - lasting only a few hours - and of limited amplitude (see Video 2). Thus, higher phloem unloading, required to sustain lateral root emergence, appears to be the primary driver of increased exudation at this site.

Unlike exudation, mucilage secretion and cap cell release were maximal at the root tip. Few studies have investigated the spatial distribution of mucilage secretion, but previous finding indicate that it peaks within the first millimeter of the primary root of maize (Paull and Jones, 1975). Pankievicz *et al*. (2022) similarly reported that mucilage secretion by the aerial roots of a high-secreting maize genotype was highest at the root tip and decreased basipetally, becoming nearly undetectable 4 mm from the tip. Due to the lack of precise data, we modeled the decline in mucilage secretion as a gamma function of distance from the root tip, as previously suggested for exudation by Personeni *et al*. (2007). Additionally, since no evidence supports mucilage secretion by root hairs (Peterson and Farquhar, 1996; Nguyen, 2003), we excluded them from the surface contributing to mucilage secretion. Cap cell release was concentrated at the root tip and ceased beyond the elongation zone. Due to the distinct spatial distributions of the three rhizodeposition processes, their relative contribution to total rhizodeposited C varied along the root. For example, in the first mm of the primary root 60 days after germination, cap cell sloughing and mucilage secretion contributed 10% and 4% of total C rhizodeposition, respectively, but these values dropped to just 1.2% and 0.4% when considering the entire root (Fig. 3a).

Although the first 4 cm of the primary root corresponded to the main rhizodeposition zone along this axis, the rest of the root also contributed significantly. As the primary root elongated then stopped, the proportion of total rhizodeposited C within the first 4 cm decreased from 86% at 30 days to just 8% at 150 days (Table S2). Similarly, at the scale of the root system, the contribution of apical parts of roots to the overall rhizodeposition of the plant decreased over time, from 100% at germination stage to less than 15% at harvest (Fig. S18). This highlights the limitations of simplified approaches that assume a uniform rhizodeposition rate along roots or that restrict rhizodeposition to root tips alone, and fail to capture the complexity of spatial and temporal rhizodeposition patterns.

### 5.3 Quantifying the impact of specific root traits

Beyond its contribution to understanding rhizodeposition patterns, *RhizoDep* serves as a valuable tool for *in silico* experimentation, allowing us to assess how specific root traits affect rhizodeposition. While a full sensitivity analysis was beyond our scope, we examined the relative importance of three root traits that we expected to play a key role for rhizodeposition.

Contrary to our hypothesis, altering root hair density had virtually no impact on rhizodeposition (Table 1). Holz *et al*. (2018) compared a hairless barley mutant to a wild genotype and found that zones with root hairs contributed up to 75% of the total rhizodeposited C, yet the presence of root hairs only marginally increased the mass flux of rhizodeposition per gram of root (by 1% to 7%). More recently, Santangeli *et al*. (2024) investigated a maize root hair defective mutant and observed a slight increase in rhizodeposition per plant at specific growth stages in the presence of root hairs, though the rate of rhizodeposition per unit of root external surface area actually decreased. Our simulations provide insight into these seemingly contradictory findings: while growing root hairs consume additional C, their contribution to the total root-soil exchange surface remains minimal. This total surface area includes not only the external surface but also the accessible symplast surface within the root. Over the entire root system, living root hairs accounted for less than 4% of the total exchange surface area during the first week of growth, less than 2% over the first three months, and even less at later stages. Even within the first 4 cm of the primary root, where rhizodeposition was highest, root hairs contributed at most 8% to the total exchange surface area (Fig. S13). Notably, we assumed that dead root hairs do not contribute to exudate release, though they likely contribute to water uptake (McElgunn and Harrison, 1969). In our simulations, root hairs had a short lifespan, equivalent to 2 days at 20°C (McElgunn and Harrison, 1969; Fusseder, 1987; Nguyen, 2003). If we had considered that dead root hairs continued to released exudates, their contribution to rhizodeposition might have been slightly higher.

Since rhizodeposition is maximal at root tips, we expected that reducing the number of lateral roots would significantly decrease the amount of rhizodeposited C. However, halving the number of lateral roots actually resulted in a 5% increase in rhizodeposited C, while doubling their number led to a slight 3% decrease (Table 1). This outcome underscore the complex relationship between root biomass production and rhizodeposition, particularly when accounting for root C balance within distinct root orders (Galindo-Castañeda *et al*., 2024). Reducing lateral root density limited potential growth, leading to an accumulation of mobile C within the root, which in turn stimulated root exudation despite the decrease in the number of root tips. Furthermore, the short growth duration of most lateral roots may have constrained their contribution to rhizodeposition, as they received considerably less C once they ceased elongating. Experimental studies on this relationship are limited. Henry *et al*. (2005) observed that increasing N availability in *Lolium multiflorum* grown in sand reduced root biomass but increased the number of root apices, which coincided with higher rhizodeposition per plant. Aoki *et al*. (2012) also reported a positive correlation between organic acid exudation rate per gram of root biomass and the number of root apices in tropical trees. Conversely, Meier *et al*. (2013) found a negative correlation between C exudation rate per gram of root biomass and the number of root apices in loblolly pine, possibly due to pathogenic fungal infection.

Among the three investigated traits, root tissue density (RTD) had the most significant impact, highlighting a potential trade-off between root biomass production and rhizodeposition. When RTD was halved, the C cost of root production decreased accordingly, which led an accumulation of mobile C that was not used for growth, resulting in an 83% increase in rhizodeposited C. This finding aligns with the “root economic spectrum” theory, which associates low RTD with an “acquisitive” strategy and high RTD with a “conservative” strategy (Roumet *et al*., 2016). When considering root exudation within this framework, Wen *et al*. (2022) identified a low RTD as one of the best predictors of high-exuding plant species. A negative correlation between RTD and exudation has also been documented within a single plant species, as demonstrated in two maize genotypes grown in different soils (Santangeli *et al*., 2024).

*RhizoDep* provides a mechanistic foundation for interpreting the effects of such root traits on rhizodeposited C, considering the interaction between root C balance and the evolution of root structure. However, it is important to note that our conclusions here are specific to wheat and do not account for possible feedback mechanisms, such as changes in root architecture influencing C supply from the shoots via enhanced soil nutrient uptake.

### 5.4 Limitations and possible improvements

While our study successfully simulated the expected belowground C flows of spring wheat, several limitations of our approach should be mentioned. First, the simulation time - approximately 16 hours on a single computer to simulate 6 months in the reference scenario (Sc1) - could pose challenges when numerous simulations are needed, e.g. for exploring environmental effects or conducting extensive sensitivity analyses. Since simulation time correlates directly with the number of root elements in *RhizoDep*, efficiency could be improved by optimizing root segment sizes (e.g., using variable sizes along the roots) or parallelizing computation across different root groups.

Another limitation arises from the large number of parameters in this mechanistic model. Of the 31 parameters related to root architecture, most can be measured from experimental observations of root systems. However, approximately 100 parameters associated with root metabolism are not readily measurable without specific experiments. In this study, we primarily relied on literature data, noting that 25% of parameter values were unavailable and thus were estimated based on their expected similarities with other parameters. This parametrization highlights several research gaps, e.g. the need for better quantification of phloem unloading, root mortality, mucilage secretion, and cell sloughing. Although our study focused on spring wheat, *RhizoDep* was designed as a generic root model. We hypothesize that most root metabolism parameters are conserved across plant species, eliminating the need for extensive re-parametrization in future applications. Validating the model with other species will test this assumption, though experimental datasets comparable to Swinnen *et al*. (1994b) remain scarce.

In its current form, *RhizoDep* models only the dynamics of sugars and their effects on root growth and rhizodeposition, simplifying sucrose transport and assuming that mobile hexose represent all soluble compounds available for exudation. Integrating carboxylates - a key component of exudates - would enable the model to account for additional soil-plant interactions, such as the solubilization of soil mineral phosphate or the destabilization of mineral-associated organic matter. Furthermore, integrating N metabolism (e.g. N uptake, transformation, and transport) and symbiotic associations in the model will allow exploration of diverse plant strategies along the root economic spectrum.

## 6 Conclusions

*RhizoDep* offers a novel tool for investigating the spatial and temporal dynamics of belowground C flows in detail, enabling the identification of conditions that favor rhizodeposition and/or root biomass production. Using spring wheat as a case study, we demonstrated that root exudation is the predominant rhizodeposition process, with its spatial distribution primarily governed by the preferential unloading of sugars to support root growth. Root traits influencing root architecture had a lesser effect on rhizodeposition compared to traits affecting the consumption of C for growth. Our findings also revealed that simpler modelling approach—such as prescribing a uniform rhizodeposition rate along roots or restricting it to root tips - will fail to capture realistic root C dynamics. Moving forward, we plan to couple *RhizoDep* with a shoot model and a soil model to incorporate additional feedbacks between root architecture and metabolism, such as increased C allocation to roots triggered by an enhance soil nutrient uptake. We anticipate that *RhizoDep*, along with its future refinements, will shed new light on the complex interactions within the soil-plant-atmosphere continuum.

## 7 Supplementary data

Additional information is available in following files:

- SI: This document contains details about the model, its parametrization for wheat, and the calculations of belowground C flows from the data of Swinnen et al. (1994b), and provides complementary simulation results.
- ST: This spreadsheet provides the list of the main variables in the model and their units, as well as the list of parameters and their estimated values for spring wheat.

## 8 Ackowledgements

We thank Loïc Pagès for his advice regarding *ArchiSimple*, Elisa Clerjeault and Soukeye Gadiaga for helping to determine root architecture parameters, and Raia Silvia-Massad for hosting a delightful writing retreat.

## 9 Authors contribution

FR designed and performed the research, and wrote the manuscript with the help of all authors. FR, BA and TG conceptualized the model, and FR, CP, and MG implemented the model.

## 10 Conflict of interest

No conflict of interest declared.

## 11 Funding

This work benefited from the discussions hold within the IMPULSE project (2019-2022) funded by INRAE and the project MIXROOT-C (2021-2024) funded by the European Union’s Horizon 2020 research and innovation program under grant agreement No. 862695 EJP SOIL.

---

**Embedded Video 1:**
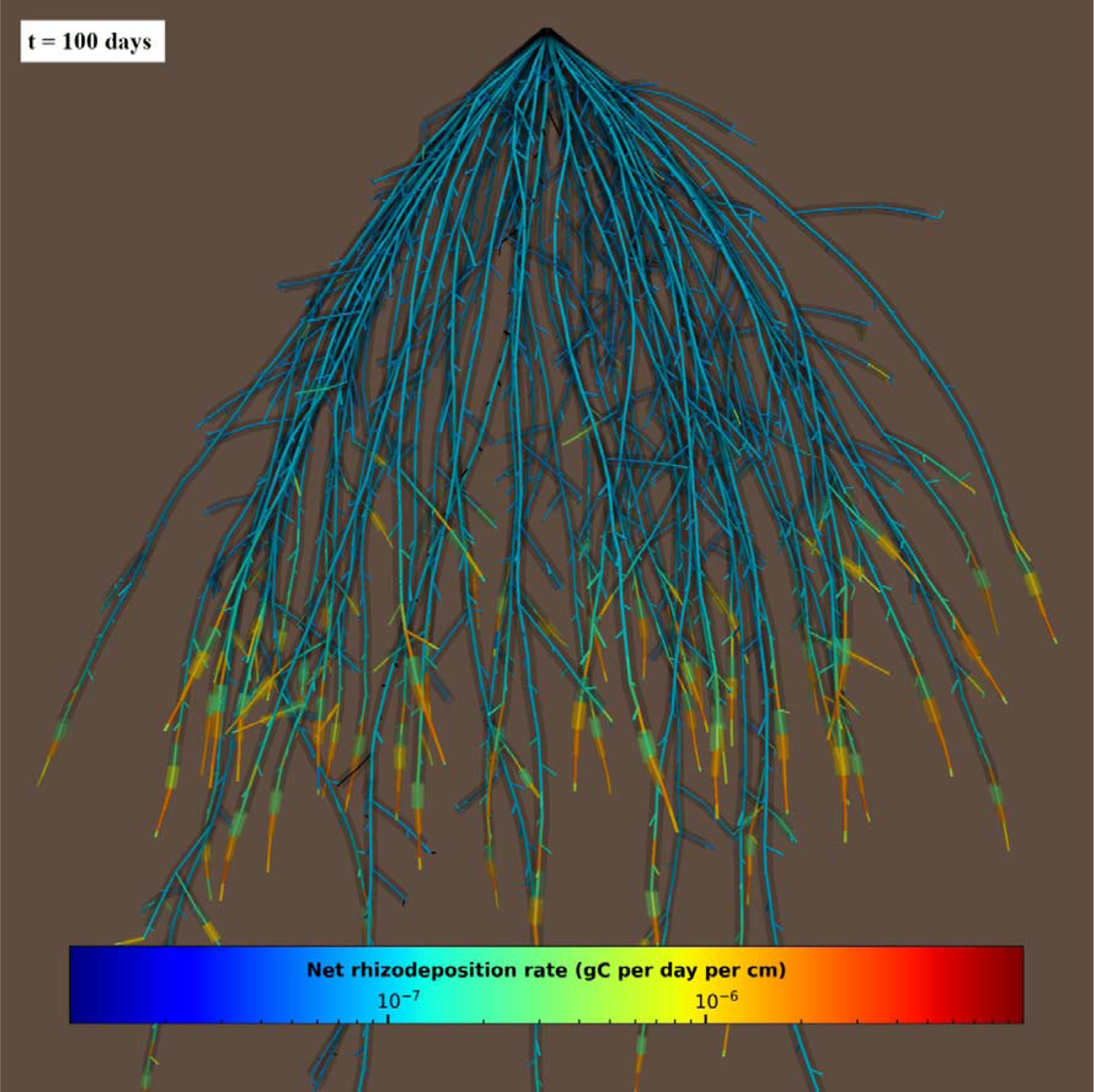
Visualization of the dynamic growth and rhizodeposition of spring wheat’s root system between March and August 1989, as simulated by *RhizoDep* in scenario Sc1 based on the work of Swinnen *et al*. (1994b). Colors represent the net rhizodeposition rate (in gC per cm per day) along each root over time. The volume of living root hairs is represented as a semi-transparent colored cylinder, while dead roots and dead root hairs are shown in black.

**Embedded Video 2:**
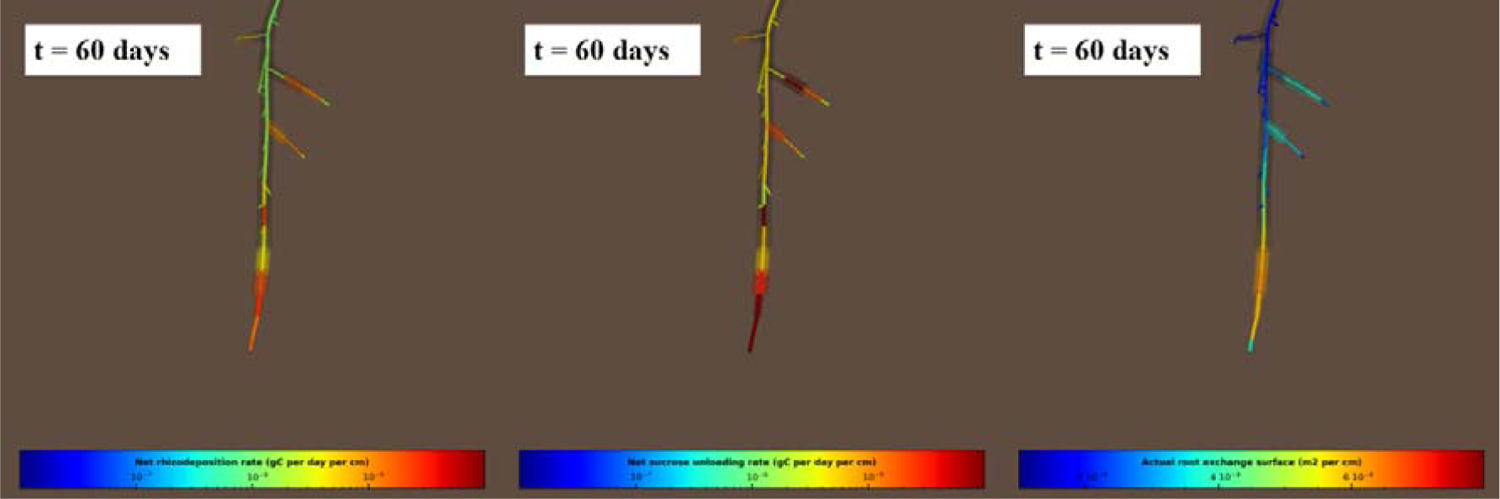
Distribution of the rate of rhizodeposition (left), phloem unloading (middle) and root-soil exchange surface area (right) along the first seminal root of spring wheat from 55 to 65 days after germination, as simulated by *RhizoDep* in scenario Sc1 based on the work of Swinnen *et al*. (1994b). The volume of living root hairs is represented as a semi-transparent colored cylinder, while dead roots and dead root hairs are shown in black.

## Notes

https://github.com/openalea/rhizodep

### Competing Interest Statement

The authors have declared no competing interest.

