## Supporting Information file SI for "Deciphering spatiotemporal patterns of rhizodeposition with a functional-structural root model: *RhizoDep*"

**Supporting Information for the manuscript *Exploring spatiotemporal patterns of rhizodeposition with a functional-structural root model: RhizoDep***

Frédéric Rees^1*^, Tristan Gérault^1^, Marion Gauthier^1^, Romain Barillot^2^, Céline Richard-Molard^1^, Alexandra Jullien^1^, Claire Chenu^1^, Christophe Pradal^3,4^, Bruno Andrieu^1^

^1^Université Paris-Saclay, INRAE, AgroParisTech, UMR EcoSys, 91120 Palaiseau, France; ^2^INRAE, UR P3F, F-86600 Lusignan, France; ^3^CIRAD, UMR AGAP, and Inria, F-34398 Montpellier, France; ^4^AGAP, Univ Montpellier, CIRAD, INRAE, Montpellier SupAgro, F-34398 Montpellier, France

### 1 Formalism of specific processes in *RhizoDep*

Several metabolic functions have been implemented in the model, and are described below, followed by a brief justification of the formalism that was chosen.

#### 1.1 Supply of sucrose & distribution within the root system

The supply of C fixed by photosynthesis in the aerial parts and transferred to the root system represents the main input variable of *RhizoDep*, i.e. it has to be specified by the user or simulated by another model. This supply of photoassimilates is expressed as the net exchange of sucrose from shoots to roots over a given period (moles of sucrose supplied per second). This net input of sucrose is spread over the whole root system without any delay or resistance, and the same, new sucrose concentration (moles of sucrose per gram of root structural mass) can be uniformly applied to each root element, using a mass balance over the whole root system. First, the initial total amount of sucrose in the root system is calculated by summing the amount of sucrose present in each root element (product of the local sucrose concentration with the root structural mass of the root element). The additional amount of sucrose supplied by the shoots over a given time step is then summed with this initial sucrose, and a new average concentration of sucrose is calculated by dividing the new total amount of sucrose with the total structural mass of the root system.

This simplified representation of sucrose transport relies on two important assumptions: i) the concentration of sucrose tends to be uniform over the whole root system, ii) any addition of sucrose at the base of the root system results in a rapid spreading and re-homogenization of sucrose concentration within the whole root phloem network. Such assumptions apparently contradict modelling approaches that simulate a gradient of sugar concentration along the roots by considering a resistance to the transport of sucrose that increases with root length (Bidel *et al.*, 2000). Our assumptions may also seem incompatible with the fact that the C fixed in the shoots can take several hours or days to reach the end of a root (Kuzyakov & Gavrichkova, 2010). However, our assumptions can reconcile with these observations when considering the existence of pressure-concentration waves that propagate along the phloem vessels much faster than the phloem’s solution itself, at least when the sieve tube’s axial pressure drop is low (Thompson & Holbrook, 2004). This theory predicts that in such case, an addition of sucrose at one end of a phloem vessel results in a rapid homogenization of the concentration in the whole vessel, even if the transport of the recently photoassimilated C remains slow. This may explain the relative lack of spatial variations of the sucrose concentration along the mature parts of the roots reported for small plants such as maize (Hellebust & Forward, 1962; Sharp *et al.*, 1990; Jones & Darrah, 1996), pea (Lyne & ap Rees, 1971), and Arabidopsis (Freixes *et al.*, 2002), and for larger plants such as ricinus (Chapleo & Hall, 1989).

#### 1.2 Net exchange of sugars between phloem vessels and the rest of the root cells

Sucrose can be unloaded from the phloem vessels in any living root element *i* and converted into hexose. These two processes are described by a single equation, based on the gradient of sugar concentration:

$$\begin{aligned} \frac{d{Hexose}_{production \left( i \right)}}{dt}= 2\cdot P_{phloem \left( i \right)}(t)\cdot S_{phloem \left( i \right)}\left( t \right)\cdot\left( \left[ Sucrose \right]_{phloem \left( i \right)} \left( t \right)-oemle equation, based on the gradient of concentrationn and to the concentrations of hexose in the mobile pool \frac{\left[ Hexose \right]_{symplast \left( i \right)} \left( t \right)}{2} \right) \end{aligned}$$

(Eq. S1 )

where ${Hexose}_{production \left( i \right)}$ (moles of hexose) is the amount of hexose generated by the complete hydrolysis of the sucrose that has been unloaded from the phloem in that particular root element, *P_phloem (i)_ (t)* is the permeability coefficient of phloem’s vessels membranes to sucrose within the root element *i* (per square meter and per second), *S_phloem (i)_ (t)* is the total surface of exchange of all phloem vessels within the root element *i* (square meter), *[Sucrose]_phloem (i)_* (*t*) is the concentration of sucrose in the root element *i* at time *t* (moles of sucrose per gram of root structural mass), and *[Hexose]_symplast (i)_* (*t*) is the symplastic concentration of mobile hexose in the root element *i* at time *t* (moles of hexose per gram of root structural mass). The factor 2 corresponds to the conversion of sucrose into hexose.

Eq. S1 implicitly assumes that all the sucrose unloaded from the phloem is converted into hexose and that no sucrose is stored outside the phloem compartment. While this contradicts evidence that sucrose is not confined to phloem and can be stored in the vacuoles inside cortical or epidermal root cells (Ho, 1988; Hedrich *et al.*, 2015), this rough spatial segregation between sucrose and hexose is actually supported by the high concentrations of sucrose observed in phloem tissue compared to the predominance of hexose in the cortex, as shown for example in pea roots (Lyne & ap Rees, 1971). Eq. S1 also assumes that the unloading rate is proportional to the gradient of sucrose concentration between the phloem vessels and the symplast of root non-vascular cells. This simplified vision is in line with the conclusion that unloading is mainly passive and occurs by diffusion through plasmodesmata or into phloem’s apoplasm, both driven by the concentration of sucrose in the sieve tubes (Patrick, 1997; Lalonde *et al.*, 2003; De Schepper *et al.*, 2013).

In order to ensure that the sucrose unloading would closely match the needs of hexose for growth, we introduced a linear dependency of the permeability coefficient with the rate of hexose consumption by growth:

$$\begin{aligned} P_{phloem \left( i \right)}(t)= P_{phloem ref}\cdot\left( 1+oemle equation, based on the gradient of concentrationn and to the concentrations of hexose in the mobile pool \frac{\left( \frac{d{Hexose}_{growth \left( i \right)}}{dt} \right)(t-dt)}{\left( \frac{d{Hexose}_{growth}}{dt} \right)_{ref}} \right) \end{aligned}$$

(Eq. S2)

where *P_phloem ref_* is the reference permeability coefficient of phloem’s vessels membranes to sucrose in the absence of growth (per square meter and per second), and the two terms of the ratio correspond to the rate of hexose consumption for sustaining growth for the element *i* at the previous time step, and to an equivalent rate of hexose consumption for which the potential unloading of sucrose is doubled (moles of hexose per second).

Conversely, sucrose can also be synthesized from the available hexose in the symplast and reloaded into the phloem vessels following a Michaelis-Menten formalism, as shown in Eq. S2:

$$\begin{aligned} \frac{d{Sucrose}_{production \left( i \right)}}{dt}= {Loading}_{max}\cdot S_{phloem \left( i \right)}\left( t \right)\cdot\frac{\left[ Hexose \right]_{symplast \left( i \right)} \left( t \right)}{K_{loading}+ \left[ Hexose \right]_{symplast \left( i \right)} \left( t \right)} \# \end{aligned}$$

(Eq. S3)

where *Sucrose _production_ _(i)_* (moles of sucrose) is the amount of sucrose synthetized over time from hexose and reloaded in the phloem in that particular root element *i*, *Loading_max_* is the maximal surfacic rate of sucrose loading (moles of sucrose per square meter per second), *[Hexose]_symplast (i)_* (*t*) is the symplastic concentration of mobile hexose in the root element *i* at time *t* (moles of hexose per gram of root structural mass), and *K_loading_* represents the concentration of hexose for which sucrose synthesis and reloading is equal to half of the maximal rate. Such reloading enables to account for the mechanism of leakage-retrieval that is observed along the phloem, e.g. in basal parts of the root system (De Schepper *et al.*, 2013).

#### 1.3 Root growth, structural mass production and growth respiration

In the current version of *RhizoDep*, we have adapted the growth rules implemented in *ArchiSimple* and described in details by Pagès et al. (2014). Only the main principles of *ArchiSimple*’s rules are stated here:

- Any root element can grow longitudinally (elongation) or radially (thickening), or disappear (root decay and abscission). New root elements can appear in the root system in two distinct situations: i) when the primordium of a new potential lateral root is formed, ii) when a root tip elongates and its length becomes higher than the prescribed root segment length *l_segment_*, in which case the tip is segmented in different root segments and a final root tip, which length is lower than *l_segment_*.
- The primordia of lateral roots are formed at the root tip of the “mother” root, and a regular, constant distance *d_interprimordia_* is conserved between the primordia of the same axis. One of the main characteristics of *ArchiSimple* is the absence of distinction between types or orders of roots. Thus, instead of prescribing distinct properties according to the type of roots, *ArchiSimple* considers that most of the properties derive from the diameter of the root tip. The diameter of the new primordia is drawn from a statistical distribution centered on a mean diameter that is proportional to the diameter of the mother root, and the new primordium can only emerge after a prescribed dormancy period. The potential elongation rate of a root tip is also proportional to the diameter of the root tip. However, the period for which a root tip can continue to elongate and the time for which it remains alive are both proportional to the square diameter of this root tip (Pagès *et al.*, 2014).
- The trajectory of each root element in 3D space and the actual elongation also depend on a vertical and horizontal tropism parameter, and on a soil friction parameter representing the reduction of elongation by mechanical constraints. Radial growth is based on the geometrical rule of the “pipe model”, which states that the area of the section of a root element must be proportional to the sum of the sections of the adjacent root elements located distally from it, so that the diameter of a root element is always equal to or higher than the diameter of the distal root elements.
- Eventually, the abscission of a root occurs when the apex of this root and the apices of every lateral roots supported by this root have reached their life duration, as prescribed by their diameter.

The volume and the structural mass of a given cylindrical root element *i*, can be defined as:

$$v_{\left( i \right)}(t)=\pi{\cdot\left( r_{\left( i \right)}\left( t \right) \right)}^{2}{\cdot l}_{\left( i \right)}\left( t \right)$$

(Eq. S4)

$$m_{struc \left( i \right)}\left( t \right)=v_{\left( i \right)}\left( t \right)\cdot{density}_{root (i)}\left( t \right)$$

(Eq. S5)

where $v_{\left( i \right)}(t)$ is the volume of the root element *i* (m^3^), $r_{\left( i \right)}$ its radius (m), ${. l}_{\left( i \right)}\left( t \right)$ its length (m), $m_{struc \left( i \right)}\left( t \right)$ its dry structural mass (grams), and ${density}_{root (i)}\left( t \right)$ is its root tissue density (grams of structural mass per m^3^ of root).

As in *ArchiSimple* model, the elongation potential of each root apex is proportional to the radius of the apex, but in *RhizoDep* an additional regulation of the elongation rate by the concentration of locally available hexose in the root is made:

$$\frac{dl_{\left( i \right)}}{dt}={elongation}_{max}\cdot r_{\left( i \right)}\left( t \right)\cdot\frac{\left[ Hexose \right]_{available (i)} (t)}{K_{elongation}+\left[ Hexose \right]_{available \left( i \right)}(t)}$$

(Eq. S6)

where $l_{\left( i \right)}$ is the length of the root element *i* (m), ${elongation}_{max}$ is the maximal rate of root elongation per unit of root radius (s^-1^), $K_{elongation}$ represents the concentration of hexose for which half of the maximal rate is reached, and $\left[ Hexose \right]_{available (i)} (t)$ corresponds to the average concentration of hexose available in the growth-supplying zone associated to the element *i* (moles of hexose per gram of root structural mass).

The growth-supplying zone for elongation involved in *Eq. S6* corresponds to the root volume in which mobile hexose can be taken and used to sustain growth; its volume is defined as:

$$v_{growth-supplying \left( i \right)}\left( t \right)=\pi{\cdot\left( r_{\left( i \right)}\left( t \right) \right)}^{2}\cdot r_{\left( i \right)}\left( t \right)\cdot\alpha_{growth-supply}$$

(Eq. S7)

where $\alpha_{growth-supply}$ is an empirical coefficient (dimensionless) expressing the ratio between the theoretical length of the growth-supplying zone - assuming it is located on the same root axis as element *i* - and the radius of the root element *i*. For instance, if we assume that the root meristem and the root elongation zone together represent the growth-supplying zone where hexose can be used to sustain root elongation, and if their cumulative length corresponds to 8 times the radius of the meristem, then $\alpha_{growth-supply}$ would be equal to 8. Hence, the growth-supplying zone may encompass not only the root element *i* itself, but also several preceding root elements, including elements from a mother root (see Fig. S1). This concept of a growth-supplying zone independent on root segmentation was introduced in *RhizoDep* so that the supply of hexose for growth remains virtually independent on the (variable) size of the actual growing root tip.

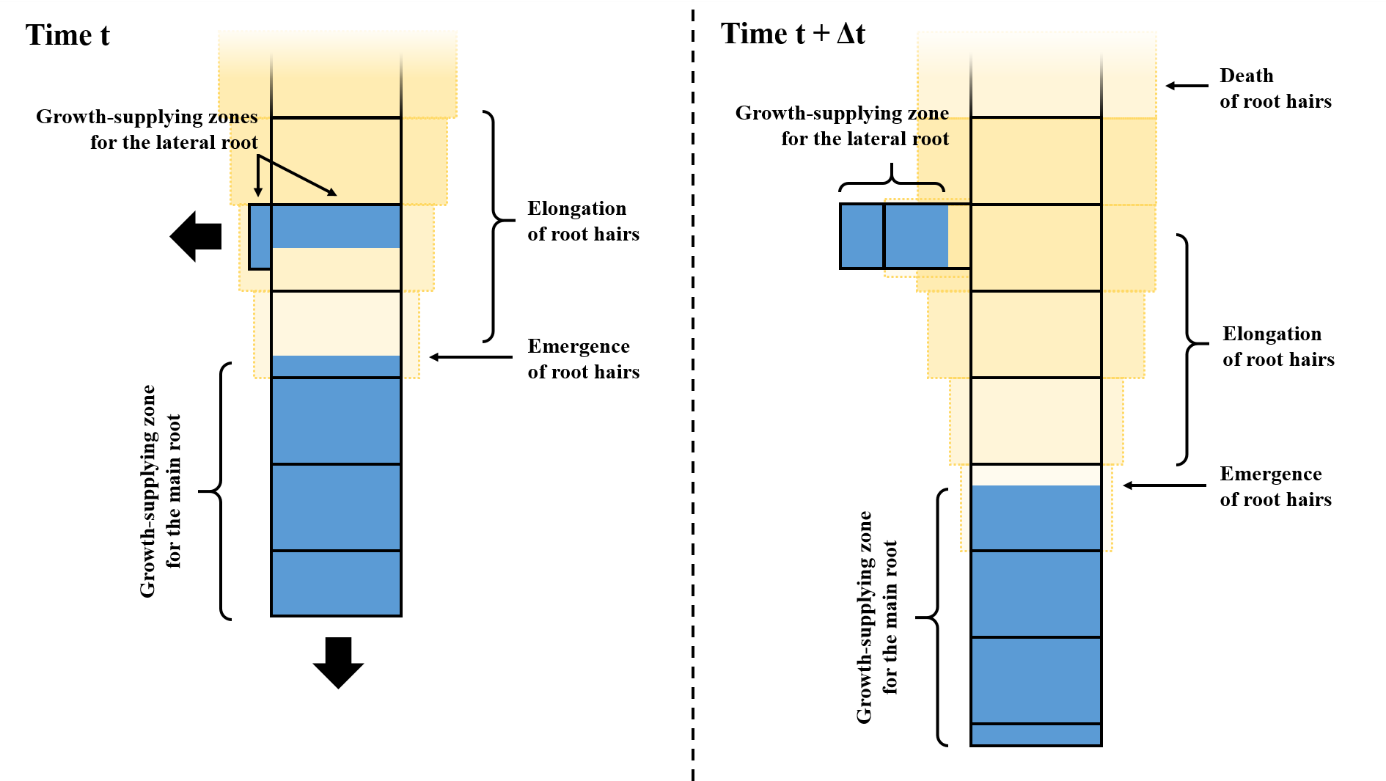

***Fig. S1: Spatial discretization of the elongation of root axes and root hairs in the model* RhizoDep*.*** *The elongation corresponds to the transformation of hexose into structural mass for producing the new root segments; for a given root axis, this hexose can be taken from the entire growth-supplying zone (in blue), which length is proportional to the root diameter and which may encompass several root elements. For a lateral root, the growth-supplying zone can include part of the mother root segment. Root hairs start to elongate right after the growth-supplying zone and grow until reaching their maximal length. The density (represented by the intensity of yellow color) of living root hairs on a given segment depends on their average age, and starts to decrease when the age of the first root hairs to have emerged on this segment gets higher than root hairs’ lifespan.*

The potential thickening rate of a root segment is dependent on both its initial radius and the concentration of mobile hexose in this root element:

$$\left( \frac{dr_{\left( i \right)}}{dt} \right)_{hexose}={thickening}_{max}\cdot r_{\left( i \right)}\left( t \right)\cdot\frac{\left[ Hexose \right]_{symplast (i)}}{K_{thickening}+\left[ Hexose \right]_{symplast (i)}}$$

(Eq. S8)

However, thickening also occurs according to the pipe model, which states that the section of a root element must be large enough to support the supply to (or collection from) the roots located distally from it. Mathematically, this means that the section *S* _(_*_i_*_)_ of a given root segment *i* is calculated proportionally to the sum of the section of the next segment on the same root *S* _(_*_i+1_*_)_ and of a fraction of the sum of the sections of the lateral roots directly supported by the root segment:

$$S_{\left( i \right)}= S_{\left( i+1 \right)}+ p\cdot\sum S_{lateral (i)}$$

(Eq. S9)

where *p* is a coefficient, which can be close to 1 for dicotyledonous plants and to 0 for monocotyledonous species, which do not support root radial growth (Pagès *et al.*, 2014). The pipe model therefore sets a geometrical condition on the variation of root radius over time. Eventually, the final variation is calculated as the minimum value between what is controlled by the maximal possible rate of growth and what is geometrically possible:

$$\frac{dr_{\left( i \right)}}{dt}=min\left( \left( \frac{dr_{\left( i \right)}}{dt} \right)_{hexose}, \left( \frac{dr_{\left( i \right)}}{dt} \right)_{geometry} \right)$$

(Eq. S10)

Whether it corresponds to root elongation or thickening, root growth requires a specific amount of hexose for: i) building the new structural dry biomass that corresponds to the increase in volume of the root element, based on a pre-determined root structural mass density, and ii) covering energy costs linked to root growth, generating a so-called “growth respiration” (Thornley & Cannell, 2000). The variation of hexose required for the growth of a root element *i* is therefore proportional to the variation of its volume:

$$\frac{d{Hexose}_{growth (i)}}{dt}= \frac{dv_{(i)}}{dt} \cdot\frac{{density}_{root (i)}\left( t \right) \cdot struct\_mass\_C\_content}{6\cdot{yield}_{growth}}$$

(Eq. S11)

where *Hexose _growth (i)_* (*t*) is the total amount of hexose (moles of hexose) consumed by growth processes, *v_i_* is the volume of the root element *i* (in m^3^), *density _root (i)_* is the apparent dry density of the root element *i* (grams of structural mass per m^3^ of root), *struct_mass_C_content* is the content of C in the root structural mass (moles of C per gram of dry structural biomass), and *yield _growth_* is the molar fraction of C that is effectively used for building structural mass according to Thornley and Cannell (2000). The coefficient 1/6 corresponds to the conversion factor between moles of C and moles of hexose.

The amount of CO_2_ generated by this growth (moles of CO_2_) is calculated accordingly:

$$\frac{d{{CO}_{2}}_{growth (i)}}{dt}=6\cdot\frac{d{Hexose}_{growth (i)}}{dt} \cdot\left( 1-{yield}_{growth} \right)$$

(Eq. S12)

In *RhizoDep*, we implemented a double control of growth by C. Besides the limitation of the rate of elongation or thickening by the concentration of hexose, the growth a root element *i* is also limited when the amount of locally available hexose at time *t* is lower than the required amount of hexose to cover all potential growth costs associated to the root element *i*. In such case, the amount of locally available hexose is converted into a maximal volume of new root structural mass that can be produced, by reversing *Eq. S10*. This maximal volume is then used for growth. In case several types of growth may occur within the same root element (e.g. root thickening while sustaining the emergence of a lateral primordium), growth priority rules apply. If there is not enough hexose available for reaching the potential elongation, elongation of the root element is done up to the maximal possible length. If any hexose remains available once elongation has been considered, the emergence of a new primordium can occur, and, if any hexose remains, root thickening can occur. Such priority rules among different growth possibilities have been used in other models, although not with the same priorities (Thaler & Pagès, 1998; Postma & Lynch, 2011), while other root models eventually decided to apply an indistinct, identical C-limitation for each simultaneous growth process occurring on a root element (Thaler & Pagès, 1998; Pagès *et al.*, 2014). As our simulations on wheat did not involve radial growth, these growth priorities did not play any role in the present work.

#### 1.4 Root hairs dynamics

In *RhizoDep*, root hairs are not individually described, but are represented as an extension of root external surface and root structural mass, distinguishing living root hairs and dead root hairs. For each root element, root hairs are defined by their density (number of hairs per meter of root) and by their average length (meter), diameter (meter) and age (second). Root hairs are assumed to emerge right above the elongation zone of a root axis, which length is defined using the parameter *α_growth_supply_* already described above for roots. Because the position of the root element where the first roots hairs emerge evolves over time relatively to the upper limit of the root elongation zone of the axis, the density of root hairs also progressively increases until reaching the maximal root hairs density (see Fig S1). Root hairs elongate until reaching their average maximal length *l_hair_max_* (meter). The increase in the average root hair length of living root hairs is calculated according to a maximal rate of hair elongation, and regulated by the concentration of hexose available within the fraction of the supporting root element that is covered with root hairs:

$$\frac{dl_{hairs \left( i \right)}}{dt}={hair\_elongation}_{max}\cdot r_{hairs}\cdot\frac{\left[ Hexose \right]_{symplast (i)} \left( t \right)\cdot\frac{{hair\_zone\_length}_{(i)}}{l_{\left( i \right)}}}{K_{hair elongation}+\left[ Hexose \right]_{symplast \left( i \right)}(t)}$$

(Eq. S13)

where $l_{hairs \left( i \right)}$ is the length of the root element *i* (m), ${hair\_elongation}_{max}$ is the maximal rate of root hair elongation per unit of root hair radius (s^-1^),$r_{hairs}$ is a parameter specifying the average radius of each root hair (m), $\left[ Hexose \right]_{symplast (i)} (t)$ corresponds to the symplastic concentration of mobile hexose available in the supporting root element *i* (moles of hexose per gram of root structural mass)*,* ${hair\_zone\_length}_{(i)}$ is the actual length of the root hair zone covering the root element *i* (m), $l_{\left( i \right)}$ is the length of the root element *i* (m), and $K_{hair elongation}$ represents the concentration of hexose for which half of the maximal rate is reached.

Following their emergence, root hairs remain alive for a pre-defined period *root_hairs_lifespan* (second), and then die. We assume that dead root hairs remain attached to the supporting root element, and therefore are not sloughed-off until the root element itself dies, nor replaced by new root hairs (Fusseder, 1987; Nguyen, 2003).While root hair sloughing does not directly contribute to rhizodeposition in *RhizoDep*, root hairs dynamics influences C balance (as a growth and maintenance costs are associated to living root hairs) and hexose exudation or uptake from soil because of the extension of root exchange surface by living root hairs.

#### 1.5 Root maintenance respiration

Root maintenance is represented in our model by a single equation describing the consumption of hexose for covering all costs relative to root maintenance processes, based on a Michaelis-Menten formalism:

$$\frac{d{Hexose}_{maintenance (i)}}{dt}= R_{max}\cdot m_{struct (i)}\left( t \right)\cdot\frac{\left[ Hexose \right]_{symplast(i)} (t)}{K_{maintenance}+ \left[ Hexose \right]_{symplast(i)} (t)}$$

(Eq. S14)

where *Hexose _maintenance (i)_* (*t*) (moles of hexose) is the amount of hexose used for maintenance respiration in the root element *i*, *R_max_* is the maximal rate of maintenance respiration (moles of hexose per gram of dry structural biomass per second), *m_struct (i)_* (*t*) is the dry structural mass of the root element *i*, *[Hexose]_symplast (i)_* (*t*) is the symplastic concentration of mobile hexose in the root element *i* (moles of hexose per gram of root structural mass), and *K_maintenance_* represents the concentration of hexose for which the rate of maintenance respiration is equal to half of the maximal rate.

Hence, in our model, maintenance is translated as a C cost that decreases the amount of hexose in a given root element. The steps of hexose conversion into energy (e.g. through ATP synthesis) and the use of this energy for specific maintenance processes are not explicitly represented. The maintenance cost is assumed to be proportional to the dry structural mass and to increase with the concentration of hexose in the root element. This illustrates the fact that a larger dry structural mass represents a higher number of root cells to be maintained, and that a higher concentration of hexose necessarily implies a higher C cost for handling the transport, storage or transformation of this hexose within the root element. Maintenance costs have been associated to many processes, *e.g.* phloem loading, nitrate uptake and conversion into organic N (Barillot *et al.*, 2016), and should therefore depend on additional variables and processes that have not been represented in *RhizoDep* so far. However, even in models that explicitly calculate maintenance costs based on such processes (Thornley & Cannell, 2000; Barillot *et al.*, 2016), a “residual” maintenance cost has to be calculated in order to give a reliable estimation of maintenance respiration. In many cases, this residual maintenance cost represents up to 50% of the total maintenance cost (Thornley & Cannell, 2000). We have therefore chosen to use a similar Michaelis-Menten formalism as the one suggested by Thornley and Cannell (2000) for calculating this residual maintenance, and considered that this formalism covers all maintenance costs.

The hexose consumed by root maintenance is then fully converted into respired CO_2_:

$$\frac{d{CO2}_{maintenance (i)}}{dt}= 6\cdot\frac{d{Hexose}_{maintenance (i)}}{dt}$$

(Eq. S15)

where *CO_2_ _maintenance (i)_* (moles of CO_2_) represents the amount of CO_2_ respired by the root element *i* because of maintenance processes.

#### 1.6 Exchange of sugars with the reserve pool

*RhizoDep* simulates a root reserve pool, which can be build up inside the non-vascular root cells. This reserve is considered in the model as one single pool characterized by its own concentration of hexose, *[Hexose]_reserve_* (moles of hexose per gram of dry structural biomass), although this reserve pool may be made from several constituents, such as the vacuole of root cells or individual amyloplasts. The immobilization process that enables to store mobile hexose into this reserve for any root element *i* is described as:

$$\frac{d{Hexose}_{immobilization (i)}}{dt}= {Immob}_{max}\cdot m_{struct \left( i \right)}(t)\cdot\frac{\left[ Hexose \right]_{symplast (i)} (t)}{K_{immobilization}+ \left[ Hexose \right]_{symplast (i)} (t)}$$

(Eq. S16)

where *Hexose _immobilization_ _(i)_* (moles of hexose) is the amount of hexose entering the reserve pool over time in the root element *i*, *Immob_max_* is the maximal rate of immobilization (moles of sucrose per gram of dry structural mass second), *m_struct (i)_ (t)* is the dry structural mass of root element *i* at time *t* (gram), *[Hexose]_symplast (i)_* (*t*) is the symplastic concentration of mobile hexose in the root element *i* at time *t* (moles of hexose per gram of root structural mass), *K_immobilization_* represents the concentration of hexose for which hexose immobilization is equal to half of the maximal rate. Note that if *[Hexose]_symplast (i)_* (*t*) is lower than a certain threshold concentration *[Hexose]_symplast min_*, or if *[Hexose]_reserve (i)_* (*t*) is higher than a certain threshold concentration *[Hexose]_reserve max_*, no hexose immobilization can occur.

Conversely, hexose can also be remobilized from the reserve and re-enter the mobile pool of hexose in the symplast:

$$\frac{d{Hexose}_{mobilization (i)}}{dt}= {Mob}_{max}\cdot m_{struct \left( i \right)}(t)\cdot\frac{\left[ Hexose \right]_{reserve (i)} (t)}{K_{mobilization}+ \left[ Hexose \right]_{reserve (i)} (t)}$$

(Eq. S17)

where *Hexose _mobilization_ _(i)_* (moles of hexose) is the amount of hexose mobilized over time from the reserve in root element *i*, *Mob_max_* is the maximal rate of hexose mobilization (moles of hexose per second), *[Hexose]_reserve (i)_* (*t*) is the concentration of hexose in the reserve pool of the root element *i* at time *t* (moles of hexose per gram of root structural mass), and *K_mobilization_* represents the concentration of hexose in the reserve for which hexose mobilization is equal to half of the maximal rate.

We chose to simplify immobilization and mobilization processes by a similar, unique Michaelis-Menten kinetics based on the sugar concentration in the mobile pool of hexose and in the reserve pool, respectively. Our model for this was the exchange of sugars with the vacuole through the tonoplast, which has been shown to involve a number of transmembranar transporters for facilitating the transport along or against the concentration gradient (Hedrich *et al.*, 2015), even if more complex behaviors may be expected in reality, such as the biphasic pattern (curvilinear evolution of the rate of transport at low concentrations, and linear evolution at high concentrations) described by Etxeberria et al. (2012).

#### 1.7 Exudation and uptake of hexose and sucrose

The net efflux of hexose from a living root element into the soil is described as the difference between the gross exudation of hexose, which is governed by a diffusion gradient, and an influx of hexose from the soil interface into the root considered as an active process (Jones & Darrah, 1993; Farrar *et al.*, 2003; Personeni *et al.*, 2007). The diffusion of sugars from root cells and their re-uptake are considered both for non-vascular root cells and for phloem vessels, as long as there can be a direct contact between the soil solution and those cells. Such contact depends on the existence of transport barriers (endodermis and exodermis) (see below). Sugar exudation and uptake are described by following equations:

$$\frac{d{Hexose}_{net efflux (i)}}{dt}=\frac{d{Hexose}_{exudation (i)}}{dt}- \frac{d{Hexose}_{uptake (i)}}{dt}$$

(Eq. S18)

$$\frac{d{Sucrose}_{net efflux (i)}}{dt}=\frac{d{Sucrose}_{exudation (i)}}{dt}- \frac{d{Sucrose}_{uptake (i)}}{dt}$$

(Eq. S19)

$$\frac{d{Hexose}_{exudation (i)}}{dt}= S_{symplast-soil (i)}\left( t \right)\cdot P_{hexose \left( i \right)}\left( t \right)\cdot\left( {\left[ Hexose \right]_{symplast(i)} \left( t \right)- \left[ Hexose \right]}_{soil (i)} (t) \right)$$

(Eq. S20)

$$\frac{d{Sucrose}_{exudation (i)}}{dt}= S_{phloem-soil (i)}\left( t \right)\cdot P_{sucrose \left( i \right)}\left( t \right)\cdot\left( {\left[ Sucrose \right]_{phloem (i)} \left( t \right)- 0.5\cdot\left[ Hexose \right]}_{soil (i)} (t) \right)$$

(Eq. S21)

$$\frac{d{Hexose}_{uptake (i)}}{dt}= S_{symplast-soil\left( i \right)}\left( t \right)\cdot V_{up. hexose (i)}\cdot\frac{\left[ Hexose \right]_{soil \left( i \right)} \left( t \right)}{K_{up. hexose}+ \left[ Hexose \right]_{soil \left( i \right)} \left( t \right)}$$

(Eq. S22)

$$\frac{d{Sucrose}_{uptake (i)}}{dt}= S_{phloem-soil\left( i \right)}(t)\cdot{0.5\cdot V}_{up. hexose (i)}\cdot\frac{\left[ Hexose \right]_{soil (i)} (t)}{K_{up. hexose}+ \left[ Hexose \right]_{soil (i)} (t)}$$

(Eq. S23)

where *Hexose _net efflux (i)_* (*t*) (moles of hexose) and *Sucrose _net efflux (i)_* (*t*) are the net amount of hexose and sucrose transported outside the root element *i*, respectively, *S _symplast-soil (i)_* (*t*) is the accessible surface of exchange between the soil solution and the root symplast of non-vascular cells in the element *i* (in square meter), *S _phloem-soil (i)_* (*t*) is the accessible surface of exchange between the soil solution and the phloem vessels in the element *i* (in square meter), *P_hexose (i)_* (*t*) and *P_sucrose (i)_* (*t*) are the permeability coefficient of non-vascular root cells’ membrane for hexose and phloem vessels’ membrane for sucrose (in grams of dry structural mass per square meter per second), respectively, *[Hexose]_symplast (i)_* (*t*) is the symplastic concentration of mobile hexose in the root element *i* (moles of hexose per gram of root structural mass), *[Hexose]_soil (i)_* (*t*) is the concentration of hexose at the interface between the root element *i* and the surrounding soil (moles of hexose per gram of root structural mass), *[Sucrose]_phloem (i)_* (*t*) is the concentration of sucrose in the phloem vessels (moles of sucrose per gram of root structural mass), *V_up. hexose_* is the maximal rate of hexose uptake (moles of hexose per square meter per second), and *K _up. hexose_* represents the concentration of hexose in the soil for which the rate of uptake is equal to half of the maximal rate.

We therefore assume that hexose exudation and (re-)uptake occur simultaneously, which enables to cover a range of growth conditions, from situations where re-uptake may be neglected (e.g. in cases where soil microbial degradation of hexose is fast or where the exuded hexose is rapidly transported away from the root surface) to experiments in which re-uptake becomes predominant (e.g. situations of long static hydroponic solutions or experiments in which roots are fed with external glucose). Note that no extra-cost is considered here, i.e. the energy necessary for taking hexose up into the root is assumed to derived from maintenance respiration (see above). The representations of exudation by a pure diffusion process and of uptake by a single Michaelis-Menten kinetics were identical to the formalisms used by Personeni et al. (2007), although the uptake of hexose has also been described using two Michaelis-Menten functions, depending on the range of external sugar concentration (Xia & Saglio, 1988).

The rates of sugar exudation and uptake not only depend on the concentrations of sugars insides and outside root cells, but also on the accessible surface of exchange between the soil solution and the root cells in contact with the soil solution (see Fig. S2). This surface of exchange is dynamic and can decrease because of the formation of transport barriers, i.e. the endodermis that is rapidly formed at the junction between root cortex and root stele and the exodermis that is later formed in certain species at the junction between root epidermis and root cortex (Enstone *et al.*, 2002). These barriers progressively appear along the root, and reduce or even block the free passage of the soil solution through root cortex and/or root stele up to the center of the root cylinder. Hence, the surface of exchange between soil solution and root cells may include all the root cells from epidermis, cortex and stele (including phloem vessels) in very young root elements close to the root tip, or be restricted to epidermis only in old parts of the root axis. Note that both the endodermis and exodermis barriers of a mother root may be temporarily disabled by the emergence of a lateral root, which disturbs the continuity of the barriers until the lateral root is mature enough to form its own barrier. The surfaces of exchange between root cells and soil solution are expressed as:

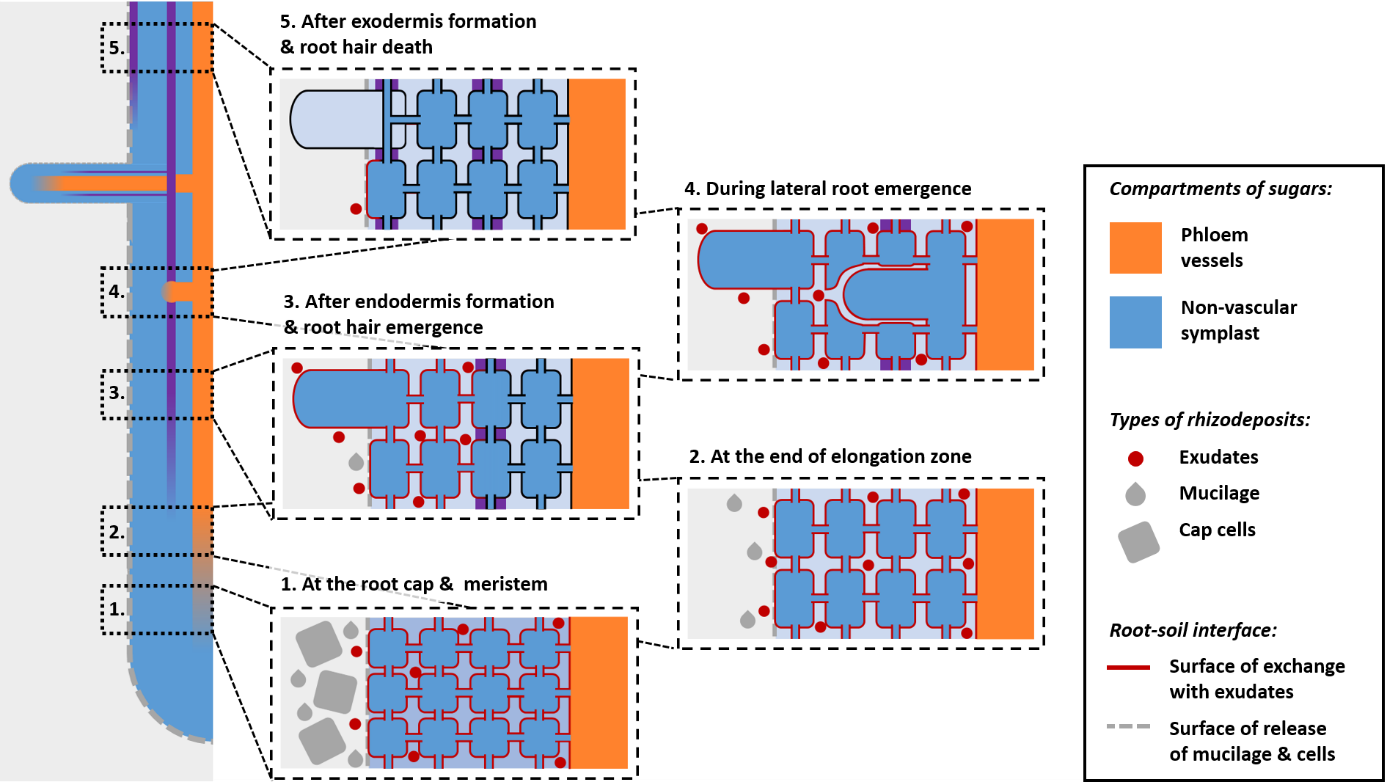

***Fig. S2: Representation of possible spatial variations of rhizodeposition along the root.*** *At the root meristem (1), mucilage and cap cells are released from the external surface of root, while exudates are released from the whole surface of plasma membrane developed by non-vascular cells and by phloem vessels. The release of cap cells progressively decreases and becomes nil at the end of the elongation zone (2). When the endodermis barrier (in purple) is formed between the stele and the cortex (3), the release of exudates is restricted to the plasma membrane of cortical and epidermal cells, including living root hairs. In case of the emergence of a lateral root (4), the barrier of endodermis is disrupted and exudates can again be released from cells within the stele. When the exodermis barrier is formed (5), the surface of release for exudates can be restricted to the external surface of living epidermal cells only.*

$$S_{symplast-soil\left( i \right)}\left( t \right)=S_{epidermis \left( i \right)}\left( t \right)+{Cond}_{walls \left( i \right)}\left( t \right)\times\left( {Cond}_{exodermis \left( i \right)}{\left( t \right)\cdot S}_{cortex \left( i \right)}\left( t \right)+{Cond}_{endodermis \left( i \right)}\left( t \right){\cdot S}_{stele \left( i \right)}\left( t \right) \right)$$

(Eq. S24)

$$S_{phloem-soil\left( i \right)}\left( t \right)={Cond}_{walls \left( i \right)}\left( t \right)\cdot{Cond}_{exodermis \left( i \right)}\left( t \right)\cdot{Cond}_{endodermis \left( i \right)}\left( t \right)\cdot S_{phloem \left( i \right)}\left( t \right)$$

(Eq. S25)

where *S _epidermis (i)_* (*t*), *S _cortex (i)_* (*t*), *S _stele (i)_* (*t*) and *S _phloem (i)_* (*t*) are the actual surface of epidermal cells, cortical cells, non-vascular cells from the stele, and phloem vessels of the root element *i*, respectively (m^2^), and *Cond _walls (i)_* (*t*), *Cond _exodermis (i)_* (*t*), and *Cond _endodermis (i)_* (*t*) are the relative conductance (between 0 and 1) of cell walls, exodermis and endodermis of the root element *i*, respectively. These relative conductance are assumed to evolve with the age of a given root element, with *Cond _walls (i)_* linearly increasing from 0.5 to 1 at very early stages to mimic the increase in cell walls permeability as root cells elongate and mature (Enstone & Peterson, 1992), *Cond _endodermis (i)_* linearly decreasing from 1 to 0 after a few days to mimic the formation of the endodermis, and *Cond _exodermis (i)_* decreasing from 1 to 0 after several days to mimic the formation of the exodermis. Because these conductances evolve continuously between 0 and 1, they relate not only to the total surface of symplast that can be accessed by the soil solution, but also convey an information about kinetic limitation. If we take for example the case where the endodermis formation is only half-completed (*Cond _endodermis_* = 0.5), soil solution would theoretically still be able to enter in contact with all cells from the stele by circulating through immature part of the endodermis; however, the average path to reach the cells from the stele has been increased by the apparition of endodermis obstacles, and the rate of exchange with the cells from the cells is therefore assumed to be halved.

The actual of exchange of the cells from the epidermis, cortex and stele were calculated from allometric relationships based on the external surface of the root:

$$S_{epidermis \left( i \right)}\left( t \right)=x_{epidermis}\cdot{2\cdot\pi\cdot r}_{\left( i \right)}\left( t \right)*l_{\left( i \right)}\left( t \right)$$

(Eq. S26)

$$S_{cortex \left( i \right)} \left( t \right)=x_{cortex}\cdot\cdot{2\cdot\pi\cdot r}_{\left( i \right)}\left( t \right)*l_{\left( i \right)}\left( t \right)$$

(Eq. S27)

$$S_{stele \left( i \right)} \left( t \right)=x_{stele}\cdot\cdot{2\cdot\pi\cdot r}_{\left( i \right)}\left( t \right)*l_{\left( i \right)}\left( t \right)$$

(Eq. S28)

$$S_{phloem \left( i \right)} \left( t \right)=x_{phloem}\cdot\cdot{2\cdot\pi\cdot r}_{\left( i \right)}\left( t \right)*l_{\left( i \right)}\left( t \right)$$

(Eq. S29)

where *x_epidermis_*, *x_cortex_*, *x_stele_*, and *x_phloem_* are allometric coefficients (m), and *r_(i)_ (t)* and l*_(i)_ (t)* are the radius and length of the root element *i* at time *t*.

#### 1.8 Mucilage secretion

The secretion of mucilage is considered in *RhizoDep* as a unidirectional flux from the root epidermis into the soil, which is maximal at the growing tip of the root and decreases with the distance from the tip until becoming negligible. This relationship with the distance to apex is expressed as:

$${Muc}_{max (i)}\left( t \right)= \frac{{Muc}_{ref}}{\left( 1+ \frac{d_{tip \left( i \right)}\left( t \right)-0.5 . l_{\left( i \right)}\left( t \right)}{r_{apex (i)}} \right)^{\gamma_{secretion}}}$$

(Eq. S30)

where *Muc_max (i)_ (t)* is the maximal possible surfacic rate of mucilage secretion and *Muc_ref_* is the reference rate of mucilage secretion (mol of equivalent-hexose per square meter of root external surface per second), *d_tip (i)_ (t)* and *l _(i)_ (t)* are the distance of the root element *i* to the root tip and the length of the root element *i* (m), *r_apex (i)_* is the initial radius of the apex of the root axis to which the root element *i* belongs (m), and γ*_secretion_* is an empirical adimensional coefficient modulating the decrease of *Muc _(i)_ (t)* with the distance from root tip.

The actual rate of mucilage depends on the external surface of the root (excluding root hairs), and is limited both by the concentration of hexose inside peripheral root cells, and by the surfacic concentration of mucilage accumulated at the root-soil interface:

If *[Mucilage]_soil (i)_ (t)* ≤ *[Mucilage]_max_*:

$$\frac{d{Hexose}_{secretion \left( i \right)}}{dt}= S_{external \left( i \right)}\left( t \right)\cdot{Muc}_{\max\left( i \right)}\left( t \right)\cdot\frac{\left[ Hexose \right]_{symplast \left( i \right)} \left( t \right)}{K_{secretion}+ \left[ Hexose \right]_{symplast \left( i \right)} \left( t \right)}\times\left( 1- \frac{\left[ Mucilage \right]_{soil (i)} \left( t \right)}{\left[ Mucilage \right]_{max}} \right)$$

If *[Mucilage]_soil (i)_ (t)* > *[Mucilage]_max_*:

$$\frac{d{Hexose}_{secretion (i)}}{dt}=0$$

(Eq. S31)

where *Hexose _secretion (i)_* (*t*) (moles of hexose) is the amount of hexose converted into mucilage and transported outside the root element *i*, *S _external (i)_* (*t*) is the external surface of the root (in square meter) without root hairs, *[Hexose]_symplast (i)_ (t)* is the concentration of mobile hexose inside the non-vascular root cells (moles of hexose per gram of root structural mass), *K _secretion_* represents the concentration of hexose in the root for which the rate of secretion is equal to half of the maximal rate, and *[Mucilage]_soil (i)_ (t)* and *[Mucilage]_max_* are the actual and maximal surfacic concentration of mucilage at the root-soil interface (moles of equivalent-hexose per square meter of external surface), respectively.

The last term of the equation assumes that the rate of mucilage secretion linearly decreases with the accumulation of mucilage around the root, up to becoming zero when the resulting surfacic concentration of mucilage becomes equal or higher than the maximal prescribed surfacic concentration. This spatial pattern of mucilage secretion has been suggested by several observations along root tips (Paull & Jones, 1975; Werker & Kislev, 1978; Pankievicz *et al.*, 2022). The formalism used in *Eq. S30* has been adapted from the one suggested by Personeni *et al.* (2007) for describing the decrease of sugar exudation with the distance from the tip, although no experimental data is currently available to prove the relevance of such formalism in the specific case of mucilage secretion.

We chose to exclude root hairs from the calculation of the external surface involved in the secretion of mucilage, as to our knowledge, no study has been able to demonstrate that root hairs are actively involved in the secretion of mucilage (Peterson & Farquhar, 1996). While the process of mucilage secretion appears to be linked to the vesicular transport of pectic polysaccharides derived from starch in the cap cells (Morré *et al.*, 1967; Paull & Jones, 1976b), we considered that this starch does not belong to the reserve pool of sugar represented in *RhizoDep*, as it has been necessarily synthesized very recently in cap cells (Paull & Jones, 1976b). Hence, we represented a dependency of mucilage secretion to the pool of mobile sugars in the root, not the reserve pool. The regulation of mucilage secretion by the availability of root hexose introduced in *Eq. 31* is justified by a few studies that suggested that the availability of sugars (e.g. sucrose, hexose) to the root cap cells controls the amount of mucilage secretion (Jones & Morré, 1973; Paull *et al.*, 1975; Iijima *et al.*, 2003). Conversely, a limitation of the secretion rate by the accumulation of mucilage outside the root cell membranes has been shown by some authors, e.g. when there is not enough water to enable the diffusion of the polysaccharides away from the root (Morré *et al.*, 1967; Paull & Jones, 1976a). We did not introduce a direct dependency of the rate of mucilage secretion with the rate of root elongation, as the release of mucilage was shown to be independent of root growth rate (Jones & Morré, 1973), even if such a correlation has been suggested by others (Iijima *et al.*, 2003). Note however that in *RhizoDep*, the secretion of mucilage along a root axis is stopped once the axis has stopped elongating.

#### 1.9 Root cap cells release

The release of root cap cells has been described in *RhizoDep* as a unidirectional flux of matter from the root cap cells surface into the soil, which linearly decreases with the distance from the root tip and becomes zero above the elongation zone:

$${Sloughing}_{\left( i \right)}\left( t \right)= {Sloughing}_{ref} \cdot\frac{l_{elongation \left( i \right)}-\left( d_{tip \left( i \right)}\left( t \right)-0.5\cdot l_{\left( i \right)}\left( t \right) \right)}{l_{elongation \left( i \right)}}$$

(Eq. S32)

where *Sloughing _(i)_ (t)* is the maximal possible surfacic rate of cap cells release and *Sloughing _ref_* is the reference rate of cells release (mol of equivalent-hexose per square meter of root external surface per second), *l _elongation (i)_* is the length of the elongation zone of the root axis to which the root element *i* belongs (meter), and *d_tip (i)_ (t)* and *l_(i)_ (t)* are the distance of the root element *i* to the root tip and the length of the root element *i* (meter).

The actual rate of cells release depends on the external surface of the root element, and is limited by the surfacic concentration of root cap cells accumulated at the root-soil interface:

If *[Cells]_soil (i)_ (t)* ≤ *[Cells]_max_*:

$$\frac{d{Hexose}_{sloughing \left( i \right)}}{dt}= S_{external \left( i \right)}\left( t \right) \times{Sloughing}_{\left( i \right)}\left( t \right)\times\left( 1- \frac{\left[ Cells \right]_{soil} \left( t \right)}{\left[ Cells \right]_{max}} \right)$$

If *[Cells]_soil (i)_ (t)* > *[Cells]_max_*:

$$\frac{d{Hexose}_{sloughing (i)}}{dt}=0$$

(Eq. S33)

where *Hexose _sloughing (i)_* (*t*) (moles of hexose) is the equivalent amount of hexose converted into the production of cap cells that are released over time for the root element *i*, *S _external (i)_* (*t*) is the external surface of the root (in square meter) without root hairs, and *[Cells]_soil (i)_ (t)* and *[Cells]_max_* are the actual and maximal surfacic concentration of cells at the root-soil interface (moles of equivalent-hexose per square meter of external surface), respectively.

The dependency of root cap cell release with the distance from the tip introduced in *Eq. S32* illustrates the fact that the detachment of cells is only possible within the root cap, which only covers a short length of the root tip. We did not introduce an explicit parameter for calculating the exact length of the extension of the cap along the root, and instead assumed that no cell release should be allowed above the root elongation zone. As we did not find any quantitative information regarding the distribution of cap cells release along the root, we assumed that the amount of C lost as cap cells linearly decreases along the root, since cap cells become thinner as the cap extends further away from the root tip and are probably less frequently sloughed off.

We did not introduce a direct dependency of the rate of cap cells release with the rate of root elongation, as the mitotic activity of the cap, responsible for the release of cap cells, was shown to be independent on the root elongation rate (Brigham *et al.*, 1998; Curlango-Rivera *et al.*, 2010). However, as done for mucilage secretion, the release of cells is prevented in the model once the root axis has stopped elongating.

As cap cells release was also shown to be independent from external applications of sugars to pea root tips (Curlango-Rivera *et al.*, 2010), we further assumed that the rate of release was not regulated by the amount of root available sugars, unlike the processes of root exudation and mucilage secretion. However, similarly to the process of mucilage secretion, we introduced in *Eq. S33* a limitation of the release of cells by the accumulation of cap cells at the root-soil interface, as it was shown that conditions where cap cells are prevented from moving away from the root (e.g. when no water is available) also inhibit the synthesis of new cap cells (Brigham *et al.*, 1998).

Note that the process of cells release described here excludes the contribution of root hairs sloughing, which was not introduced in *RhizoDep* considering the absence of evidence that dead root hairs are in fact sloughed off over time (Fusseder, 1987; Nguyen, 2003).

#### 1.10 Rhizodeposits degradation in the rhizosphere

A simple function has been implemented in *RhizoDep* to describe the disappearance of rhizodeposits at the soil-root interface, which can be considered as their consumption/transformation by microorganisms in the rhizosphere, their immobilization by sorption on soil particles, or the transport away from the root. For the sake of simplification, we introduced only one unique function based on a Michaelis-Menten formalism for each type of rhizodeposit:

$$\frac{d{Hexose}_{degradation (i)}}{dt}= S_{exchange \left( i \right)}\left( t \right)\times V_{deg. hexose} . \frac{\left[ Hexose \right]_{soil (i)} (t)}{K_{deg. hexose}+ \left[ Hexose \right]_{soil (i)} (t)}$$

(Eq. S34)

$$\frac{d{Mucilage}_{degradation (i)}}{dt}= S_{external (i)}(t) . V_{deg. mucilage} . \frac{\left[ Mucilage \right]_{soil (i)} (t)}{K_{deg. mucilage}+ \left[ Mucilage \right]_{soil (i)} (t)}$$

(Eq. S35)

$$\frac{d{Cells}_{degradation (i)}}{dt}= S_{external (i)}(t) . V_{deg. cells} . \frac{\left[ Cells \right]_{soil (i)} (t)}{K_{deg. cells}+ \left[ Cells \right]_{soil (i)} (t)}$$

(Eq. S36)

where *Hexose _degradation (i)_* (moles of hexose), *Mucilage _degradation (i)_* (moles of equivalent-hexose) and *Cells _degradation (i)_* (moles of equivalent-hexose) correspond to the amount of hexose, mucilage and cap cells degraded (or made unavailable) at the root-soil interface of element *i*, respectively, *S _exchange (i)_* is the accessible surface of exchange between the soil solution and the root (sum of *S _symplast-soil (i)_* and *S _phloem-soil (i)_*) of the root element *i* (in square meter), *S _external (i)_* is the external surface of the root element i (excluding root hairs), *V_deg. hexose_*, *V_deg. mucilage_*, and *V_deg. cells_* are the maximal rate of degradation of hexose, mucilage and cells (moles of hexose per square meter per second), respectively, *[Hexose]_soil (i)_* is the mass concentration of hexose at the interface between the root element *i* and the surrounding soil (moles of hexose per gram of root structural mass), *[Mucilage]_soil (i)_* and *[Cells]_soil (i)_* are the surfacic concentration of mucilage and cells on the external surface of the root element *i* (moles of equivalent-hexose per square meter), and *K_deg. hexose_, K_deg. mucilage_*, and *K_deg. cells_* represent the concentration of hexose, mucilage and cells for which the rate of degradation is equal to half of the maximal rate, respectively.

*Eq. S34* reuses a similar Michaelis-Menten formalism at the one used for (re-)uptake of hexose by the root (see above), which enables to easily predict the range of concentrations of hexose in the soil for which degradation may dominate the fate of hexose at the soil-root interface, compared to the possibility of re-uptake.

Using *Eq. S34-S36* to describe the disappearance of rhizodeposits from the root-soil interface has been made necessary in this work in the absence of an actual soil model able to predict the degradation of distinct rhizodeposits in the soil. In the future, we expect that functions corresponding to first-order kinetics (microbial degradation) or diffusion/advection (movement of rhizodeposits away from the roots) would be managed by a soil model coupled to *RhizoDep*, therefore avoiding the need of using *Eq. S33-S35*.

#### 1.11 Carbon balance

A local C balance is eventually made, so that the change in the amount of sugars and rhizodeposits in each root element *i* can be calculated according to all the rates previously calculated:

$$\frac{d{Sucrose}_{phloem (i)}}{dt}=\frac{d{Sucrose}_{production (i)}}{dt}- \frac{1}{2} \cdot\frac{d{Hexose}_{production \left( i \right)}}{dt}+ \frac{d{Sucrose}_{allocation \left( i \right)}}{dt}- \frac{d{Sucrose}_{exudation (i)}}{dt}- \frac{d{Sucrose}_{uptake (i)}}{dt}$$

(Eq. S37)

$$\frac{d{Hexose}_{symplast\left( i \right)}}{dt}=\frac{d{Hexose}_{production \left( i \right)}}{dt}- 2 \cdot\frac{d{Sucrose}_{production \left( i \right)}}{dt}- \frac{d{Hexose}_{growth \left( i \right)}}{dt}- \frac{d{Hexose}_{maintenance \left( i \right)}}{dt}- \frac{d{Hexose}_{immobilization \left( i \right)}}{dt}+ \frac{d{Hexose}_{mobilization \left( i \right)}}{dt}- \frac{d{Hexose}_{exudation \left( i \right)}}{dt}+ \frac{d{Hexose}_{uptake \left( i \right)}}{dt} - \frac{d{Hexose}_{secretion \left( i \right)}}{dt}- \frac{d{Hexose}_{sloughing \left( i \right)}}{dt}$$

(Eq. S38)

$$\frac{d{Hexose}_{reserve (i)}}{dt}= \frac{d{Hexose}_{immobilization \left( i \right)}}{dt}- \frac{d{Hexose}_{mobilization \left( i \right)}}{dt}$$

(Eq. S39)

$$\frac{d{Hexose}_{soil (i)}}{dt}=\frac{d{Hexose}_{exudation \left( i \right)}}{dt}- \frac{d{Hexose}_{uptake \left( i \right)}}{dt}+ 2 \left( \frac{d{Sucrose}_{exudation \left( i \right)}}{dt}- \frac{d{Sucrose}_{uptake \left( i \right)}}{dt} \right)-\frac{d{Hexose}_{degradation (i)}}{dt}$$

(Eq. S40)

$$\frac{d{Mucilage}_{soil (i)}}{dt}=\frac{d{Mucilage}_{secretion \left( i \right)}}{dt}-\frac{d{Mucilage}_{degradation (i)}}{dt}$$

(Eq. S41)

$$\frac{d{Cells}_{soil (i)}}{dt}=\frac{d{Cells}_{sloughing \left( i \right)}}{dt}-\frac{d{Cells}_{degradation (i)}}{dt}$$

(Eq. S42)

where *Sucrose _phloem (i)_*, *Hexose _symplast (i)_*, *Hexose _reserve (i)_*, *Hexose _soil (i)_*, *Mucilage _soil (i)_*, and *Cells _soil (i)_* represent for the root element *i* the amounts of sucrose in the phloem vessels (moles of sucrose), of mobile hexose in non-vascular cells (moles of hexose), of sugars in the reserve pool (moles of equivalent-hexose), and of hexose, mucilage and cap cells at the root-soil interface (moles of (equivalent-)hexose), respectively.

### 2 Temperature adjustments in *RhizoDep*

*RhizoDep* considers that any growth or metabolic process *F* can be ascribed to a common law of response to soil temperature *T*, described as:

$$F\left( T \right)=F\left( T_{ref} \right) . \left[ A . \left( T-T_{ref} \right)+B \right]^{1-C} . \left[ A . \left( T-T_{ref} \right)+B \right]^{\frac{C . \left( T-T_{ref} \right)}{10}}$$

(Eq. S43)

where F(*T*) is the temperature-dependent process at temperature *T* (in Kelvin), *F*(*T_ref_*) is the value of the process at reference temperature *T_ref_*, *A* and *B* are empirical constants, and *C* is a control parameter equal to 0 or 1.

We have designed this law as a versatile general rule, where four contrasted behaviors can be expected, depending on the values of the coefficients A, B and C:

- *Case 1:* if C=0, A=0 and B=1: the process does not depend on soil temperature,
- *Case 2:* if C=0 and A>0: the relationship is equivalent to a linear increase of the process value with temperature, on which the calculation of growing degree days is usually based,
- *Case 3:* if C=1, A=0 and B>0: the relationship is equivalent to the commonly-used *Q10* function, which predicts an exponential growth of the process value with temperature, assuming that the Q10 value, which corresponds to the relative increase of the process rate for each 10-degree increase in temperature, is independent of temperature,
- *Case 4:* if C=1, A<0 and B>0: the relationship is equivalent to a bell-shaped curve that is similar to the *Q10* function at low temperature but reaches a maximum at intermediate temperature and then decreases at higher temperature.

The latter case is similar to the case described by Parent et al. (2010), which can be conceptually thought as an increase in enzymatic activity with low temperatures, coupled to the denaturation of the considered enzyme at high temperatures. Using the modified Arrhenius law proposed by Johnson et al. (1942), Parent et al. (2010) have shown that such behavior can be widely observed for a range of plant metabolic processes, even those that are not controlled by the activity of a single enzyme. Although the parameters used in the modified Arrhenius law had an original meaning in terms of enthalpy or entropy of activation/deactivation, such parameters are not easy to determine nor to interpret. Hence, we have preferred to use a slightly simpler equation that corresponds to a modified *Q10* function. This equation is based on the work of Tjoelker et al. (2001), who suggested that the *Q10* value, commonly assumed to be independent on temperature, is actually decreasing with temperature in an almost linear way.

We acknowledge that *Eq. S43* does not cover all possible cases, such as a modification of sugar transport by the change in temperature-dependent viscosity, or the damage of root tissue at low (frost) temperature. However, we consider *Eq. S43* to be satisfying enough for describing a majority of situations where different processes follow different responses to temperature. The temperature-dependence is represented either as a direct modification of the rate over the simulation following *Eq. S42*, or as a modification of the “physiological time” experienced by the root, using a temperature-compensated rate (Parent *et al.*, 2010) and a “thermal time” calculated using *Eq. S43*. In such case, the process has to be parametrize at a known reference temperature, while the time used in the corresponding equation is not the actual time but is calculated as:

$$t_{thermal}\left( t, T \right)=t*\frac{F(T)}{F(T_{ref})}$$

(Eq. S44)

where $t_{thermal}\left( t, T \right)$ is the temperature-adjusted time (in second) depending on the actual time (in second) and temperature *T* (in Kelvin), and the ratio of *F(T)* with *F(T_ref_)* represents the temperature-modifier coefficient and is calculated using *Eq. S42*.

For example, the maximal elongation of a root axis *Δl* over a period *Δt* (see *Eq. S4*) is calculated by multiplying the maximal rate of elongation *elongation_max_*, which has been defined for a temperature *T_ref_* of 20°C, by the actual time step *Δt* and by the temperature-modifier coefficient obtained from the case 2 of *Eq. S43* with *F(T_ref_)* = 1, A = 1/*T_ref_*, B = 0 and C = 0. In this case, the calculation is equivalent to the classical expression of growth according to the number of degree-days commonly used in crop models.

In *RhizoDep*, we assume that all growth-related processes (e.g. elongation, thickening, and aging until root growth cessation or until death) obey to the Case 2 of *Eq. S42*, while other processes obey to Case 3 or 4, and some are considered independent from temperature. The detailed parametrization of each process for temperature adjustment is presented in Excel file Ex1.

### 3 Modularity of *RhizoDep*

*RhizoDep* has been organized in several modules, which are successively called over a given time step (Fig. S3).

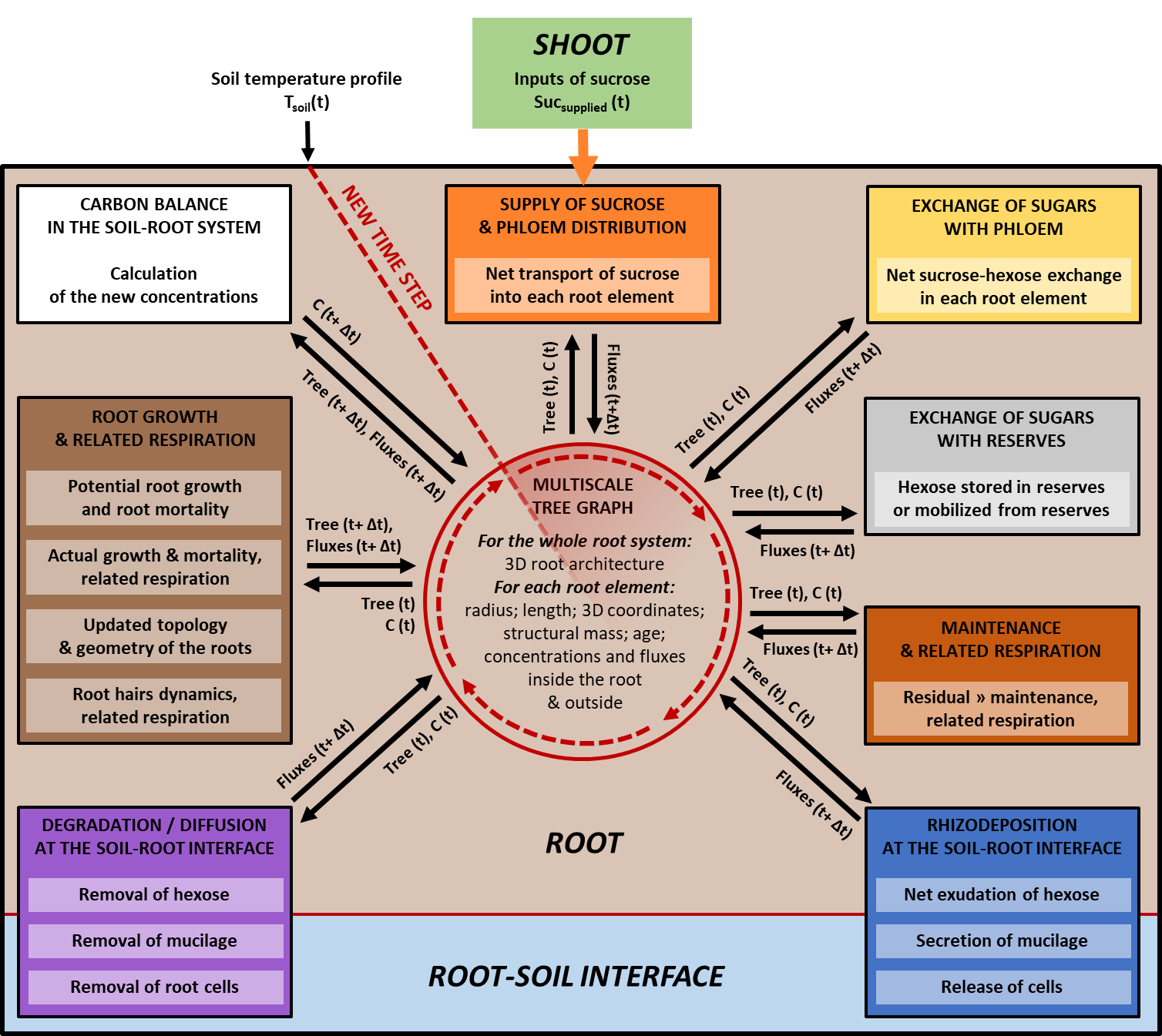

***Fig. S3: Modular organization and exchange of variables within the model RhizoDep.*** *For every new time step, the previous values of state variables stored in the Multiscale Tree Graph at time t are read by each module and used to compute the new variables related to the growth or metabolic rates at t + Δt. Tree(t) represent all variables linked to the topology and geometry of the root system, while C(t) represent the concentrations of C in any pool. The transfer of sucrose from the shoots to the root system is the main input variable of the system.*

### 4 Recalculating belowground C flows in the study of Swinnen *et al.* (1994)

We summarize here the main steps of the calculations of plant C flows from the data provided by Swinnen et al. (1994b) following the field measurement on spring wheat done in 1989. Details about the experiment can be found in Swinnen et al. (1994a, b).

#### 4.1 Calculation of the fractions of the net C fixed allocated to different processes

The allocation of C to the production of shoot biomass (%) was derived from the experimental data provided in Figure 1 by Swinnen et al. (1994b) using a new Gompertz function. We fitted the experimental data by specifying an asymptote at 100% and by adding an extra theoretical point (allocation = 1% at 70 Julian days) so that the function works also at early stages:

$${Alloc}_{shoot biomass}\left( t \right)=100\times\exp\left( -exp\left( 3.34381 - 0.0266925\times t) \right) \right)$$

(Eq. S45)

where *Alloc_shoot biomass_ (t)* is the fraction of C allocated to shoot biomass (%) and *t* is time expressed in Julian days.

Using this new curve, the experimental data from Figure 1 was slightly modified so that it takes into account an increase towards 100% of the proportion of C allocated to shoot biomass production. We used the new *Alloc_shoot_ (t)* and modified the experimental points of the allocation to respiration or soil at 151, 165, 179, 193 and 207 days proportionally to the ratio between the new proportion of C to roots (estimated by difference between 100% and *Alloc_shoot_ (t)*) and the former one used by Swinnen et al. (1994b) (Table S1).

The allocation of C to belowground respiration (%), which includes both the CO_2_ respired by roots and the CO_2_ originating from the mineralization of rhizodeposits, was derived from the experimental data provided in Figure 1 in Swinnen et al. (1994b) using a new exponential function. We fitted the readjusted experimental data on C allocation by adding an extra theoretical point (allocation = 60% at 60 Julian days) so that the function generates plausible value for early growth stages as well:

$${Alloc}_{below resp}\left( t \right)=181.453\times\exp\left( -0.0167158\times t \right)$$

(Eq. S46)

where *Alloc_below resp_ (t)* is the fraction of C allocated to belowground respiration (%) and *t* is time expressed in Julian days.

We assumed that 18% of total respiration originated from rhizodeposit mineralization, which corresponds to the mean value between the two extremes suggested by Swinnen et al. (1994b) (11% for root cell walls material and 25% for glucose), and, by complementarity, we assumed that 82% of total respiration originated from root respiration :

${Alloc}_{rhizodep. resp}\left( t \right)=0.18\times{Alloc}_{below resp}\left( t \right)$

*(Eq. S47)*

${Alloc}_{root resp}\left( t \right)=0.82\times{Alloc}_{below resp}\left( t \right)$

*(Eq. S48)*

where *Alloc_resp_ (t)*, *Alloc_rhizodep. resp_ (t)* and *Alloc_root resp_ (t)* are the fraction of C (%),allocated to belowground respiration rhizodeposits mineralization and root respiration, respectively and *t* is time expressed in Julian days.

The net allocation of C to the soil (%), which includes the rhizodeposits transferred to soil but excludes the part of the rhizodeposits that has been mineralized as CO_2_ by microorganisms, was derived from the experimental data provided in Figure 1 in Swinnen et al. (1994b) using a new linear function.

We fitted the (readjusted) experimental data on C allocation by adding an extra theoretical point (allocation = 0% at 240 Julian days) so that the function generates plausible value at late maturity:

${Alloc}_{net soil}\left( t \right)=-0.0468870\times t + 12.3941$

*(Eq. S49)*

where *Alloc_net soil_ (t)* is the fraction of C allocated to the net accumulation of rhizodeposits in the soil (%) and *t* is time expressed in Julian days.

The allocation of C to root biomass production (%) was calculated as the difference between 100% and the other allocation coefficients calculated above, based on the total C balance:

$${Alloc}_{root biomass}\left( t \right)=100-\left( {Alloc}_{shoot biomass}+{Alloc}_{below resp}+{Alloc}_{net soil} \right)$$

*(Eq. S50)*

where *Alloc_root biomass_ (t)* is the fraction of C allocated to the root biomass (%) and *t* is time expressed in Julian days.

**Table S1: Comparison of the coefficient of C allocation allocated to shoot biomass production, belowground respiration or net soil C accumulation between the study of Swinnen et al. (1994b) (*Original*) and our recalculations**

|  | C allocation to shoot biomass production (%) | | C allocation to belowground respiration (%) | | C allocation to net soil C accumulation (%) | |
| --- | --- | --- | --- | --- | --- | --- |
| Julian days | *Original* | Recalculated | *Original* | Recalculated | *Original* | Recalculated |
| 60 | *NA* | NA | *NA* | *60* | *NA* | NA |
| 151 | *62.11* | 60.46 | *15.97* | 16.66 | *4.47* | 4.66 |
| 165 | *73.18* | 70.73 | *11.61* | 12.67 | *3.44* | 3.75 |
| 179 | *80.32* | 78.8 | *10.12* | 10.90 | *4.07* | 4.38 |
| 193 | *82.01* | 84.87 | *7.73* | 6.49 | *6.21* | 5.22 |
| 207 | *79.76* | 89.33 | *8.81* | 4.65 | *5.94* | 3.13 |
| 240 | *NA* | NA | *NA* | NA | *NA* | *0* |

#### 4.2 Calculation of the evolution of the amount C in shoot biomass over time

The evolution of shoot biomass over time was calculated by reusing the same Gompertz function as used by Swinnen et al. (1994b) and shown in Figure 3:

$$C_{shoot biomass}\left( t \right)=6083\times\exp\left( -673\times\exp\left( -0.0388\times t \right) \right)$$

(Eq. S51)

where *C_shoot biomass_ (t)* is the shoot biomass C (kgC ha^-1^) and *t* is time expressed in Julian days.

#### 4.3 Calculation of the daily flows of C allocated to different processes

We calculated the net amount of C fixed by photosynthesis using the shoot C content evolution and the allocation to shoot biomass calculated as explained above.

The final net amount of C fixed by photosynthesis every day (corresponding to the gross primary production minus the shoot respiration) was estimated from the evolution of shoot C content over time and the allocation of C to shoot production determined previously:

$$R_{fixed} \left( t \right)=\frac{C_{shoot}\left( t \right)- C_{shoot}\left( t-1 \right)}{{{Alloc}_{shoot biomass}\left( t \right)}/{100}}$$

(Eq. S52)

where *R_fixed_* *(t)* is the net amount of C fixed on day *t* (kgC ha^-1^ day^-1^) and *t* is the time expressed in Julian days.

We then used the C allocation coefficients (%) and *R_fixed_ (t)* (kgC ha^-1^ day^-1^) to calculate the other daily flows of aboveground and belowground plant C:

$$R_{shoot biomass} \left( t \right)=\frac{{Alloc}_{shoot biomass}\left( t \right)}{100}\times C_{fixed} \left( t \right)$$

(Eq. S53)

$$R_{belowground} \left( t \right)=\frac{100- {Alloc}_{shoot biomass}\left( t \right)}{100}\times C_{fixed} \left( t \right)$$

(Eq. S54)

$$R_{root biomass} \left( t \right)=\frac{{Alloc}_{root biomass}\left( t \right)}{100}\times C_{fixed} \left( t \right)$$

(Eq. S55)

$$R_{root resp} \left( t \right)=\frac{{Alloc}_{root resp}\left( t \right)}{100}\times C_{fixed} \left( t \right)$$

(Eq. S56)

$$R_{rhizodeposition} \left( t \right)=\frac{{Alloc}_{net soil}\left( t \right)+{Alloc}_{rhizodep. resp}\left( t \right)}{100}\times C_{fixed} \left( t \right)$$

(Eq. S57)

where *R_shoot biomass_(t)*, *R_belowground_* *(t)*, *R_root biomass_* *(t)*, *R_root resp_* *(t)* and *R_rhizodeposition_* *(t)* are the net amount of C used on day *t* (kgC ha^-1^ day^-1^) to produce shoot biomass, to transit to roots, to produce root biomass, to generate root respiration and to generate rhizodeposits, respectively.

#### 4.4 Calculation of the evolution of root biomass C and root necromass C over time

In the study of Swinnen et al. (1994b), the flow of C entering the soil corresponding to root mortality was calculated using a C balance on the estimated allocation of C to shoot biomass production and the estimated curve of net root biomass; however, this balance led to an odd behavior, with a first peak of root decay occurring very early at the end of May and a second one occurring in July. Here, the evolution of root C content (kgC ha^-1^) over time was recalculated using either the plausible evolution of shoot:root ratio and the evolution of shoot C content calculated previously, or a C balance based on the principle that no significant root decay should occur at early growth stages.

From Day 62 to Day 130, the root C content was calculated based on the daily amount of C used for root biomass production calculated above, assuming that no significant root decay would occur over that period:

$$C_{root biomass}\left( t \right)=C_{root biomass}(t-1)+R_{root biomass} \left( t \right)$$

(Eq. S58)

where *C_root biomass_ (t)* and *C_root biomass_ (t-1)* represent the amount of C in the root biomass (kg_C_ ha^-1^) at day *t* and *t-1*, respectively, *R_root biomass_* *(t)* (kg_C_ ha^-1^ day^-1^) is the net amount of C used over day *t* to produce root biomass, and *t* is time expressed in Julian days.

From Day 131 onwards, we recalculated the shoot:root ratio (kg_C_ kg_C_^-1^) with the empirical Gompertz function suggested by Swinnen et al. (1994b):

$$Shoot:Root \left( t \right)=15.2\times\exp\left( -24.7\times\exp\left( -0.0171\times t \right) \right)$$

(Eq. S59)

where *Shoot:Root (t)* is the shoot-to-root biomass (kg_C_ kg_C_^-1^) and *t* is time expressed in Julian days.

We then recalculated the corresponding root C content (kg_C_ ha^-1^) based on the evolution of the shoot C content:

$$C_{root biomass}\left( t \right)=\frac{C_{shoot biomass} \left( t \right)}{Shoot:Root \left( t \right)}$$

(Eq. S60)

Over the whole monitoring period, the amount of dead roots (kg_C_ ha^-1^) was eventually calculated based on C balance, as suggested by Swinnen et al. (1994b):

$$C_{root necromass}\left( t \right)=R_{root biomass}\left( t \right)-\left( C_{root biomass} \left( t \right)- C_{root biomass} \left( t-1 \right) \right)$$

(Eq. S61)

where *C_root necromass_ (t)* represents the amount of C lost as root necromass (kg_C_ ha^-1^) at day *t*, *R_root biomass_* *(t)* (kg_C_ ha^-1^ day^-1^) is the net amount of C used over day *t* to produce root biomass, and *t* is time expressed in Julian days.

#### 4.5 Calculation of the cumulative amounts of C allocated to a specific use

The cumulative amounts of C allocated to a specific use over the whole growth were calculated from the daily amounts *R* calculated above:

$${Cum}_{X}\left( t \right)={Cum}_{X}\left( t-1 \right)+R_{X}(t)$$

(Eq. S62)

where *Cum_X_ (t)* and *Cum_X_ (t-1)* represent the cumulative amount of C (kgC ha^-1^) allocated over time to *X* at day *t* and *t-1,* respectively, and *R_X_* *(t)* (kgC ha^-1^ day^-1^) is the net amount of C used over day *t* to *X*, where *X* represents either shoot or root biomass production, root respiration or rhizodeposition, and *t* is time expressed in Julian days.

#### 4.6 From C per hectare to individual plant C

According to Swinnen et al. (1994b), the estimated plant density after emergence on 18 April was 240 plant m^-2^, i.e. 2.4 10^6^ plants ha^-1^. We assumed that no plants died from that moment up to the harvest, and therefore used this plant density to convert all values in kg_C_ per hectare into g_C_ per plant.

#### 4.7 Determination of soil temperature

No measurements of air or soil temperature was provided in the study of Swinnen et al. (1994a,b). For estimating soil temperature over time, we therefore used another source of information, using the data on soil temperature collected from the same site, but one year later than the study of Swinnen et al. (1994b) (i.e. 1990 instead of 1989). Soil temperature recordings from Lovinkhoeve's farm meteorological station were available in Fig. 5.12 of the PhD thesis manuscript of de Vos (1997). To obtain daily average temperature, we used linear relationships between two recording points so that an acceptable estimation of average temperature could be obtained every day. The resulting estimation of soil temperature evolution is represented in Fig. S4a.

| **a. Soil temperature** | **b. Amount of C allocated to roots** |
| --- | --- |
| 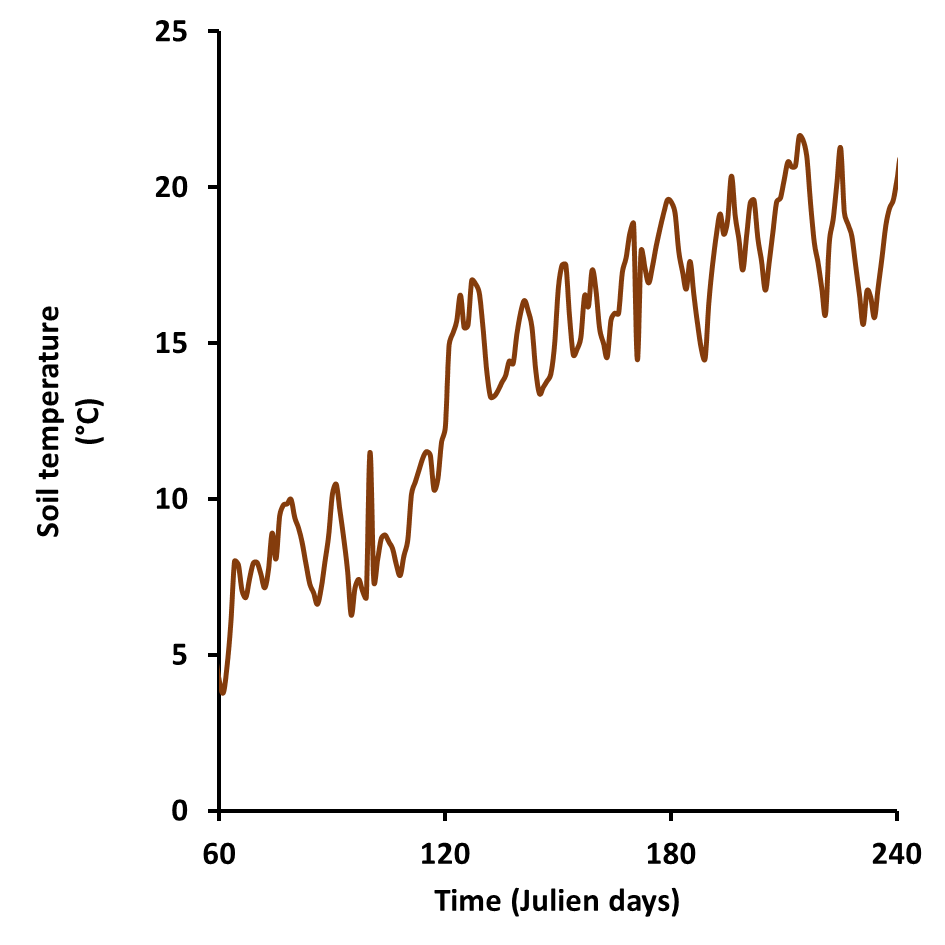 | 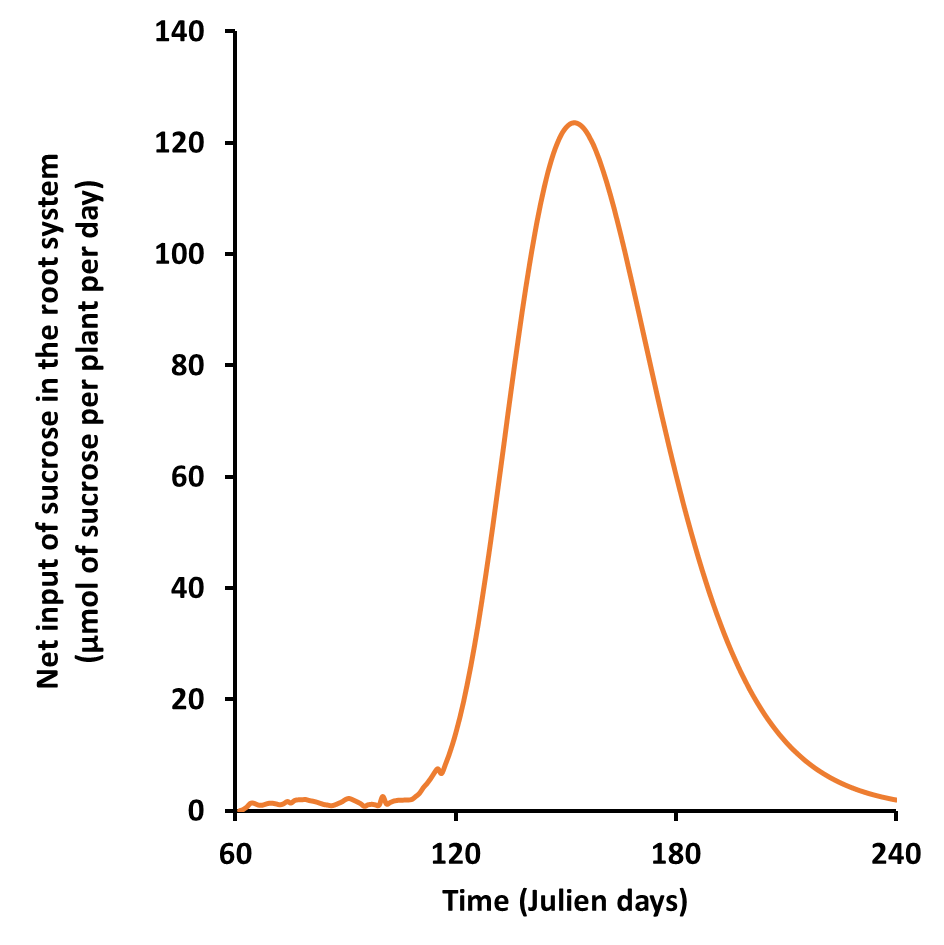 |

***Fig. S4: Evolutions of the soil temperature (a, left) and the net input of C into the root system (b, right) used to simulate the experiment of Swinnen et al. (1995b) done in 1989.*** *Soil temperature was measured on the same site in Lovinkhoeve's farm one year later in 1990. The net input of C to the root system was recalculated by using the data of Swinnen et al. (1995b) and by adding an extra amount of C over the first days corresponding to the mobilization of C from the germinated seed, recalculated from the data of Sun et al. (2020).*

### 5 Recalculation of C supply at germination stage

The measurements and calculations using the study of Swinnen et al. (1994b) were not precise enough to estimate the contribution of the reserve of C mobilized from the endosperm of the wheat seed at germination and early growth stages. For doing this, we used the data reported by Sun et al. (2020) on the evolution of endosperm mass and starch content of the grain during post-germination seedling growth for the wheat genotype RL4452. The calculated amount of starch measured in the germinated grain decreased linearly over the first 7 days following imbibition (R^2^ = 0.93), with a daily loss equivalent to 7.0 10^-11^ mol of equivalent-sucrose per second at 22°C. We therefore applied this constant input of C over the first days of wheat growth in our simulation, after temperature adjustment with the daily soil temperature calculated from the study of Swinnen et al. (1994b) until reaching a cumulative supply of 7.85 10^-5^ moles of supplied sucrose. This early input of sucrose was finally added to the one estimated from the data of Swinnen et al. (1994b), and the sum of the two curves resulted in the overall evolution of the rate of sucrose supply to the root system indicated here in Fig. S4b, and used thereafter as the input variable of *RhizoDep* for simulating wheat growth.

### 6 Estimation of parameters for simulating wheat growth and rhizodeposition

The list of *RhizoDep*’s parameters with their values estimated in the case of wheat growth as well as additional justifications is available in Supporting Tables ST. 133 numerical parameters were used in our simulations (excluding options for designing the simulation or for displaying the results), from which 50 corresponded to temperature-adjustment parameters alone. In particular, 31 parameters were related to root growth and morphology, and 24 controlled the processes of rhizodeposition and hexose uptake from soil. Growth parameters were mostly derived from our own experiments on spring wheat, and completed by literature data when needed. Non-growth parameters were generally derived from literature data, or, when no data was available, were estimated from other parameters (see below).

#### 6.1 Wheat-dependent growth parameters

Wheat root growth parameters were mainly obtained from two unpublished experiments conducted in 2022 (data not shown). In the first experiment, spring wheat (c.v. Lennox) was grown from April to June in a greenhouse in pots (7.0 x 9.8 x 41.5 cm) filled with an agricultural Luvisol sampled from the control experimental plot *QualiAgro* (Kpemoua *et al.*, 2023) and fertilized with 150 kgN.ha^-1^, 25 kgP.ha^-1^, and 75 kgK.ha^-1^. Wheat was grown at an equivalent density of 300 plants m^-2^, with two plants per pot. At flowering stage, 56 days after sowing, roots were sampled, cleaned and scanned at 2400 dpi for determining i) the minimal and maximal root diameters, ii) the inter-branching distance and iii) the diameter-related relationships among different root orders. A complementary experiment was conducted with the same wheat cultivar and soil in a growth chamber at a mean daily temperature of 20°C, using small rhizoboxes (20 cm x 47 cm x 0.5 cm) equipped with a transparent window. Time-lapse images of the root system were taken and were used together with high-resolution scans of the roots after three weeks to estimate: i) root elongation rate as a function of root apical diameter, ii) the delay of root primordium emergence, and iii) the statistical distribution of growth duration.

According to our observation of lateral roots over time in rhizoboxes, the growth duration varied dramatically among the lateral roots emerging from seminal roots, from 0.2 to 6 days at 20°C, with no obvious relationship with their apical diameter - contrary to what is assumed in *ArchiSimple* (Pagès *et al.*, 2014). The prescribed growth duration of any lateral root (root order ≥ 2) was therefore randomly chosen from three possible classes derived from our observations, with distinct probabilities: 0.25 days (55% probability), 0.70 days (30% probability), and 6 days (15% probability). Seminal and adventitious roots were ascribed a unique growth duration ten times higher than the highest one observed for lateral roots, i.e. 60 days.

The coefficient corresponding to life duration (*LDs*) was calculated by assuming that the death of the first seminal root should start at the age of 45 days equivalent to a temperature of 20°C in order to reproduce the onset of root mortality calculated from the data of Swinnen *et al.* By assuming a seminal root of 0.7 mm diameter and of tissue density of 0.14 g cm^-3^, *LDs* was estimated to be 4 10^4^ s m^-1^ g_DW_^-1^ m^3^. We eventually chose a 20% higher value, i.e. 5 10^4^ s m^-1^ g_DW_^-1^ m^3^, as the definitive life duration coefficient, so that most nodal roots will die only a few weeks before harvest.

Root tissue density of wheat root was assumed constant and uniform over the root system and equal to 0.14 g cm-^3^, according to personal measurements on small root segments (1-2 cm) of winter wheat (c.v. Nara) grown in rhizoboxes (data not shown), and according to the study on six Poaceae species made by Pagès & Picon-Cochard (2014), which gave a mean RTD value of 0.147 g cm^-3^.

Parameters related to root hairs were derived from microscopic observations reported on winter wheat and spring barley by Singh Gahoonia *et al.* (1997) and on barley by McElgunn & Harrison (1969). The other anatomical parameters (e.g. related to root exchange surfaces and to the dynamics of endodermis and exodermis) were also calculated from various studies related to wheat or other cereals.

Temperature adjustments were made to growth parameters and/or root age by assuming that the growth rate of roots and their hairs increases linearly with temperature.

#### 6.2 Non-growth parameters

Parameters related to sucrose transport, hexose metabolism, rhizodeposition and rhizodeposit transformation at the root-soil interface were derived from measurements reported in the scientific literature or, in 25% of the cases when no data was available, were assigned a plausible value according to the value of other parameters (see detail in Supporting Table ST).

Only two of them had to be adjusted by trial and errors from our initial guess, in order to obtain plausible simulations and avoid sugar depletion in either the pool of sucrose or the pool of mobile hexose along the roots. Both adjusted parameters were related to the dynamics of sucrose:

- The first adjusted parameter corresponded to the reference rate of hexose consumption by growth, which increases the unloading of sucrose according to the actual rate of hexose consumption by growth within a given root segment (see Eq. S2). After testing several order of magnitudes, we selected 5 10^-13^ mol of hexose per second as the reference consumption rate for which the permeability of phloem should be doubled compared to the reference permeability without growth.
- The second adjusted parameter corresponded to the Michaelis-Menten reference concentration of hexose (*K_loading_*), for which the loading rate of sugars into the phloem is doubled (see Eq. S3). We initially selected a value equivalent to the parameter used for sugar unloading into the root suggested by Barillot *et al.* (2016), i.e. 167 µmol of hexose per gram of dry structural mass, by assuming that the reverse process (i.e. phloem loading) would obey to the same rules. However, this led to an accumulation of sugars in the non-growing root segments and consequently to a high limitation of root elongation because of the low availability of sugars at root apices. We eventually chose 5 µmol of hexose per gram of dry structural mass as K_loading_ for avoiding such situation.

### 7 Complementary results of spring wheat belowground C dynamics, as simulated by *RhizoDep*

#### 7.1 Evolution of the mass flow rate of rhizodeposition in scenario Sc1

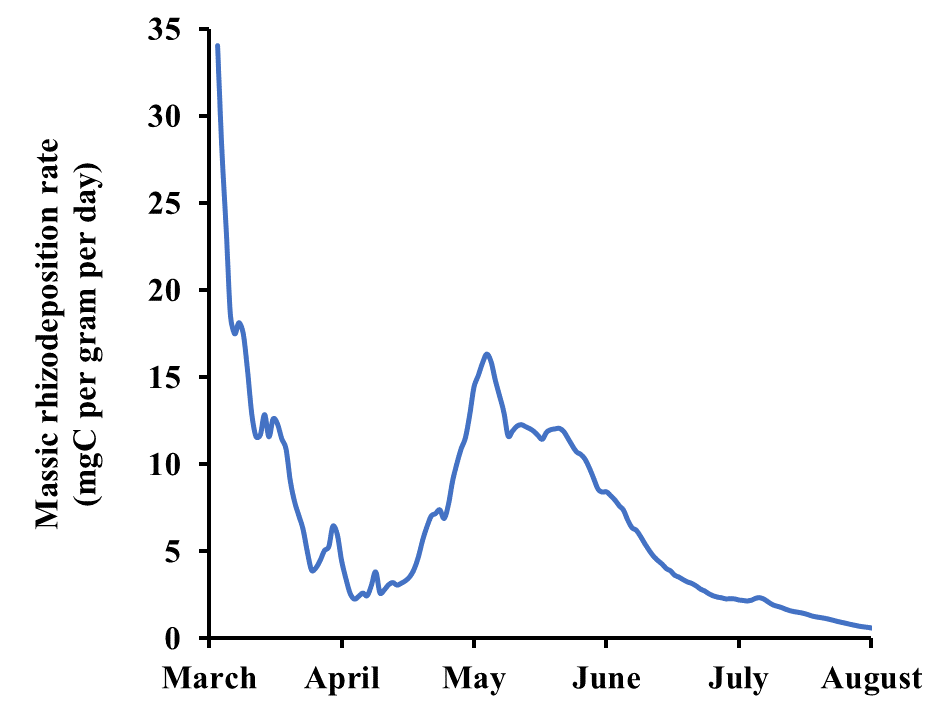

***Fig. S5: Evolution of the mass rate of rhizodeposition (mgC per gram of roots per day) over the growth of spring wheat from March to August 1989****, as simulated by RhizoDep in scenario Sc1 using the flow of belowground C allocation calculated from the case study of Swinnen et al. (1994c) and the soil temperature evolution measured on the same site in 1990.*

#### 7.2 Spatial distribution of rhizodeposition and other variables in scenario Sc1

A. **Linear scale**

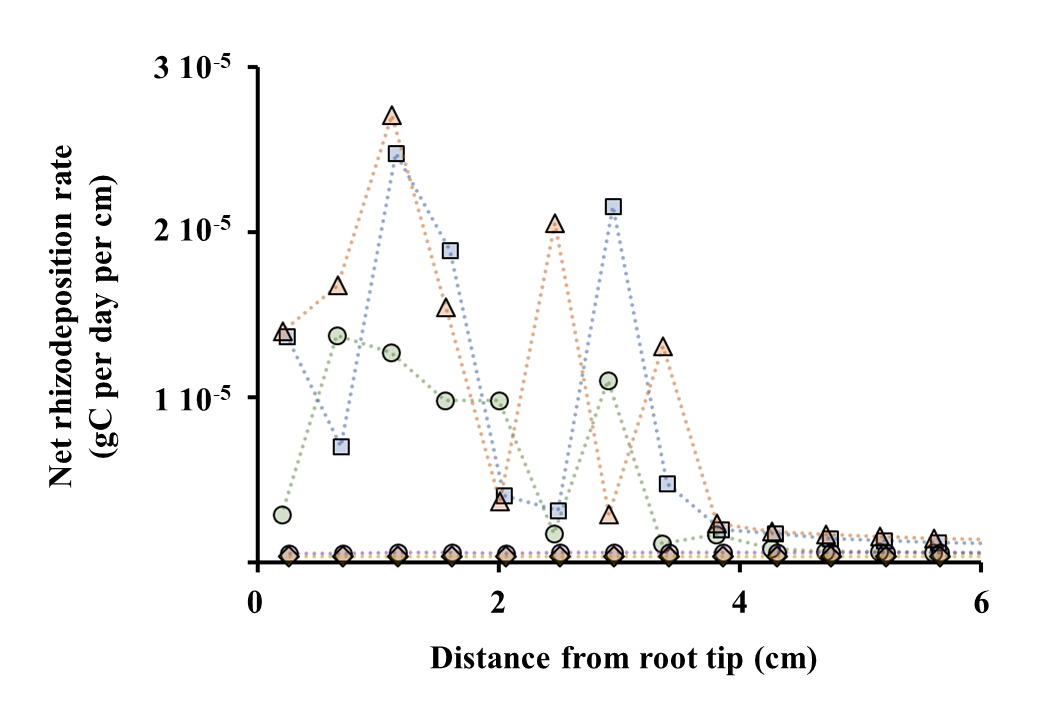

B. **Logarithmic scale**

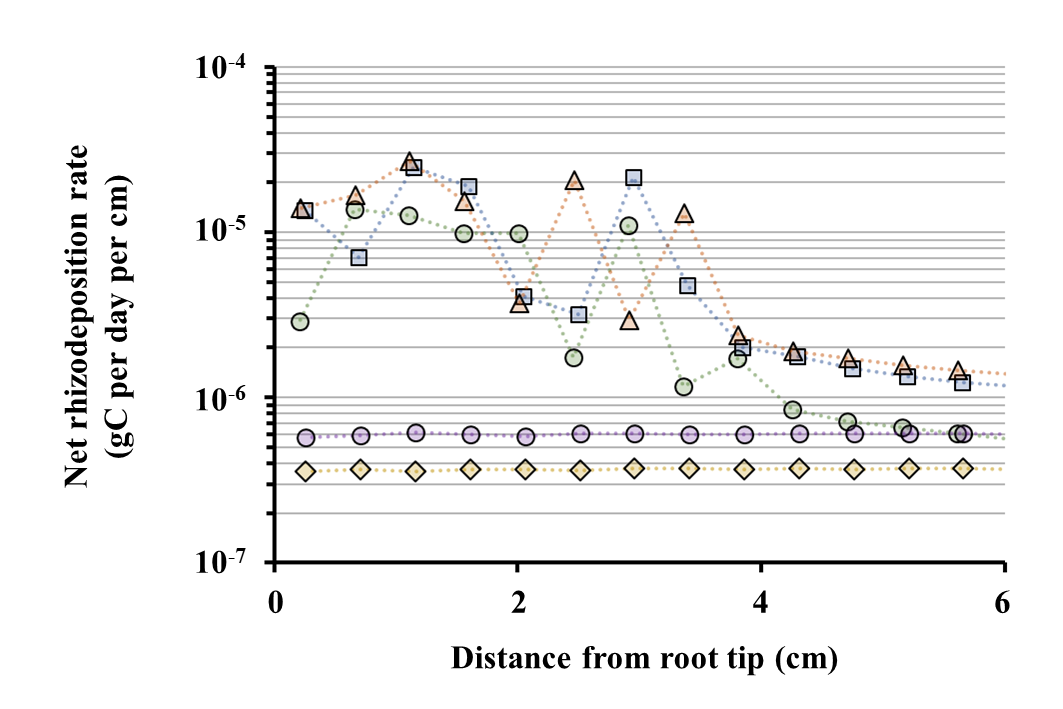

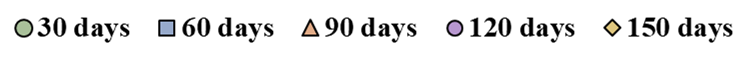

***Fig. S6: Distribution of lineic rhizodeposition rate (gC per cm per day) along the first seminal axis after 1, 2, 3, 4 and 5 months, as simulated by RhizoDep in scenario Sc1, using a linear (A) or log- (B) scale.*** *Note that the first seminal axis stopped elongating 104 days after germination.*

A. **Linear scale**

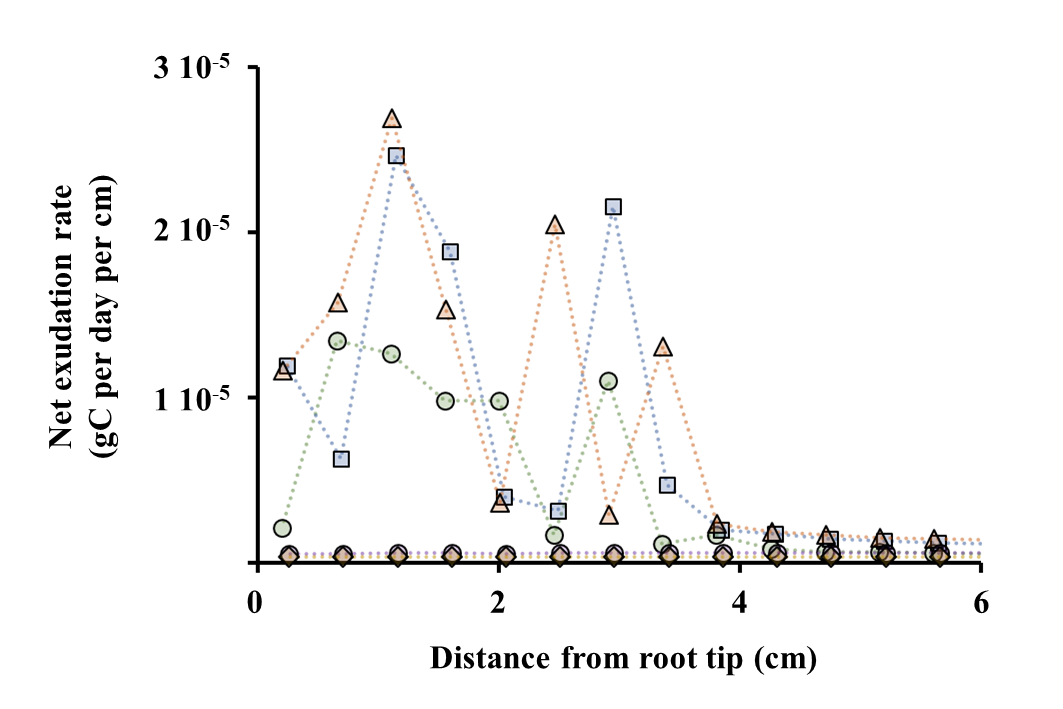

B. **Logarithmic scale**

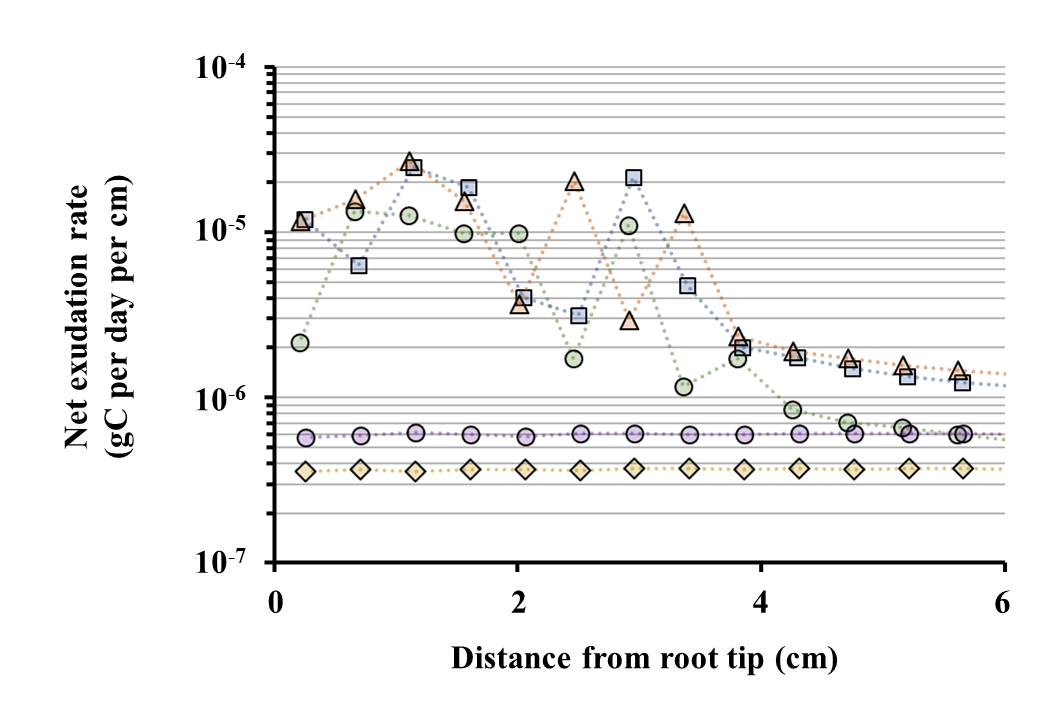

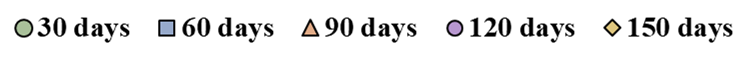

***Fig. S7: Distribution of lineic exudation rate (gC per cm per day) along the first seminal axis after 1, 2, 3, 4 and 5 months, as simulated by RhizoDep in scenario Sc1, using a linear (A) or log- (B) scale.*** *Note that the first seminal axis stopped elongating 104 days after germination.*

A. **Linear scale**

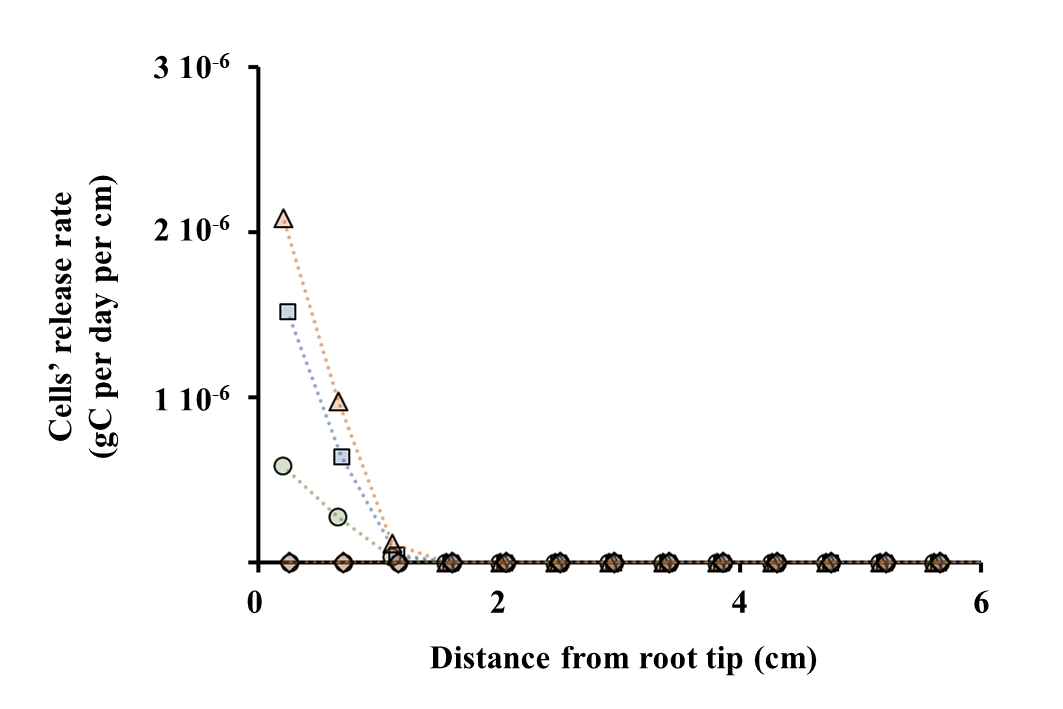

B. **Logarithmic scale**

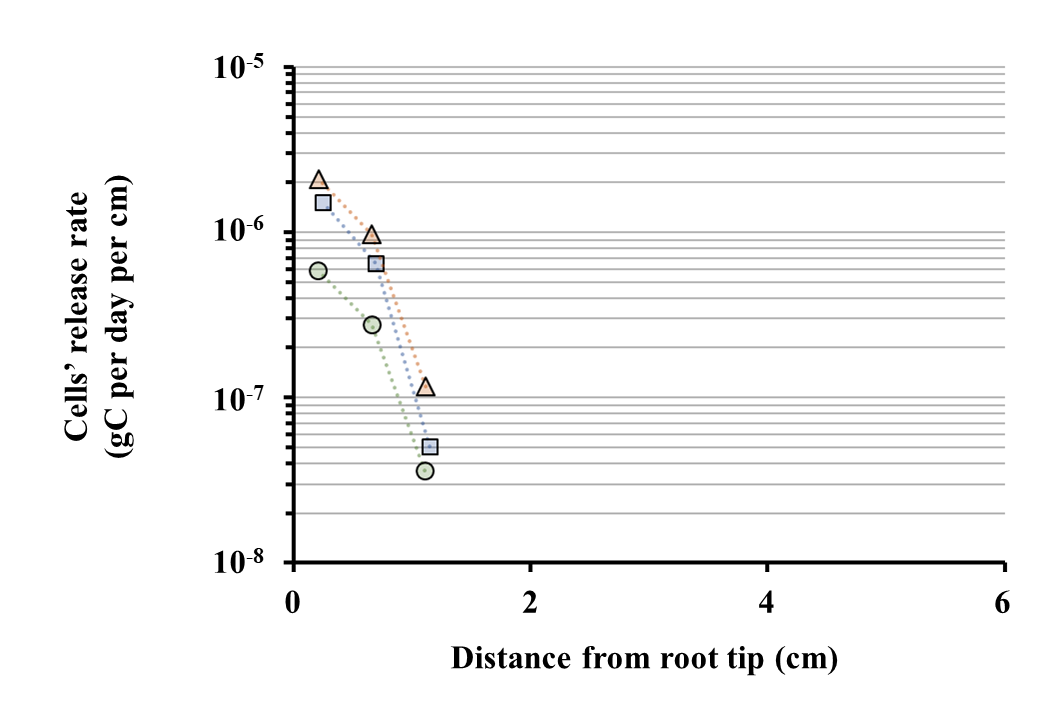

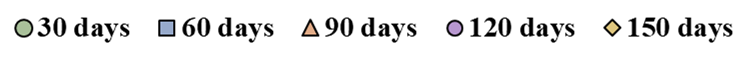

***Fig. S8: Distribution of the lineic rate of cap cells’ release (gC per cm per day) along the first seminal axis after 1, 2, 3, 4 and 5 months, as simulated by RhizoDep in scenario Sc1, using a linear (A) or log- (B) scale.*** *Note that the first seminal axis stopped elongating 104 days after germination. Note that nil values cannot be represented in the lower graph.*

A. **Linear scale**

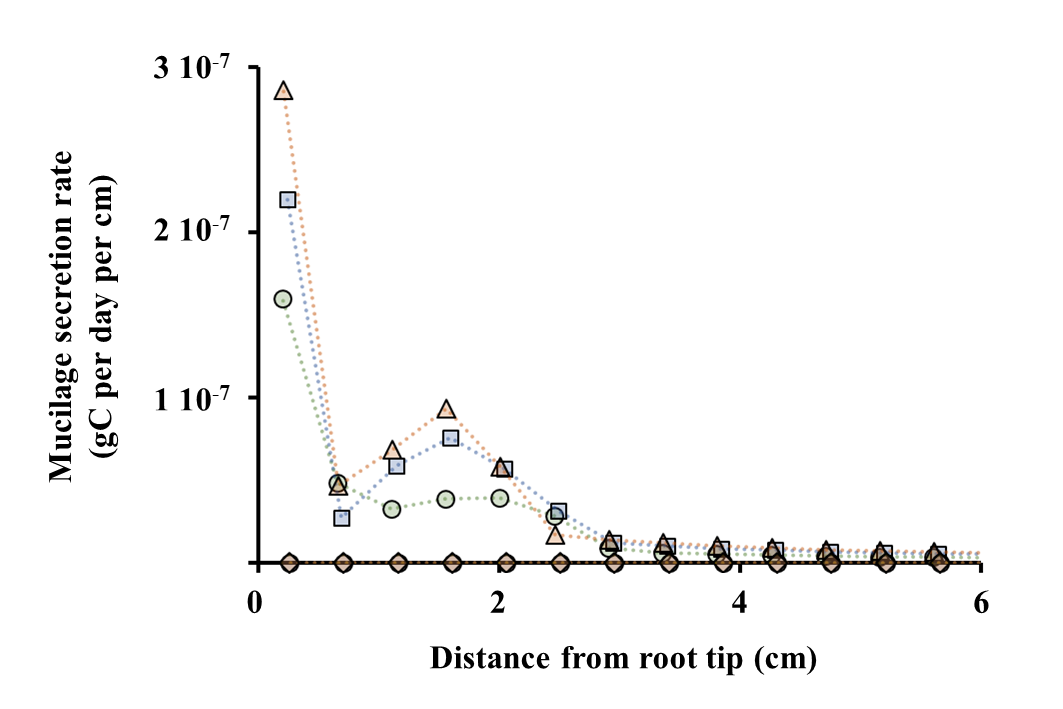

B. **Logarithmic scale**

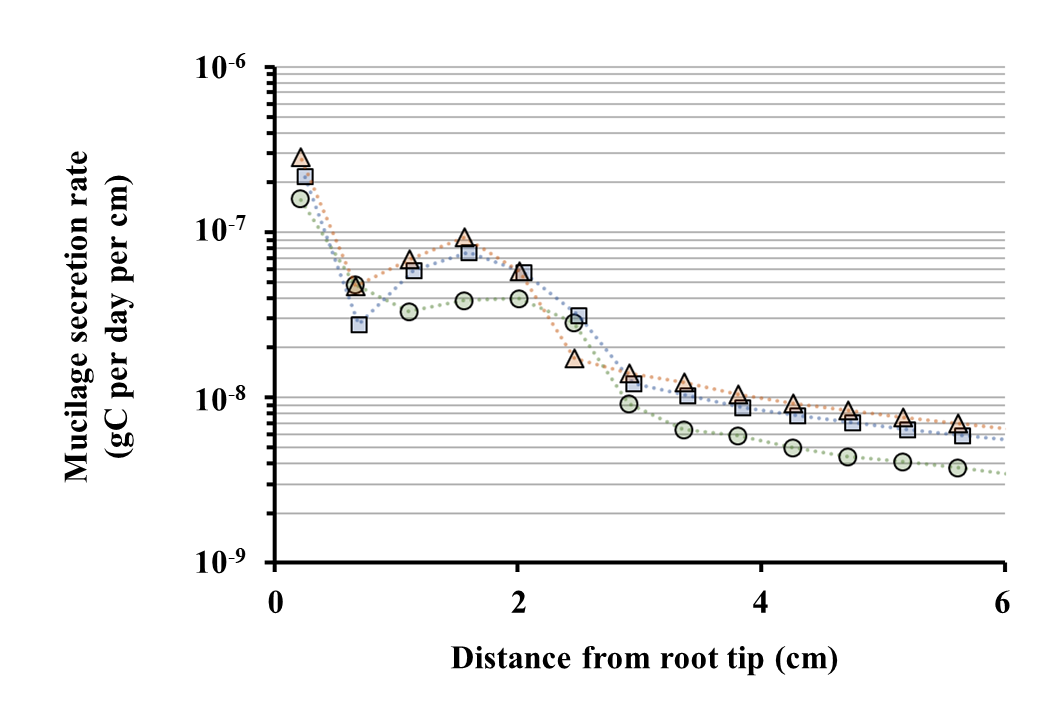

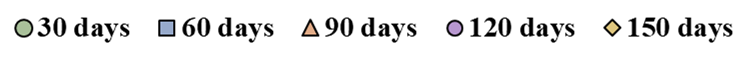

***Fig. S9: Distribution of the lineic rate of mucilage secretion (gC per cm per day) along the first seminal axis after 1, 2, 3, 4 and 5 months, as simulated by RhizoDep in scenario Sc1, using a linear (A) or log- (B) scale.*** *Note that the first seminal axis stopped elongating 104 days after germination.*

A. **Linear scale**

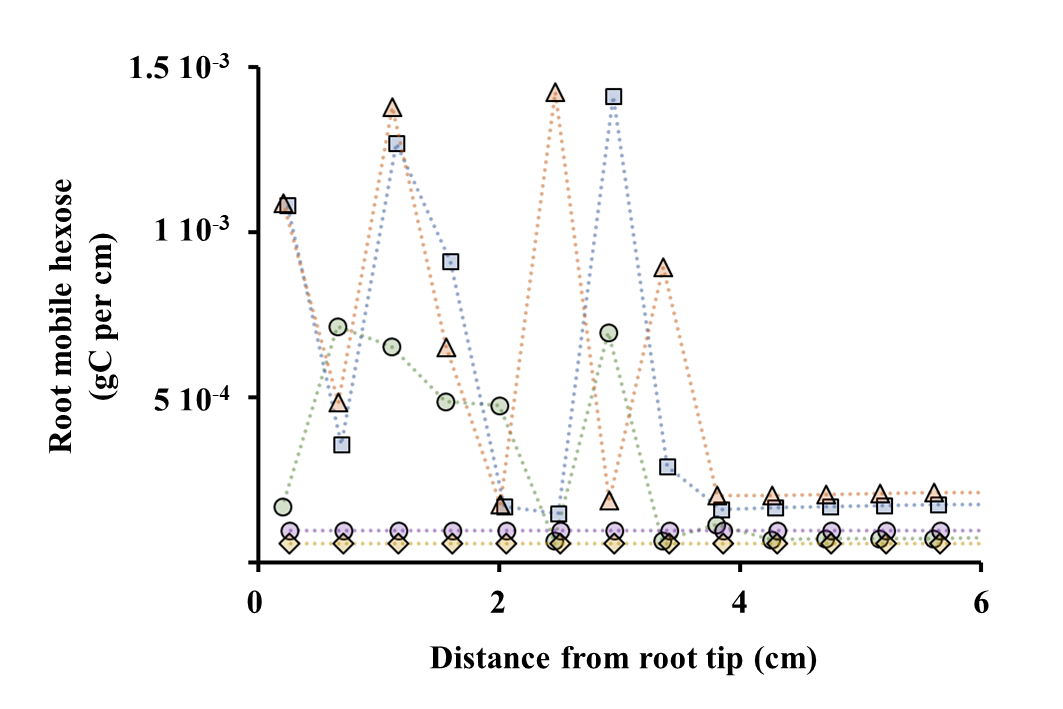

B. **Logarithmic scale**

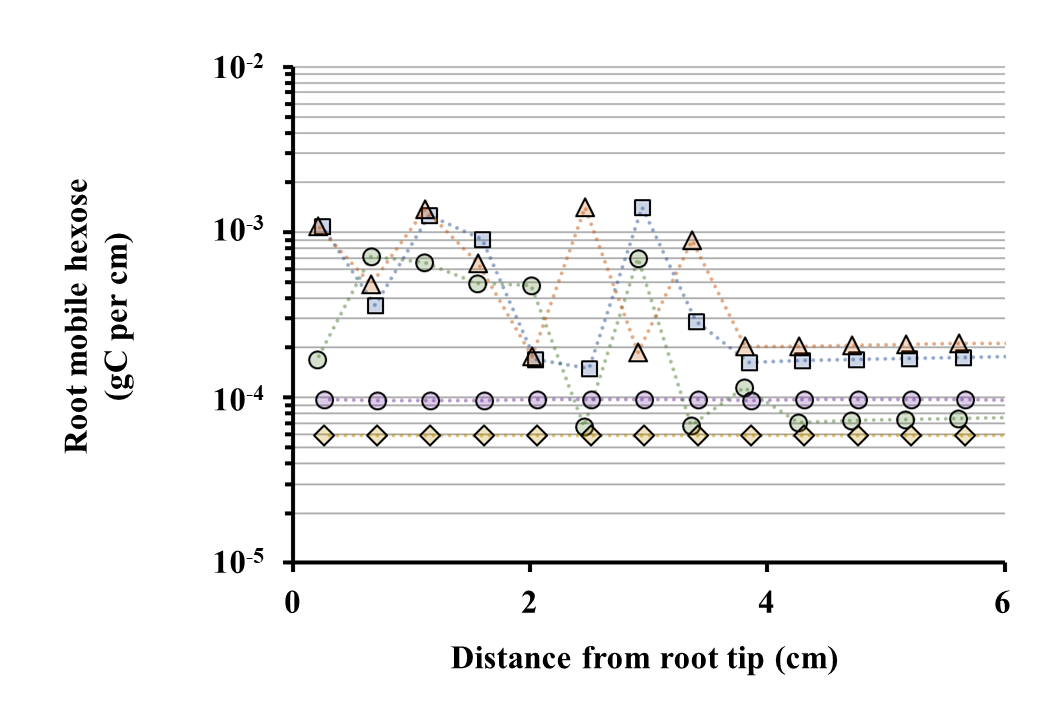

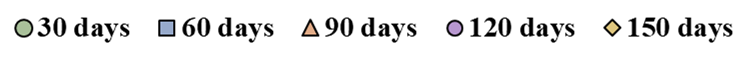

***Fig. S10: Distribution of the root mobile hexose (gC per cm) along the first seminal axis after 1, 2, 3, 4 and 5 months, as simulated by RhizoDep in scenario Sc1, using a linear (A) or log- (B) scale.*** *Note that the first seminal axis stopped elongating 104 days after germination.*

A. **Linear scale**

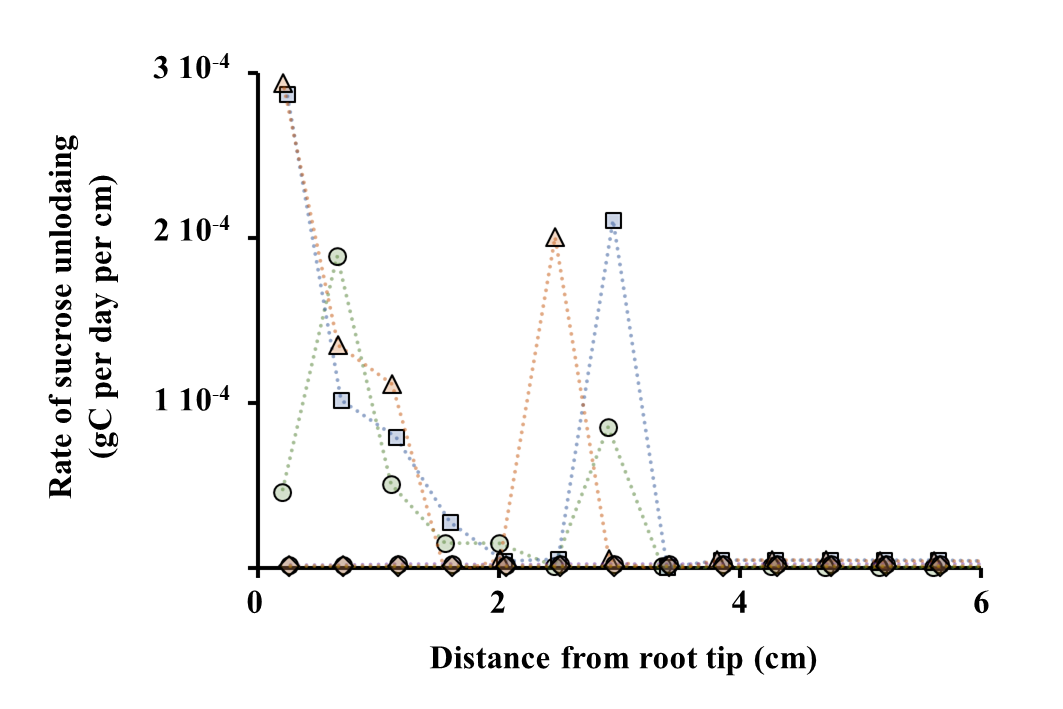

B. **Logarithmic scale**

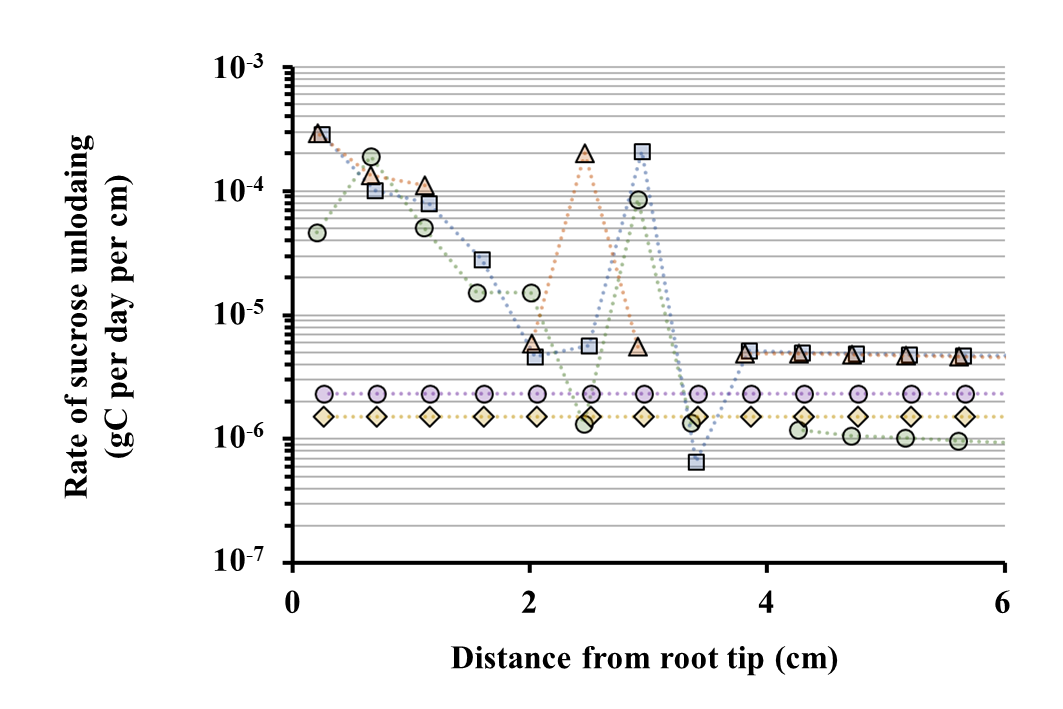

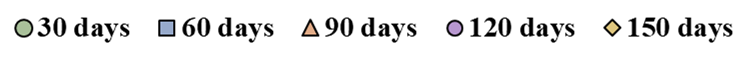

***Fig. S11: Distribution of the rate of net sucrose unloading (gC per day per cm) along the first seminal axis after 1, 2, 3, 4 and 5 months, as simulated by RhizoDep in scenario Sc1, using a linear (A) or log- (B) scale.*** *Note that the first seminal axis stopped elongating 104 days after germination.*

A. **Linear scale**

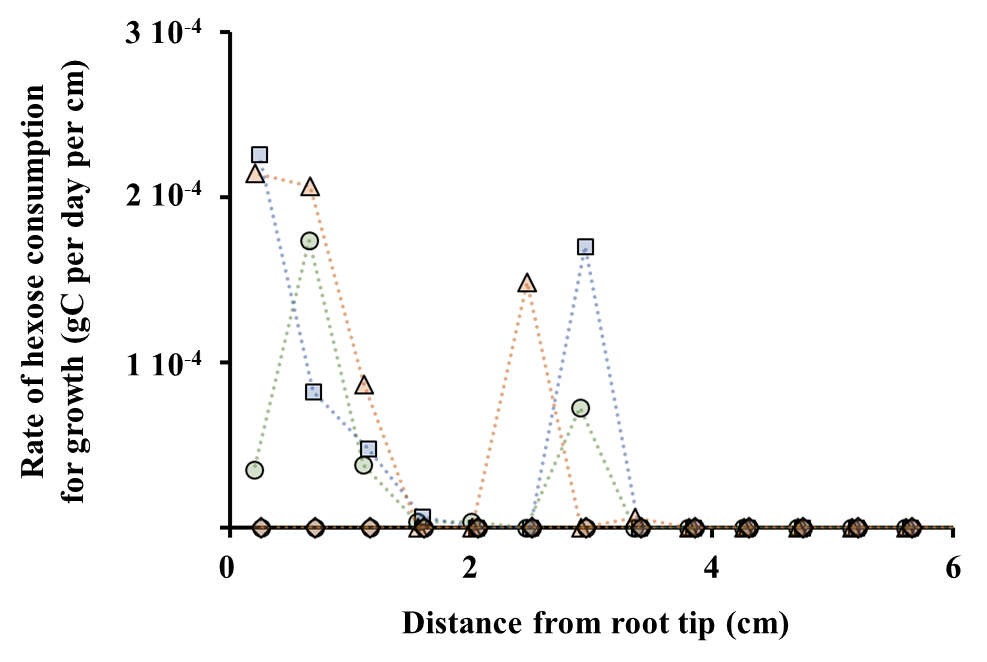

B. **Logarithmic scale**

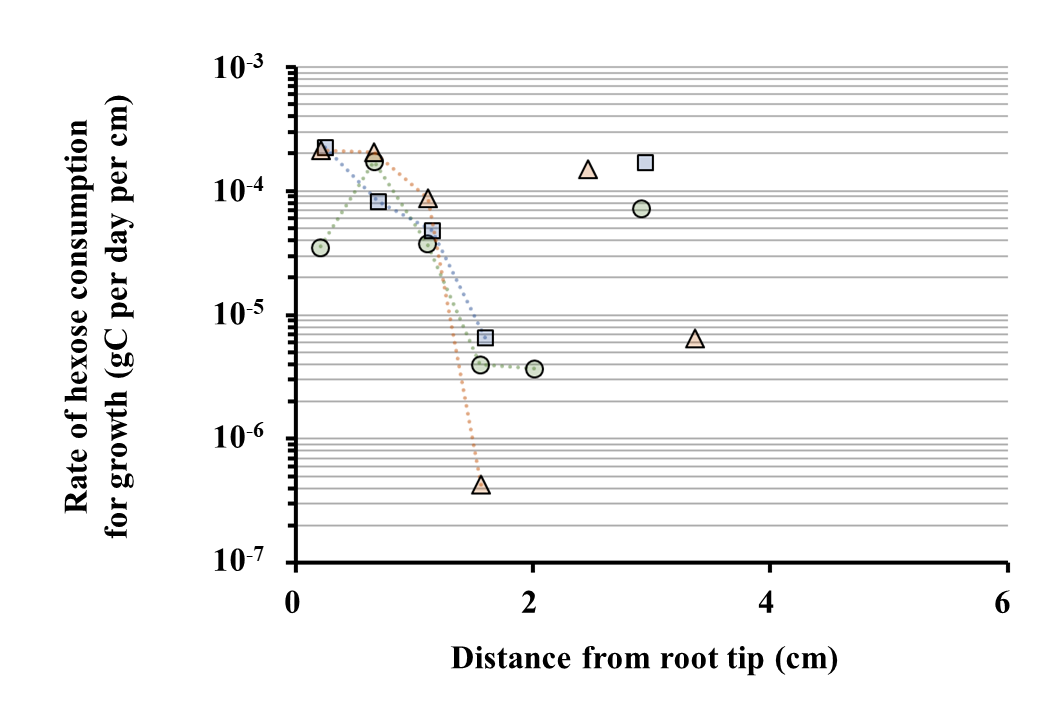

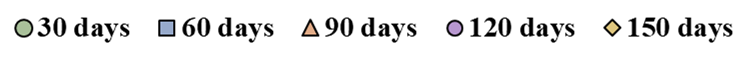

***Fig. S12: Distribution of the rate of hexose consumption for sustaining growth (gC per day per cm) along the first seminal axis after 1, 2, 3, 4 and 5 months, as simulated by RhizoDep in scenario Sc1, using a linear (A) or log- (B) scale.*** *Note that the first seminal axis stopped elongating 104 days after germination. Note that nil values cannot be represented in the lower graph.*

A. **Root-soil exchange surface**

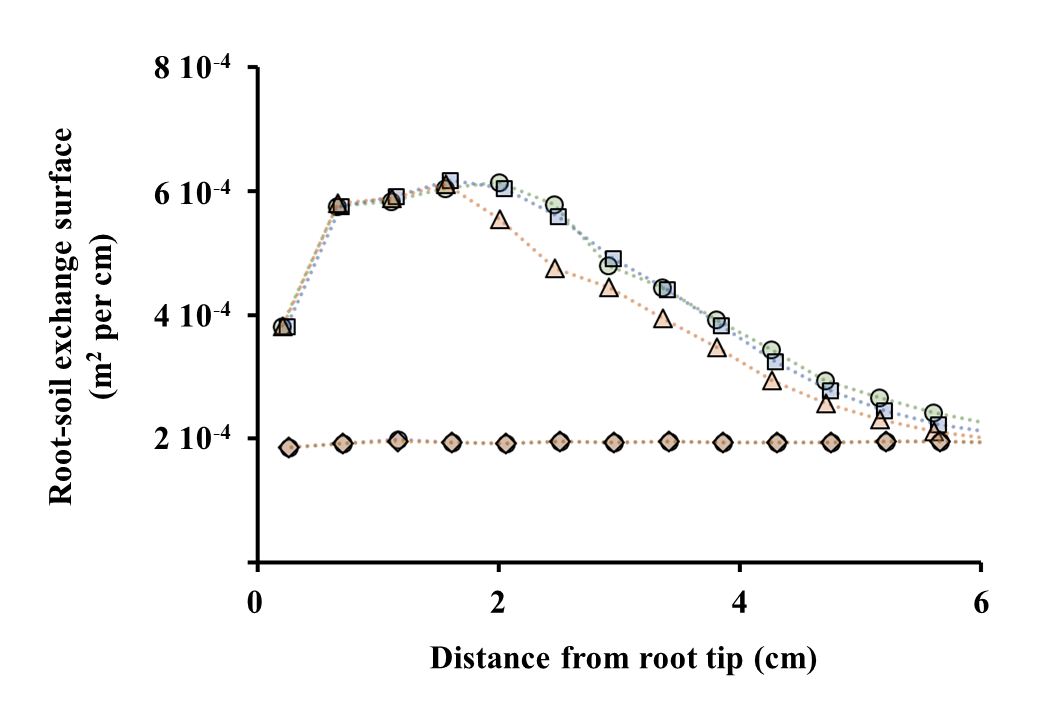

B. **External surface of living root hairs**

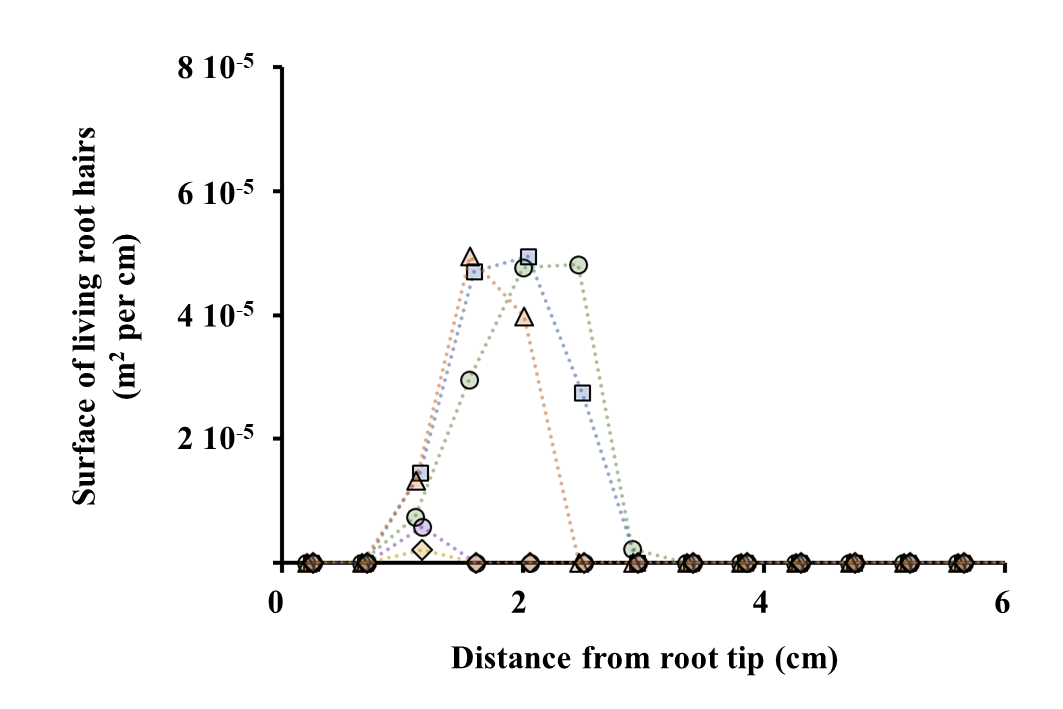

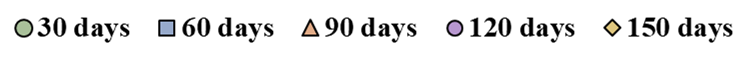

***Fig. S13: Distribution of the area of root-soil exchange surface (A) and of the external surface of living root hairs (B) (m^2^ per cm) along the first seminal axis after 1, 2, 3, 4 and 5 months, as simulated by RhizoDep in scenario Sc1.*** *Note that the first seminal axis stopped elongating 104 days after germination.*

A. **Rhizodeposition as a function of root mobile hexose**

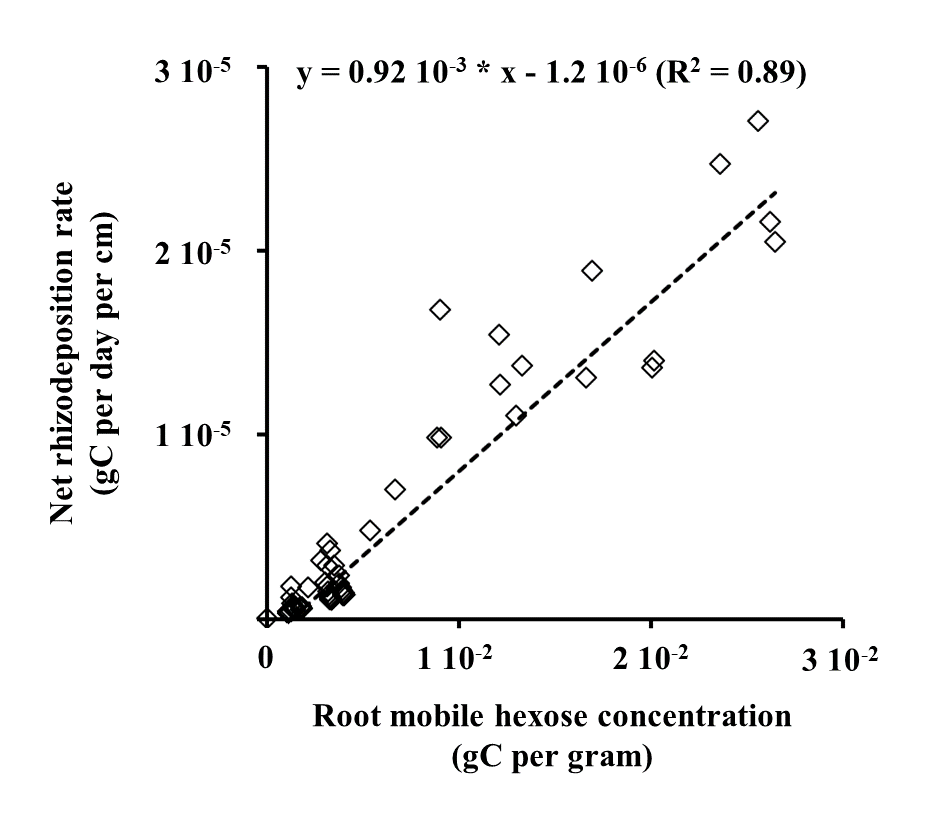

B. **Rhizodeposition as a function of root-soil exchange surface**

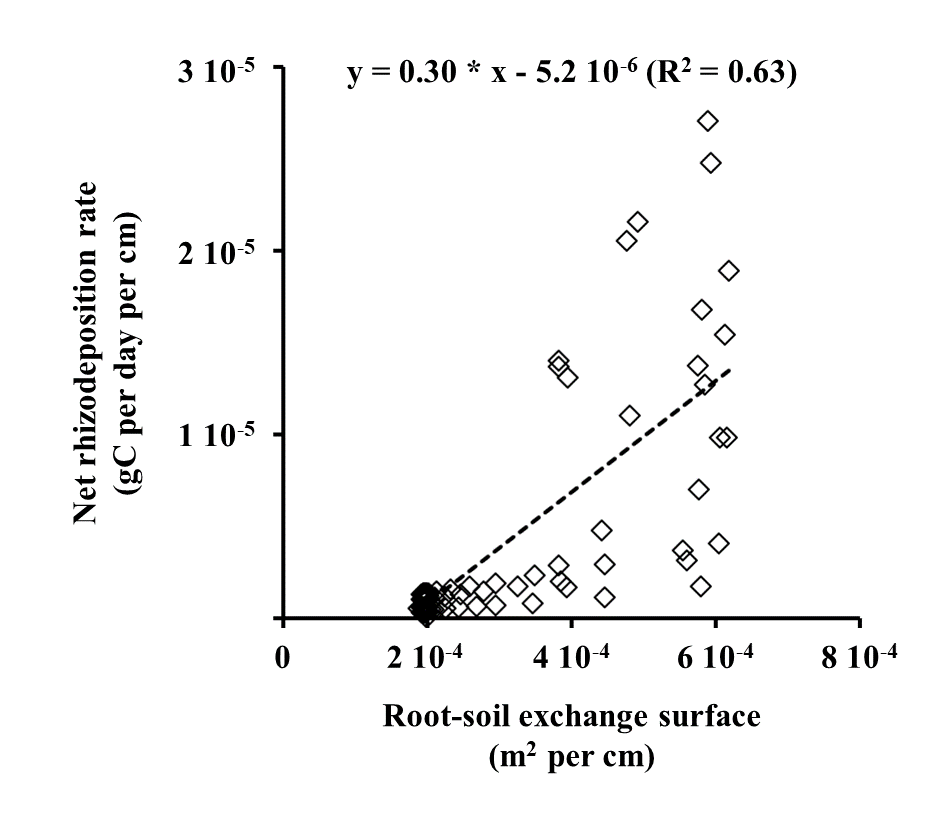

***Fig. S14: Distribution of the lineic rate of rhizodeposition with root mobile hexose (A) or root exchange surface (B) along the first seminal axis after 1, 2, 3, 4 and 5 months, as simulated by RhizoDep in scenario Sc1.*** *Note that the first seminal axis stopped growing 104 days after germination.*

A. **Rhizodeposition as a function of root mobile hexose**

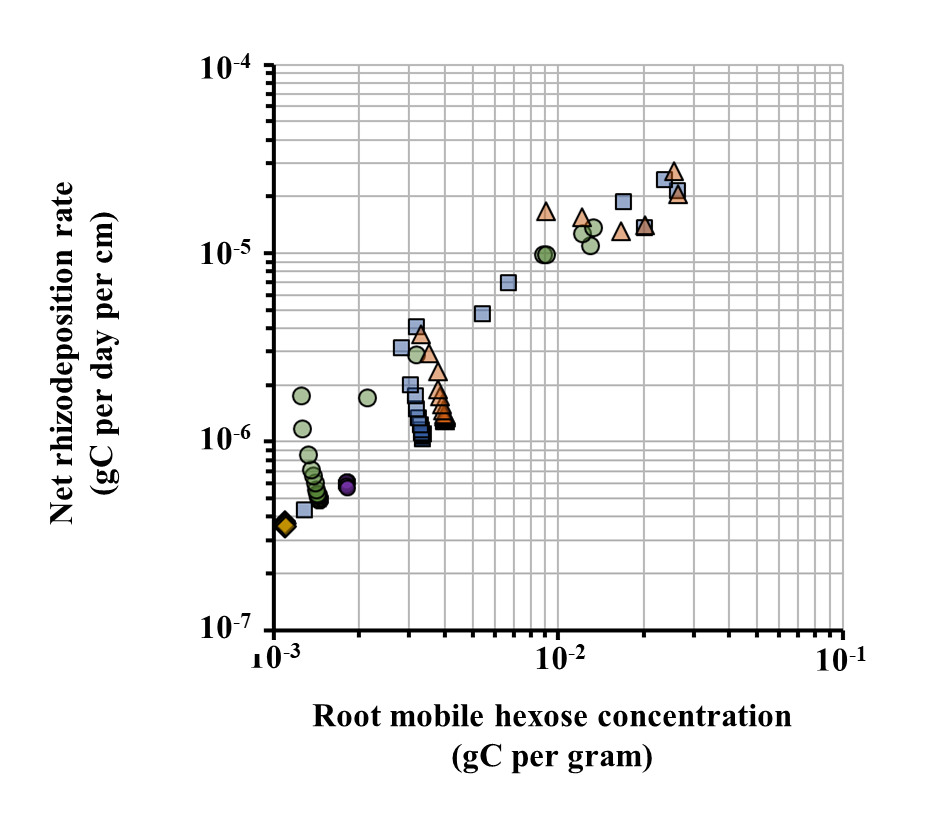

B. **Rhizodeposition as a function of root-soil exchange surface**

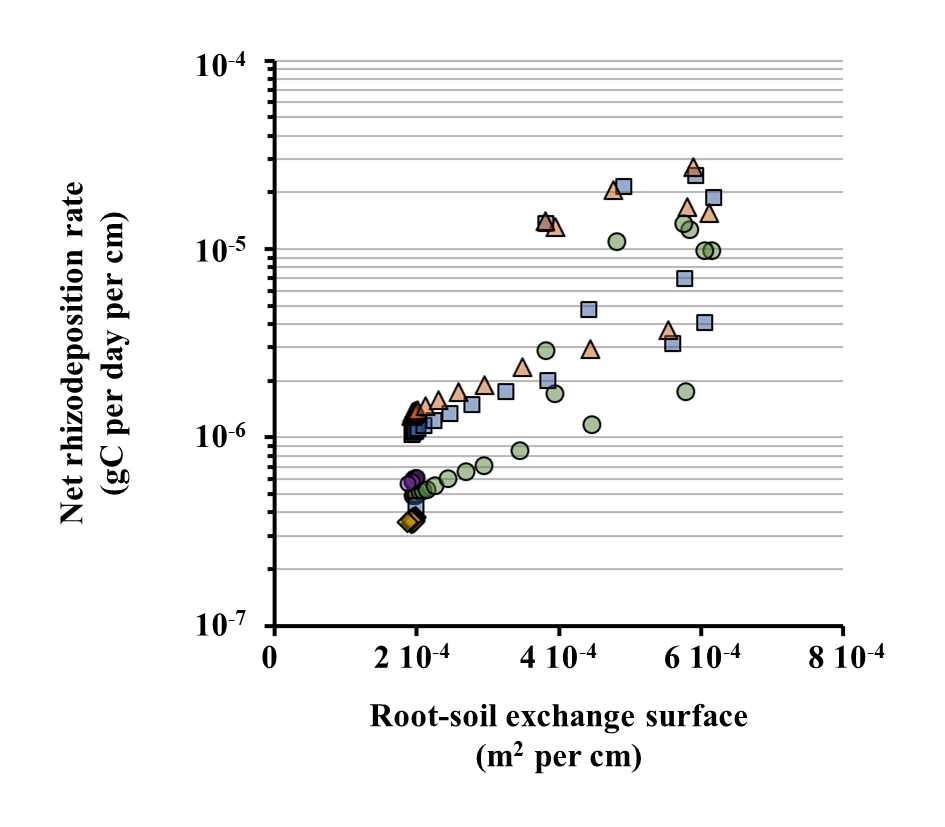

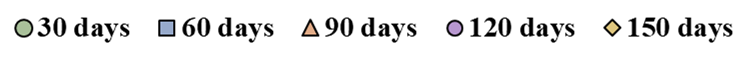

***Fig. S15: Distribution of the lineic rate of rhizodeposition with root mobile hexose (A) or root exchange surface (B) along the first seminal axis after 1, 2, 3, 4 and 5 months, as simulated by RhizoDep in scenario Sc1.*** *Note that the first seminal axis stopped growing 104 days after germination, and that the rhizodeposition rate and the hexose concentration are displayed in log scale.*

**Rhizodeposition as a function of root-soil exchange surface**

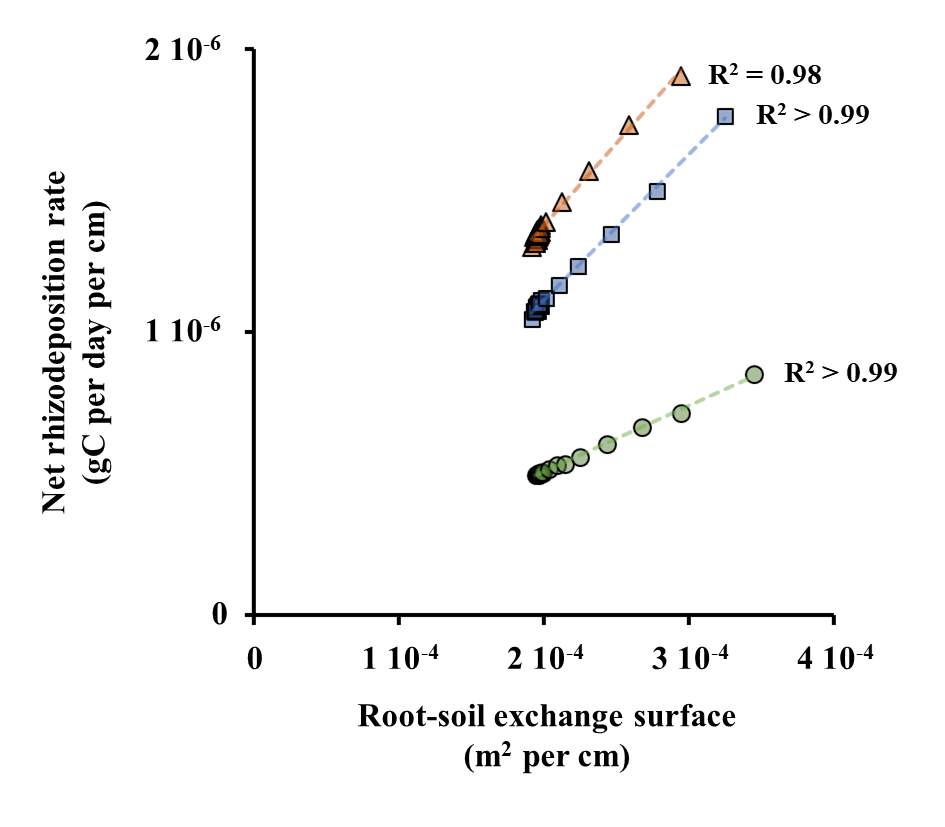

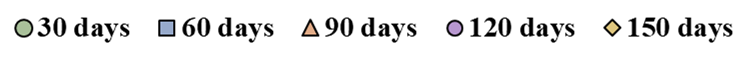

***Fig. S16: Distribution of the lineic rate of rhizodeposition with root exchange surface along the mature part (distance from tip > 4 cm) of the first seminal axis after 1, 2, and 3 months, as simulated by RhizoDep in scenario Sc1.*** *The first segment at the base of the root system was excluded from this graph, as its rhizodeposition rate was significantly lower and did not correspond to the linear relationship observed here with the exchange surface.*

#### 7.3 Relative contribution of different parts of the root system

A. **Distribution of root length among different root classes**

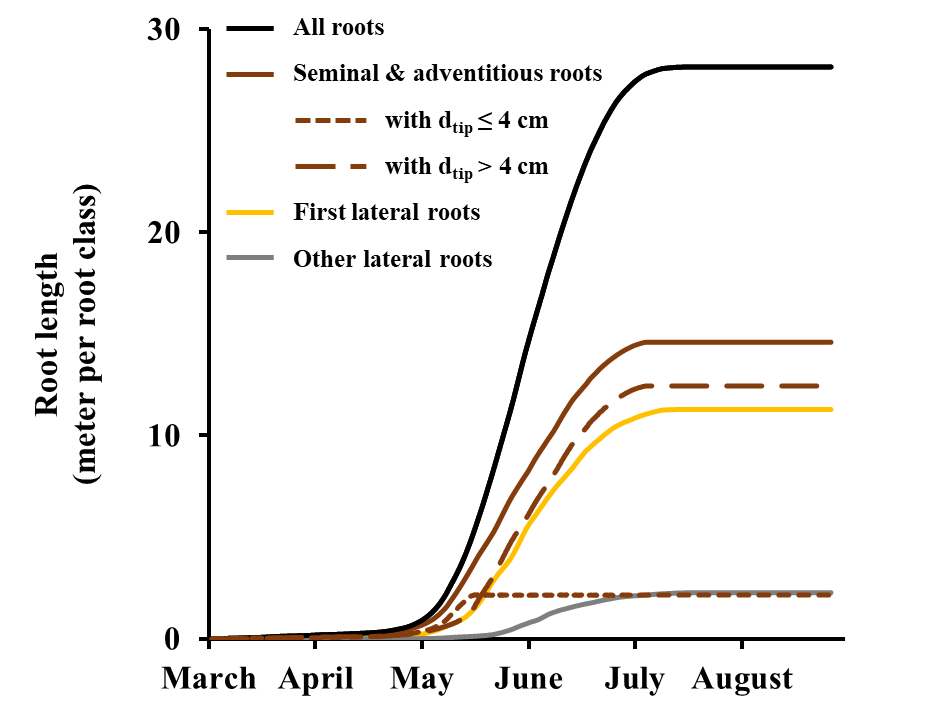

B. **Distribution of rhizodeposited C among different root classes**

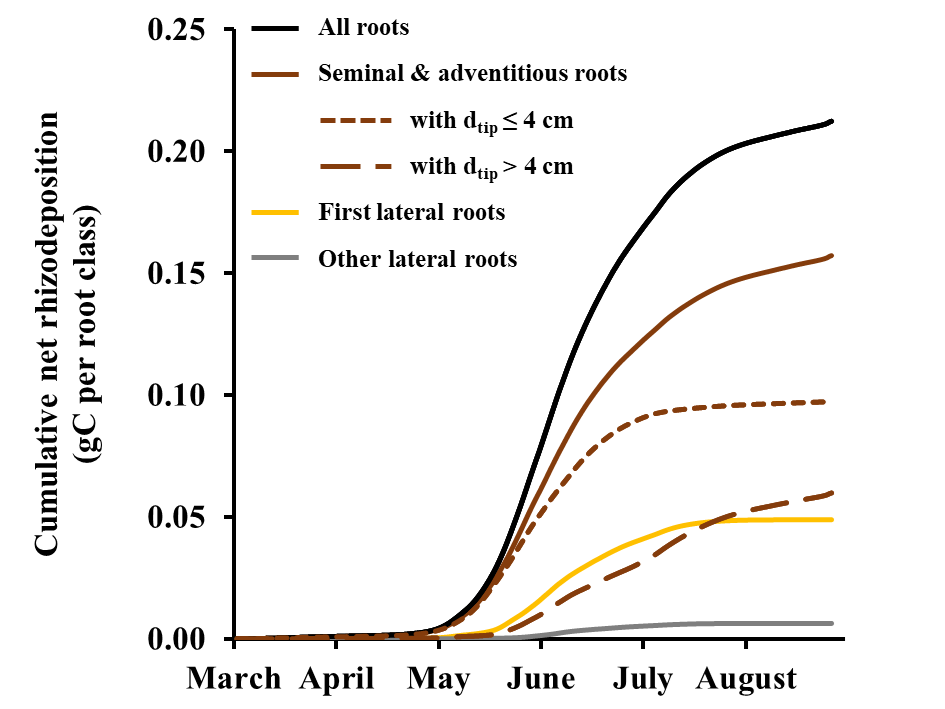

***Fig. S17: Distribution of root length (A) and cumulative rhizodeposition (B) among different classes of roots over the growth of spring wheat from March to August 1989****, as simulated by RhizoDep in scenario Sc1 using the flow of belowground C allocation calculated from the case study of Swinnen et al. (1994c) and the soil temperature evolution measured on the same site in 1990. Roots are distinguished according to their root order, and are further segregated, in the case of the seminal and adventitious roots, in two groups depending on the distance from root tip (d_tip_).*

A. **Proportion of total root length in apical and basal segments**

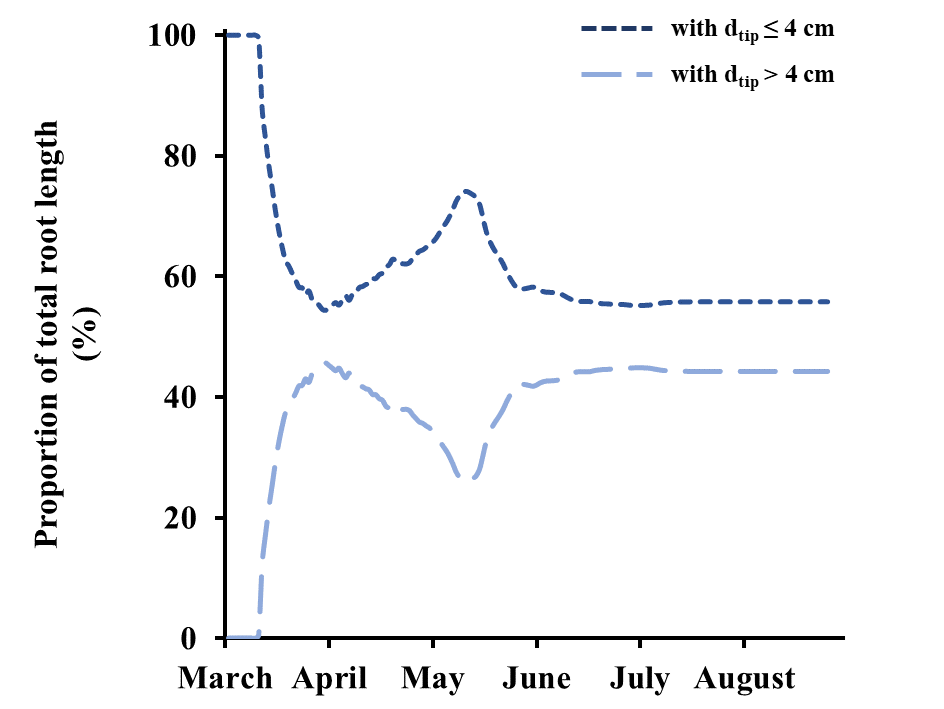

B. **Proportion of rhizodeposited C in apical and basal segments**

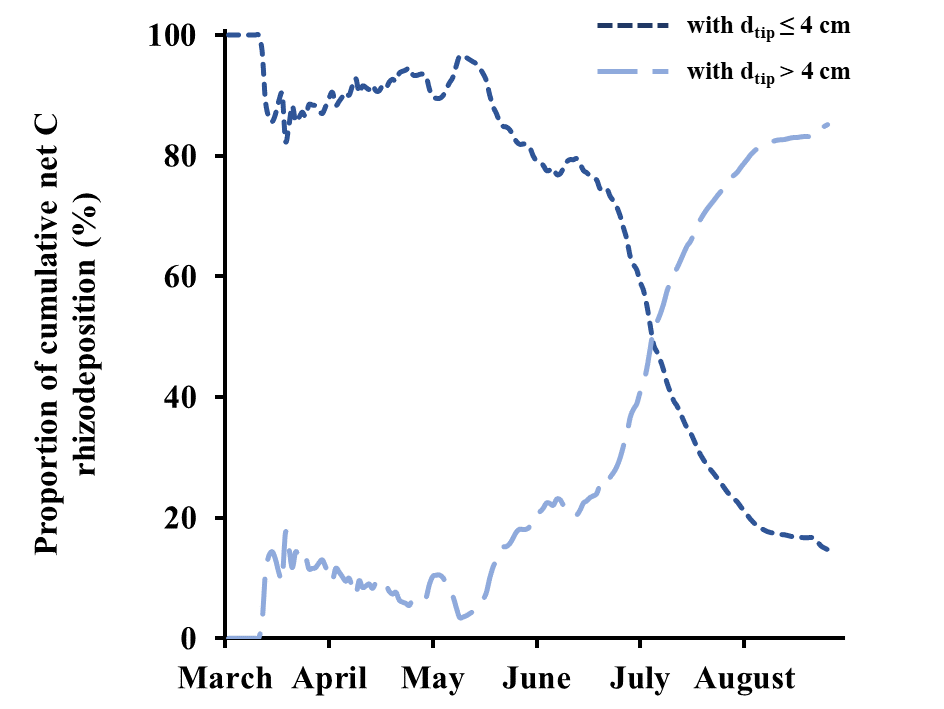

***Fig. S18: Distribution of total root length (A) and cumulative rhizodeposition (B) in apical (distance from root tip d_tip_ ≤ 4 cm) and basal (d_tip_ > 4 cm) root segments over the growth of spring wheat from March to August 1989****, as simulated by RhizoDep in scenario Sc1 using the flow of belowground C allocation calculated from the case study of Swinnen et al. (1994c) and the soil temperature evolution measured on the same site in 1990. Note that dead root segments are not distinguished from living root segments in the calculation of root length.*

***Table S2: Comparison of the proportion of the C rhizodeposited over the first 4 cm of the primary root with the C rhizodeposited by the rest of this root axis and corresponding average lineic net rhizodeposition rate for each root zone****, at 1, 2, 3, 4 and 5 months, as simulated by RhizoDep in scenario Sc1.*

| Time | Root part | Relative proportion  (%) | Net rhizodeposition rate  (µgC per cm per day) |
| --- | --- | --- | --- |
| 30 days | First 4 cm | 86 | 7.12 |
|  | Rest of the root (8 cm) | 14 | 0.56 |
| 60 days | First 4 cm | 68 | 10.57 |
|  | Rest of the root (18 cm) | 32 | 1.13 |
| 90 days | First 4 cm | 51 | 12.23 |
|  | Rest of the root (34 cm) | 49 | 1.37 |
| 120 days | First 4 cm | 8 | 0.58 |
|  | Rest of the root (41 cm) | 92 | 0.60 |
| 150 days | First 4 cm | 8 | 0.35 |
|  | Rest of the root (41 cm) | 92 | 0.37 |

#### 7.4 Main outputs from the different scenarios

The main simulation results from scenario Sc2-Sc7 are summarized below in Fig. S16-S18.

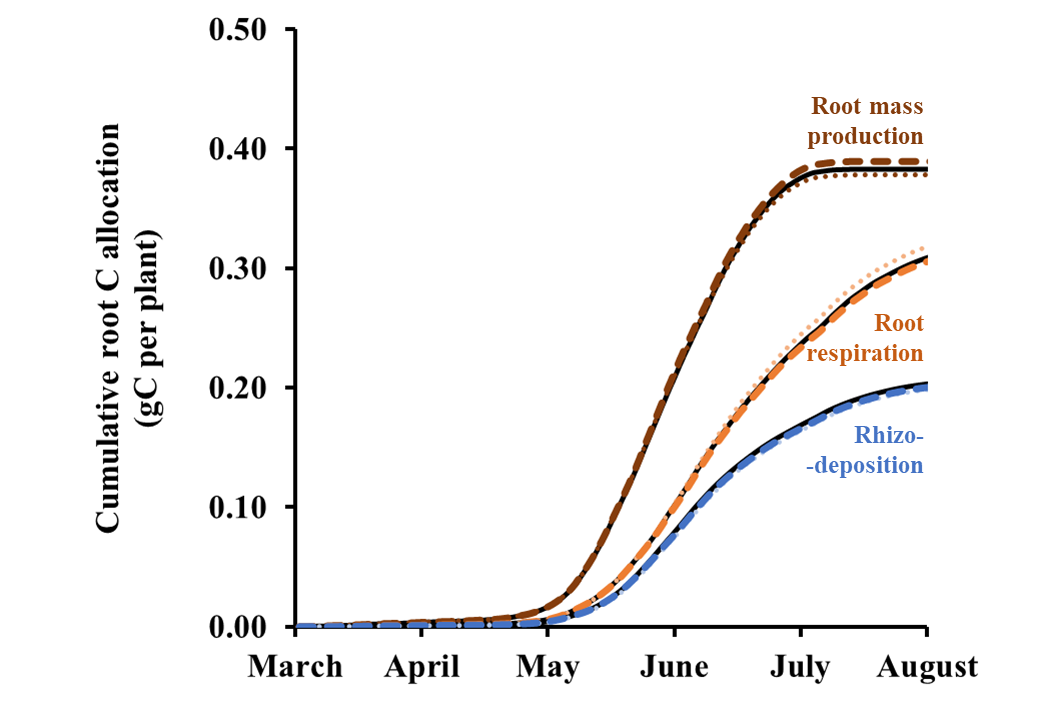

***Fig. S19: Effects of root hairs on*** ***belowground C flows*** *cumulated over the whole growth of spring wheat from March to August 1989, as simulated by RhizoDep. Sc1 (black line): reference scenario reproducing the experiment of Swinnen et al. (1994). Sc2 (dotted line): no root hairs. Sc3 (dashed line): twice more root hairs than in Sc1.*

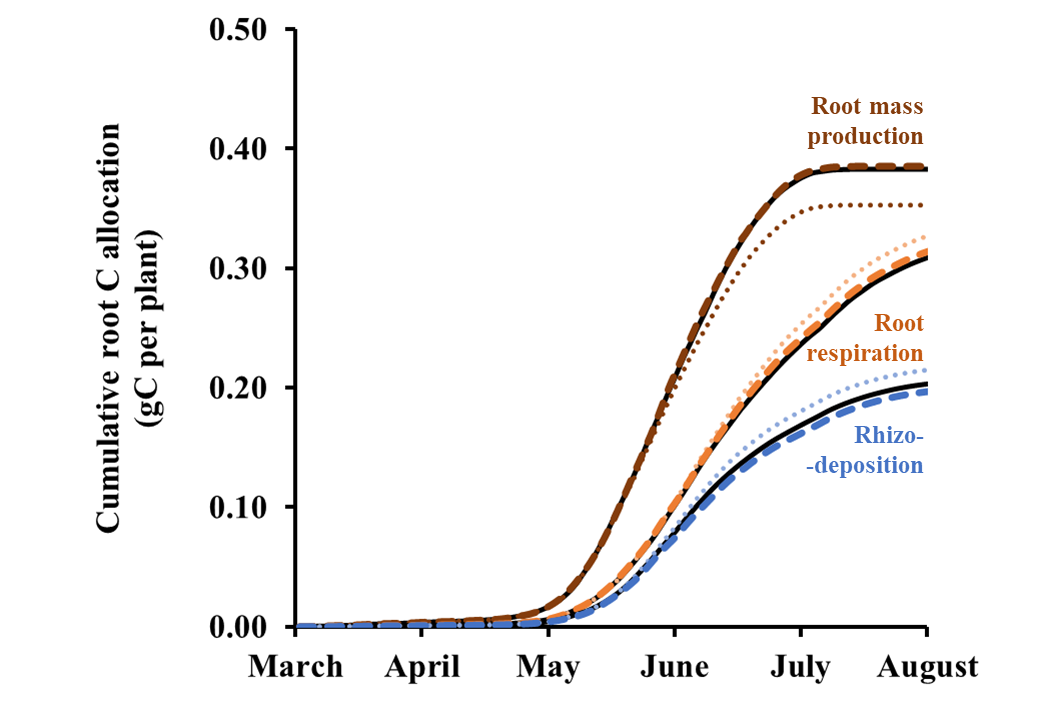

***Fig. S20: Effects of the density of lateral roots on*** ***belowground C flows*** *cumulated over the whole growth of spring wheat from March to August 1989, as simulated by RhizoDep. Sc1 (black line): reference scenario reproducing the experiment of Swinnen et al. (1994). Sc4 (dotted line): 50% less lateral roots than in Sc1. Sc5 (dashed line): twice more lateral roots than in Sc1.*

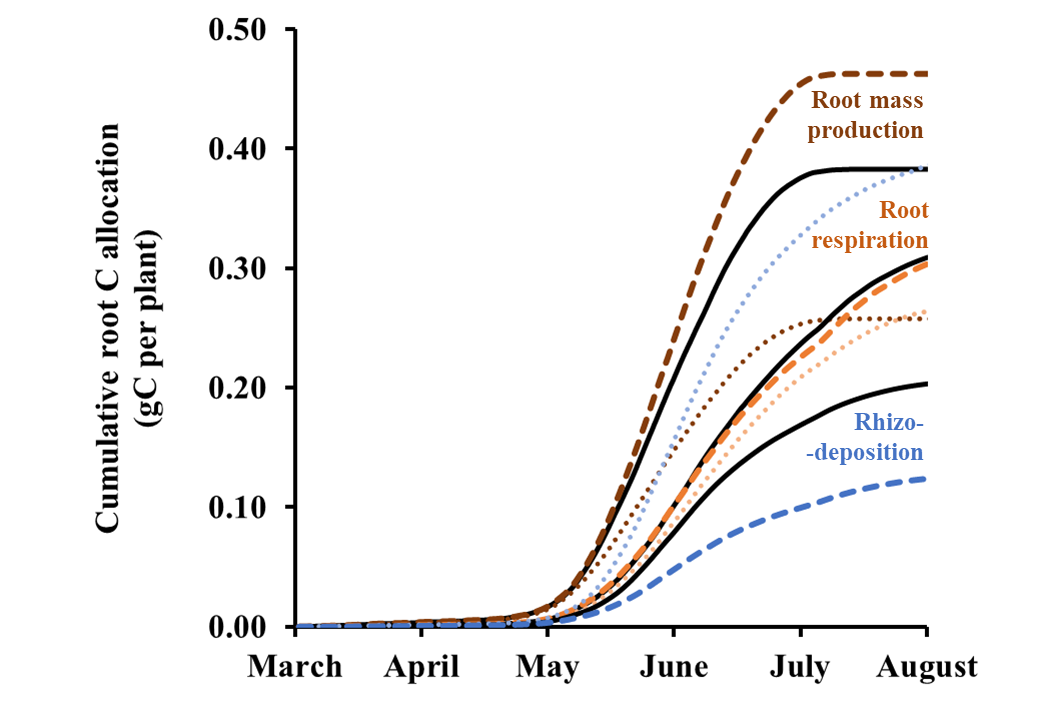

***Fig. S21: Effects of root tissue density on*** ***belowground C flows*** *cumulated over the whole growth of spring wheat from March to August 1989, as simulated by RhizoDep. Sc1 (black line): reference scenario reproducing the experiment of Swinnen et al. (1994). Sc6 (dotted line): 50% lower root tissue density than in Sc1. Sc7 (dashed line): 50% higher root tissue density than in Sc1.*

### 8 Limited assessment of root decay over time

In the absence of precise data, Swinnen *et al.* calculated root mortality as the difference between C inputs and the sum of root biomass production, root respiration and rhizodeposition, thus cumulating all the uncertainties associated to the assessment of these flows (see details in section S4.4). In *RhizoDep*, root mortality was simulated according to *ArchiSimple*’s rule, assigning a predetermined life expectancy to each root axis depending on its apical diameter. However, root growth durations and life expectancies have been shown to be often indeterminate and rather depend on the environment (Cahn *et al.*, 1989; Liljeroth, 1995; Shishkova *et al.*, 2008). As root death occurs after the apical meristem has been deprived from fresh C for too long (Marshall & Waring, 1985; Brouquisse *et al.*, 1991), a refinement of *RhizoDep* could consist in adding a dependency of the root living status to the availability of mobile hexose.

A. **Cumulative root-derived C**

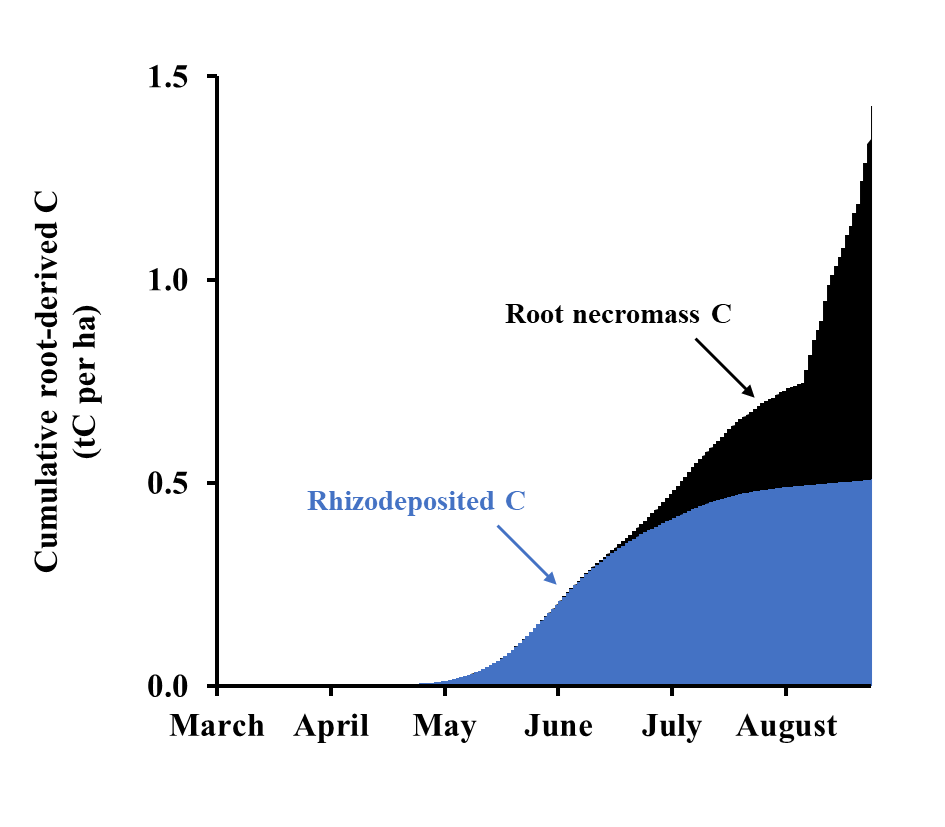

B. **“Bolinder’s ratio”**

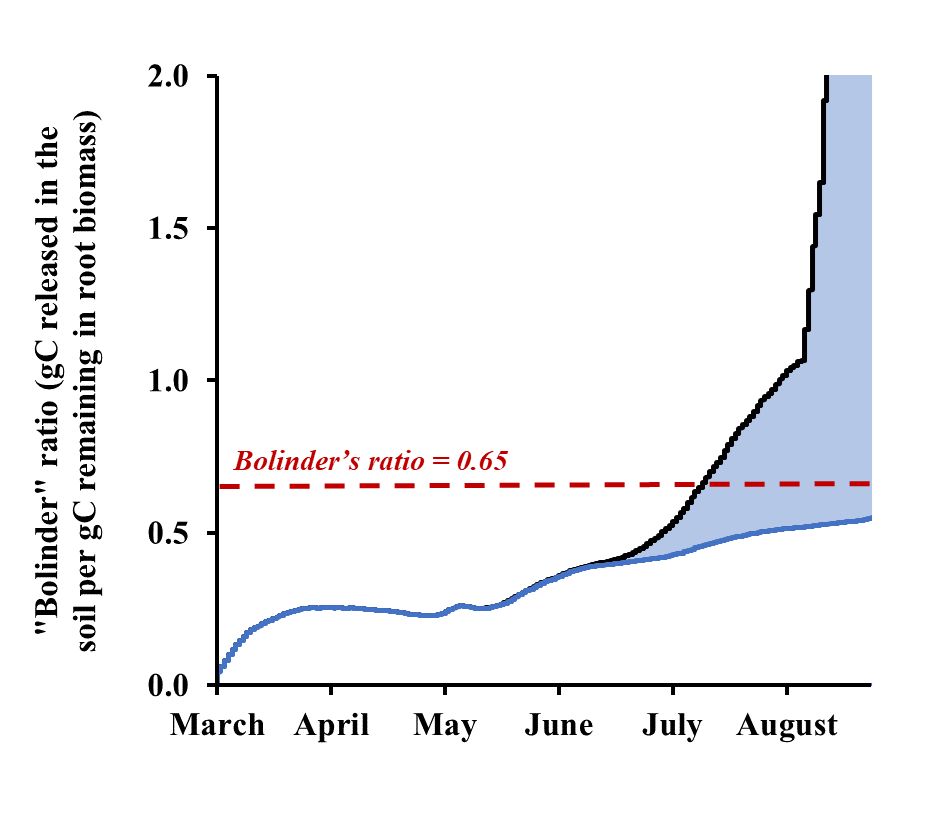

***Fig. S22: Distribution of the cumulative amount of C released in the soil (A) and of the ratio of the amount of rhizodeposited C over the standing root mass at the time of sampling (B) in scenario Sc1.*** *Note that for the graph B, a lower and upper limit of are indicated, depending on whether the actual root necromass is attributed to the rhizodeposited C or to the root biomass at harvest. The lower limits appearing in June corresponds to the ratio of rhizodeposited C over the total mass of root that has been produced (including dead roots), while the upper limit corresponds to the ratio of rhizodeposited C and root necromass C with the mass of living roots only.*

### 9 Distribution of root exchange surfaces along the primary root at 60 days

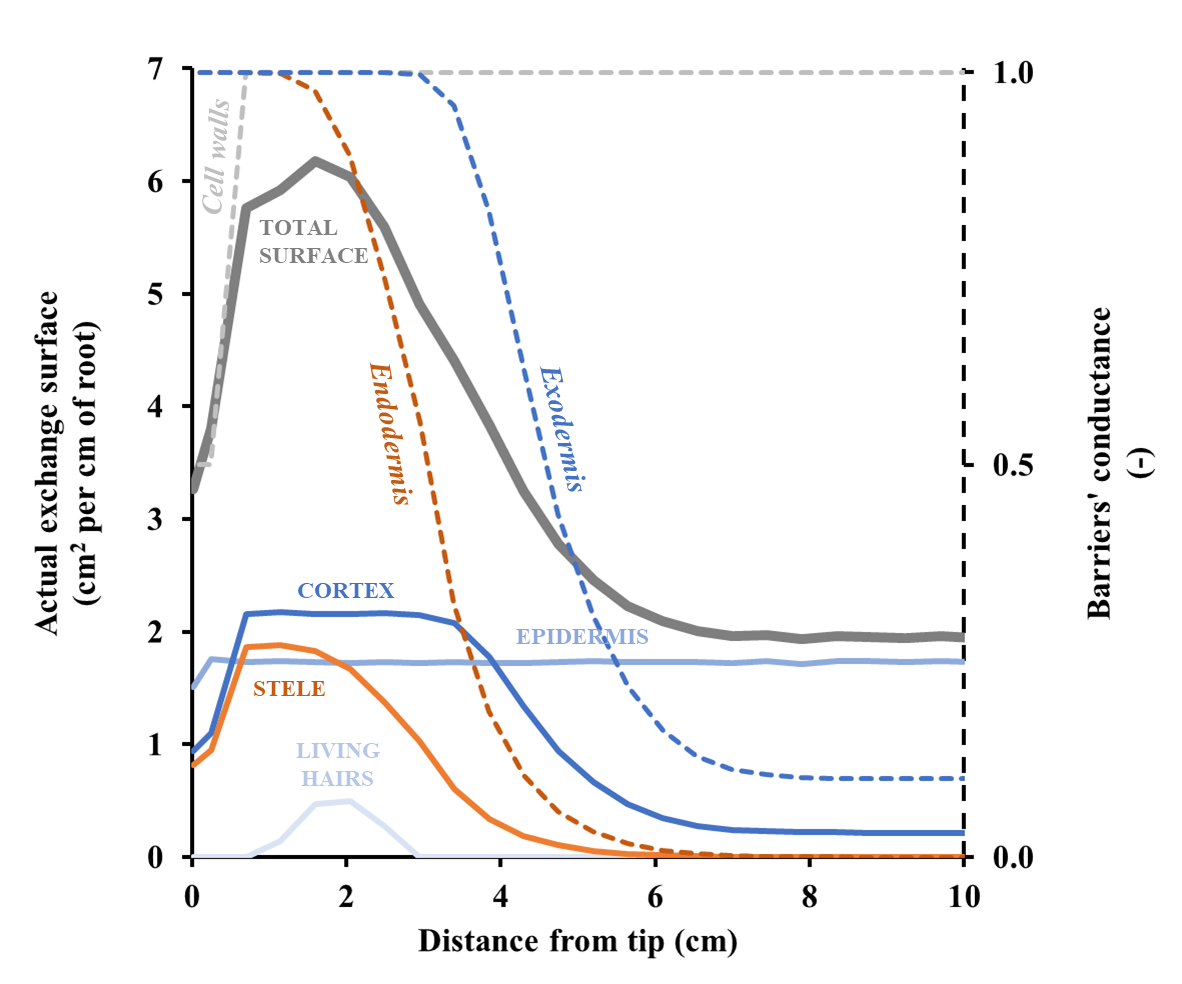

***Fig. S23: Distribution of the different surfaces of exchange between root compartments and the external soil solution (left y-axis) and of the relative conductance (right y-axis) of cell walls, endodermis and exodermis*** ***along the first 10 cm of the primary root 60 days after germination, as simulated by RhizoDep in scenario Sc1.***
